# De novo identification of maximally deregulated subnetworks based on multi-omics data with DeRegNet

**DOI:** 10.1101/2021.05.11.443638

**Authors:** Sebastian Winkler, Ivana Winkler, Mirjam Figaschewski, Thorsten Tiede, Alfred Nordheim, Oliver Kohlbacher

## Abstract

**Background:** With a growing amount of (multi-)omics data being available, the extraction of knowledge from these datasets is still a difficult problem. Classical enrichment-style analyses require predefined pathways or gene sets that are tested for significant deregulation to assess whether the pathway is functionally involved in the biological process under study. *De novo* identification of these pathways can reduce the bias inherent in predefined pathways or gene sets. At the same time, the definition and efficient identification of these pathways *de novo* from large biological networks is a challenging problem.

**Results:** We present a novel algorithm, DeRegNet, for the identification of maximally deregulated subnetworks on directed graphs based on deregulation scores derived from (multi-)omics data. DeRegNet can be interpreted as maximum likelihood estimation given a certain probabilistic model for de-novo subgraph identification. We use fractional integer programming to solve the resulting combinatorial optimization problem. We can show that the approach outperforms related algorithms on simulated data with known ground truths. On a publicly available liver cancer dataset we can show that DeRegNet can identify biologically meaningful subgraphs suitable for patient stratification. DeRegNet is freely available as open-source software.

**Conclusion:** The proposed algorithmic framework and its available implementation can serve as a valuable heuristic hypothesis generation tool contextualizing omics data within biomolecular networks.

## Introduction

Modern high-throughput technologies, in particular massively parallel sequencing [1] and high-resolution mass spectrometry [2], enable omics technologies, i.e. the determination of bioanalytes on the genome-wide scale. Many of of these omics technologies are increasingly being applied in clinical settings and publicly available large-scale data resources such as The Cancer Genome Atlas (TCGA) [3] provide ample opportunity for research. These resources can provide valuable reference data sets in the analysis of molecular profiles of individual patients and patient groups. However, one of the biggest challenges in the analysis of omics data remains functional annotation/interpretation. The interpretation of the experimental read-outs with the goal of understanding the underlying known or unknown biological processes and functions is a vital step in providing personalized, precise, and focused molecular therapies.

One of the most widely used approaches for functional annotation of large omics datasets is Gene Set Enrichment (GSE) [4]. In its most basic form, GSE entails hypergeometric and Fisher test-based approaches to detect the overrepresentation of differentially expressed genes. GSE requires a set of predefined gene sets (typically obtained from pathway databases [5] such as KEGG [6], WikiPathways [7] or Reactome [8]) and a measure of ”deregulation” (e.g., a binary indication of differential gene expression). The goal of the GSE analysis is to identify those gene sets from the collection which show ”high” deregulation. Here, the term ”high” is defined by the method’s specific underlying statistical model. In the simplest case, the method examines if each gene set contains a higher number of differentially expressed genes than would be expected by chance, under the assumption that differentially expressed genes are represented uniformly across all genes. Many adaptations and variations of GSE exist [4, 9].

Classical GSE methods treat pathways as an unstructured collection of genes and do not explicitly account for the extensive biological knowledge encoded in biological networks. Networks as an abstraction for biological knowledge can be represent signaling networks, metabolic networks [10], gene regulatory networks [11], or protein-protein interaction networks [12, 13], and more.

There has been extensive research into the possibility of designing enrichment methods which take into account the topology of the pathways [14, 15, 16]. An example of such approach is the calculation of topology-dependent perturbation scores for each gene [17]. A further aspect usually ignored by GSE methods is the issue of pathway crosstalks. While ’textbook pathways’ have a solid base in biological findings and can provide useful guidance for functional interpretation of omics experiments, molecular and cellular events are often more complicated and involve the direct interaction of molecular entities across predefined pathway boundaries. Correspondingly, a range of methods were proposed which aim to extract ”deregulated” patterns from larger regulatory networks without relying on predefined pathways [18, 19]. These methods are often referred to as *de novo* pathway enrichment (de novo pathway identification, de novo subnetwork/subgraph enrichment/identification/detection) methods, emphasizing that the pathways are defined/extracted from the data itself and are not given as fixed gene sets. Here, we also call algorithms of this flavor deregulated subnetwork/subgraph detection/identification/enrichment methods.

A way to categorize these methods is based on how they handle undirected or directed interaction networks. A lot of biomolecular interactions are directed in nature, e.g. protein A phosphorylates protein B, enzyme A precedes enzyme B in a metabolic pathway in contrast to symmetric interactions such as physical interactions of proteins in protein complexes.

Some methods designed for undirected networks are described in the following studies: [20, 21, 22, 23, 24, 25, 26, 27, 28, 29, 30, 31]. More detailed review of these method is available in [19]. These methods, while achieving similar results on an abstract level, vary greatly in terms of suitable underlying networks, interpretation of outcomes and algorithmic strategies employed. Algorithmic approaches employed include ant colony optimization [31], dynamic programming [27], simulated annealing [20], integer programming [24, 23], Markov random fields [32] or message passing approaches [28].

Also, some methods are tailored to the characteristics of a particular data type. An example are methods attempting to find significantly mutated pathways/networks [33, 34, 35, 36, 37, 38], trying to factor in the pecularities of mutation data in a network context.

While methods which work natively with directed networks are rarer [39, 40, 41, 42, 43, 44], it is instrumental to be able to capture the effects of directed biomolecular interactions in the process of discovering deregulated networks. One particular approach is the one described in [40] which utilized an integer programming approach in order to find deregulated subnetworks. It uncovers deregulated subnetworks downstream or upstream of a so called root node where the latter can be fixed *a priori* or determined by the algorithm itself.

In this paper, we present an algorithm for de novo subnetwork identification which can conceptually be characterized as a mixture of the approach presented by [40] and the price-collecting Steiner tree methods proposed in [45, 46, 47, 48]. Our method natively handles directed interaction networks and adapts from [40] the general integer programming approach in such a way that it can encapsulate the general idea of sources and targets as put forward in the price-collecting Steiner tree/forest (PCST/PCSF) approaches [45, 46, 47, 48] which capture the idea of deregulated networks starting or ending at certain types of nodes, for example membrane receptors and transcription factors. Methodologically, we extend the integer programming approach of [40] (*Backes et al.*) to fractional integer programming to allow for the necessary flexibility to incorporate sources and targets. Furthermore, we show that our algorithm, DeRegNet, can be interpreted as maximum likelihood estimation under a certain natural statistical model. We demonstrate DeRegNet’s suitability as an exploratory hypothesis generation tool by applying it to TCGA liver cancer data. We introduce a personalized approach to interpreting cancer data and introduce the notion of network-defined cancer genes which allow to identify patient groups based on their similarity of their detected personalized subgraphs. The appendix *Additional File 1: Supplementary Material and Methods* furthermore contains a demonstration of the usefulness of subgraph-derived features for survival prediction. In particular, these features outperform comparable features derived from gene set enrichment indicated pathways and also improve classifiers based on clinical data alone.

## Methods and Materials

### DeRegNet: a de-novo subnetwork identification algorithm

#### Formal setting and definitions

Formally, it is given a directed graph *G* = (*V, E*), i.e. *E* ⊂ *V* × *V*, representing knowledge about biomolecular interactions in some way. To avoid certain pathologies in the models defined below, it is assumed that *G* has no self-loops, i.e. (*v,v*) ∉ *E* ∀*v* ∈ *V*. For a subset *S* ⊂ *V*, one defines δ^+^ (*S*) = {*u* ∈ *V*\*S*: ∃*v* ∈ *S*: (*v,u*) ∈ *E*} and *δ*^-^(*S*) = {*u* ∈ *V*\*S*: ∃*v* ∈ *S*: (*u,v*) ∈ *E*}, i.e. the sets of outgoing nodes from and incoming nodes into a set of nodes *S*. For a node *v* ∈ *V* one writes *δ*^±^(*v*):= *δ*^±^({*v*}). Furthermore, it is given a score function *s*: *V* → ℝ, describing some summary of experimental data available for the biomolecular entities represented by the nodes. For a given graph *G* = (*V, E*) any node labeling function *f*: *V* → ℝ is implicitly implied to be a vector *f* ∈ ℝ|^*V*^|, subject to an arbitrary but fixed ordering of the nodes (shared across all node labeling functions). In particular, with *f_v_*:= *f*(*v*) for *v* ∈ *V*, given *f,g*: *V* → ℝ, one can write 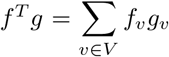. For *S* ⊂ *V* and *f*: *V* → ℝ one defines *f_S_*: *V* → ℝ via *f_S_*(*v*):= 0 for all *v* ∈ *V* \ *S* and *f_S_*(*v*):= *f*(*v*) for all *v* ∈ *S*. Defining *e*: *V* → ℝ with *e*(*v*):= 1 for all *v* ∈ *V*, one further can write 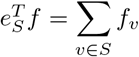 for *S* ⊂ *V* and *f*: *V* → ℝ. Comparison of node labeling functions *f, g* are meant to be understood element-wise, e.g. *f* ≤ *g* means *f_v_* ≤ *g_v_* for all *v* ∈ *V*. Apart from the graph *G* and node scores *s*, there are given possibly empty subsets of nodes *R* ⊂ *V* and *T* ⊂ *V*. It is referred to *R* as *receptors* (or sometimes *sources*) and to *T* as *terminals* (or sometimes *targets*), independent of the biological semantics underlying the definition of these sets (see below). For enforcing the topology of the subnetworks later on, strongly connected components will play a decisive role and it is said that a subset of nodes *V*′ ⊂ *V* induces a strongly connected subgraph (*V*′ *iscs*, for short) if the subgraph induced by *V*′ is strongly connected.

#### Probabilistic model for deregulated subgraphs

The mathematical optimization model which is at the heart of the DeRegNet algorithm and presented in the next subsection amounts to maximum likelihood estimation under a certain canonical statistical model. The model assumes binary node scores *s*: *V* → {0,1} which are realizations of random variables **S** = (*S_v_*)_*v*∈*V*_. Here, *S_v_* = 1 is interpreted as node *v* ∈ *V* being *deregulated*. Further it is assumed the existence of a subset of vertices *V*′ ⊂ *V* such that *S_v_* |*v* ∈ *V*′ ~ Ber(*p*′) and *S_v_*|*v* ∈ *V*\*V*′ ~ Ber(*p*) with *p,p*′ ∈ (0,1) denoting probabilites of deregulation outside and inside of the deregulated subgraph encoded by *V*′ respectively. It is assumed that *p*′ > *p* to reflect the idea of *higher* deregulation (probability) in the *deregulated* subgraph, whereas *p* represents a certain amount of background deregulation. The network context (dependency) is introduced via the restriction that 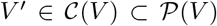. Here, 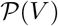 is the power set of *V* (the set of subsets of *V*) while 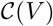 as a subset of 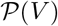 represents the set of feasible substructures and should (can) reflect topologies inspired by known biomolecular pathway topologies like the one described in Backes et al. [40] and the next subsection. Furthermore it is assumed, that the (*S_v_*), given a network context and deregulation probabilities *p,p*′, are independent. We show in the appendix that under this model and the constraints given by the fractional integer programming problem formulated in the next subsection (defining 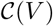 in the above notation) DeRegNet amounts to maximum likelihood estimation. Furthermore, we also show that the model put forward in Backes et al. [40] amounts to maximum likelihood estimation only under the assumption of a fixed subgraph size.

#### Finding deregulated subgraphs by fractional integer programming

Given the definitions of the preceding sections, we can now formulate the main model underlying DeRegNet. The DeRegNet model and also the model of Backes et al. [40] can be placed in the context of the so called *Maximum Weight Connected Subgraph Problem (MWCSP),* see Additional File 1: Supplementary Material and Methods. Note that in the following we formulate all problems as maximization problems and minimization may, depending on the semantics of the node score, be the proper choice. Minimization may for example be prudent in case the node scores represent p values originating from some statistical significance test. As Backes et al. [40] we model the problem of finding deregulated subnetworks in terms of indicator variables *x_v_* = **I**(*v* ∈ *V*′) and *y_v_* = **I**(*v* is the root node) where *V*′ ⊂ *V* is a set of nodes inducing a subgraph such that one can reach every node in that subgraph by means of a directed path from the root node. Here, **I**(*P*) = 1 if *P*, **I**(*P*) = 0 if not *P* for some predicate *P*. In addition the root is supposed to be a source node and all nodes in the subgraph with no outgoing edges are supposed to be terminal nodes. The proposed model then reads like this:

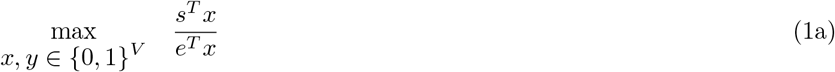

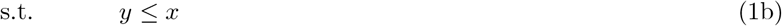

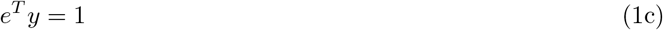

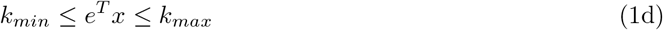

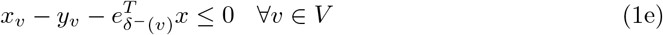

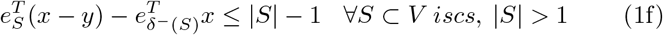

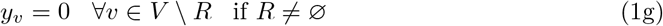

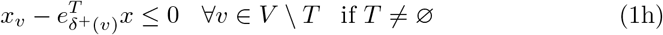

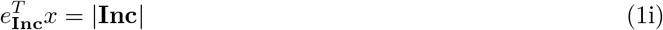

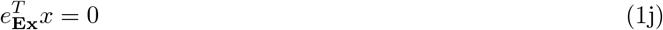

The model derives from the corresponding integer linear programming model in [40] and adapts it for the fractional case, most notably here are the constraints involving the receptors *R* (1g) and the terminals *T* (1h). (1g) just ensures that the root node is a receptor while (1h) ensures that any node in the subgraph with no outgoing edges is a terminal node. (1b) means that a node can only be the root if it is included in the subgraph, (1c) means that there is exactly one root, (1d) means that the size of subgraph has to be within the bound given by 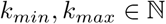, (1e) says that a node *v* ∈ *V* in the subgraph is either the root node or there is another node *u* ∈ *V* in the subgraph such that there is an edge (*u,v*) ∈ *E*. Moreover, the the constraints (1i) and (1j) trivially allow to include and exclude specific nodes from given node sets **Inc** ⊂ *V* and **Ex** ⊂ *V* respectively. In many situations specific nodes, i.e. genes in the case of gene regulatory networks, may be of interest in other topological positions than in a receptor or terminal role. In that case just requiring a certain gene to be part of the subgraph without any special constraints on its inclusion in topological terms can be of value. The constraint (1f) is the most involved one and actually describes exponentially many constraints which ensure that there are no disconnected directed circles ([40]) by requiring that any strongly connected component in the subgraph either contains the root node or has an incoming edge from another node which is part of the subgraph but not part the given strongly connected component. Finally, the objective (2.1a) describes the notion of maximizing the average score of the subgraph. This is crucial for allowing the model the flexibility to connect source nodes to target nodes and also is at the heart of DeRegNet being able to do Maximum Likelihood estimation given the presented statistical model. We summarize some crucial terminology next, before proceeding in the next subsection to describe the solution algorithms for DeRegNet.

#### Definition 1

(DeRegNet instances, data, and subgraphs)

*A tuple* (*G,R,T, **Ex**, **Inc**, s*) *is called an **instance of DeRegNet** (a **DeRegNet instance**, an **instance of the DeRegNet model**). Here, G* = (*V, E*) *is the **underlying graph**, R* ⊂ *V is the **receptor set**, T* ⊂ *V is the **terminal set**, **Ex*** ⊂ *V is the **exclude set**, **Inc*** ⊂ *V is the **include set** and s*: *V* → ℝ *is the **node score** (the **score**). Further, x_v_*: *V* → {0,1} *is called a **subgraph** with the understanding that it is referred to the subgraph of G induced by V*\* = {*v* ∈ *V*: *x_v_* = 1}. *Equivalently to x_v_*: *V* → {0, 1}, *it is also referred to the corresponding V*\* = {*v* ∈ *V*: *x_v_* = 1} *as a subgraph. A subgraph is **feasible for DeRegNet** (for the DeRegNet instance), if it satisfies DeRegNet’s constraints (1b-j). A subgraph satisfying these constraints is called a **feasible subgraph**. A feasible subgraph which optimizes problem (1) is called an **optimal subgraph**.*

Some formal properties of DeRegNet and its solutions can be found in the *Additional File 1: Supplementary Material and Methods*. A high-level depiction of the overall logic of DeRegNet can be found in figure 1. A conceptual view of the types of subgraphs determined by DeRegNet can be seen in Figure 3 whereas the high-level algorithm of DeRegNet is summarized by Algorithm 1.

**Figure 1:**
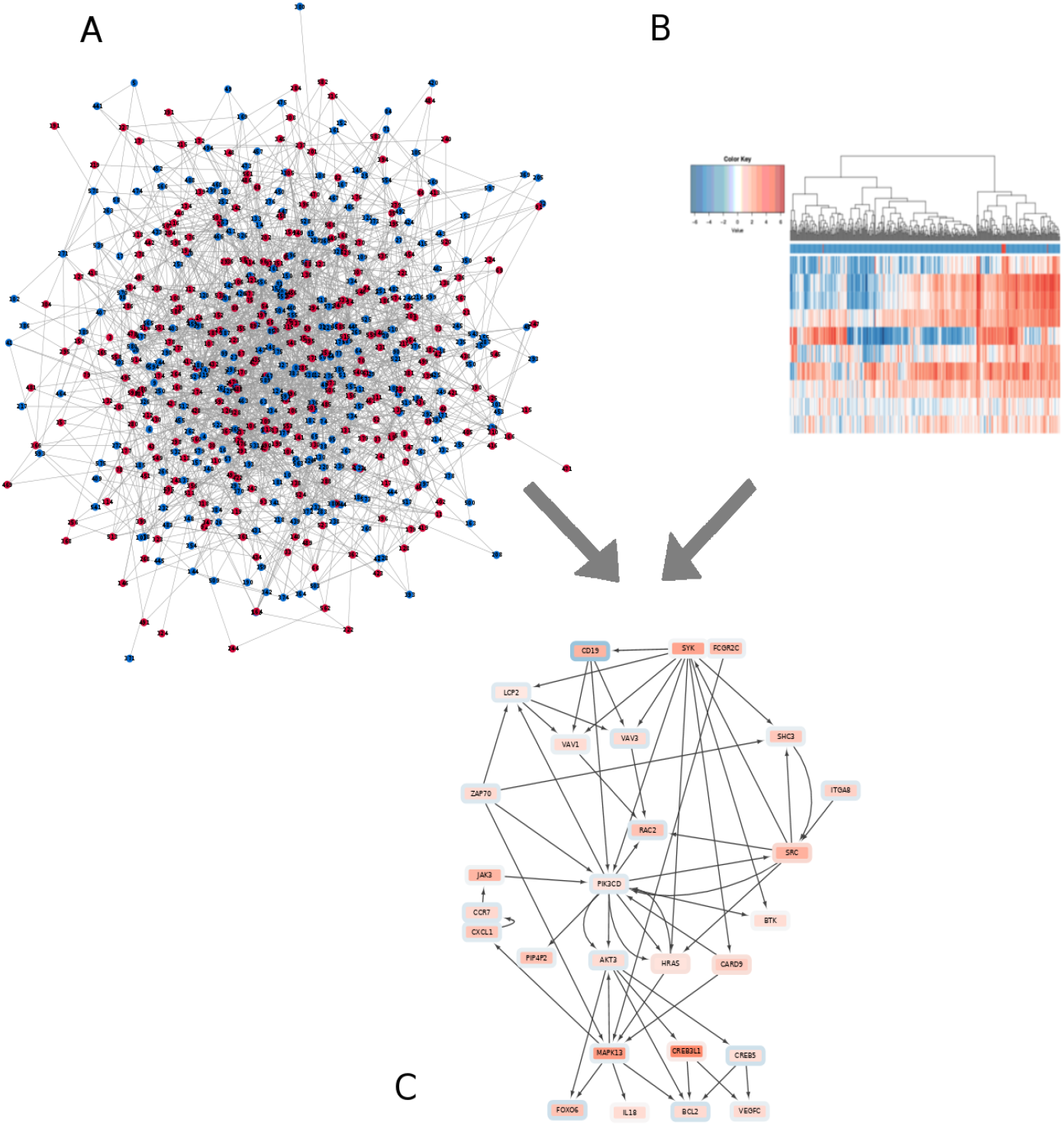
DeRegNet’s inputs are a biomolecular network **(A)**, such as a signaling or gene regulatory network, and omics measurements **(B)**, such as gene expression data. The latter are mapped onto the nodes of the network acting as node-level measures of deregulation. DeRegNet then extracts the most deregulated subnetwork from the larger regulatory network according to some definition of *most deregulated.* For a conceptual view of the progression from set enrichment to de novo subnetwork methods we refer to Figure 2

**Figure 2:**
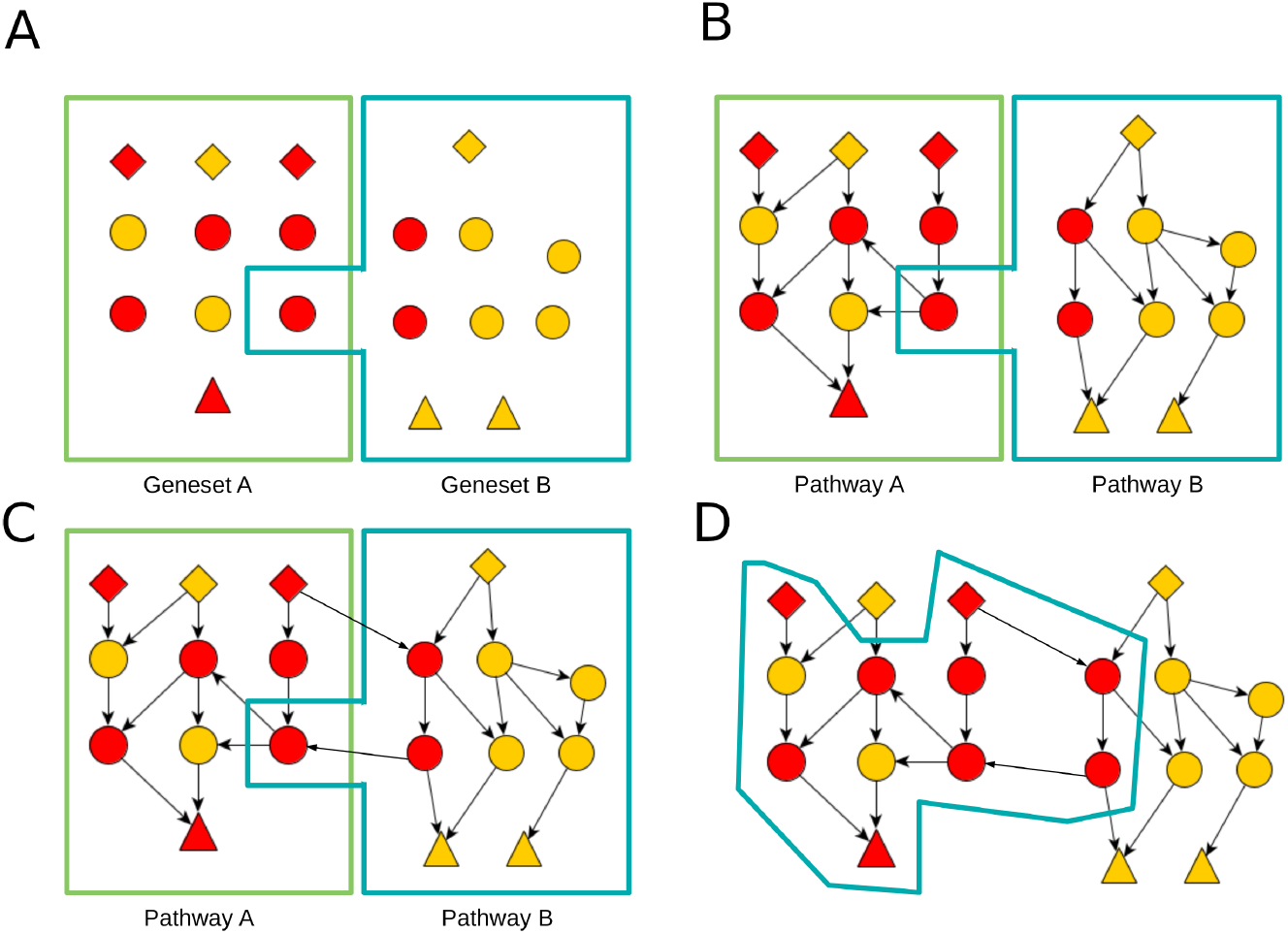
Conceptual progression from gene set enrichment to de-novo subnetwork enrichment. **(A)** Classical gene set enrichment. See Supplementary Figure S1 for additional details. **(B)** Topological pathway enrichment. See Supplementary Figure S2 for additional details. **(C)** Topological pathway enrichment with pathway cross-talk. See Supplementary Figure S3 for additional details. **(D)** De-novo subnetwork/pathway enrichment. See Supplementary Figure S4 for additional details.

**Figure 3:**
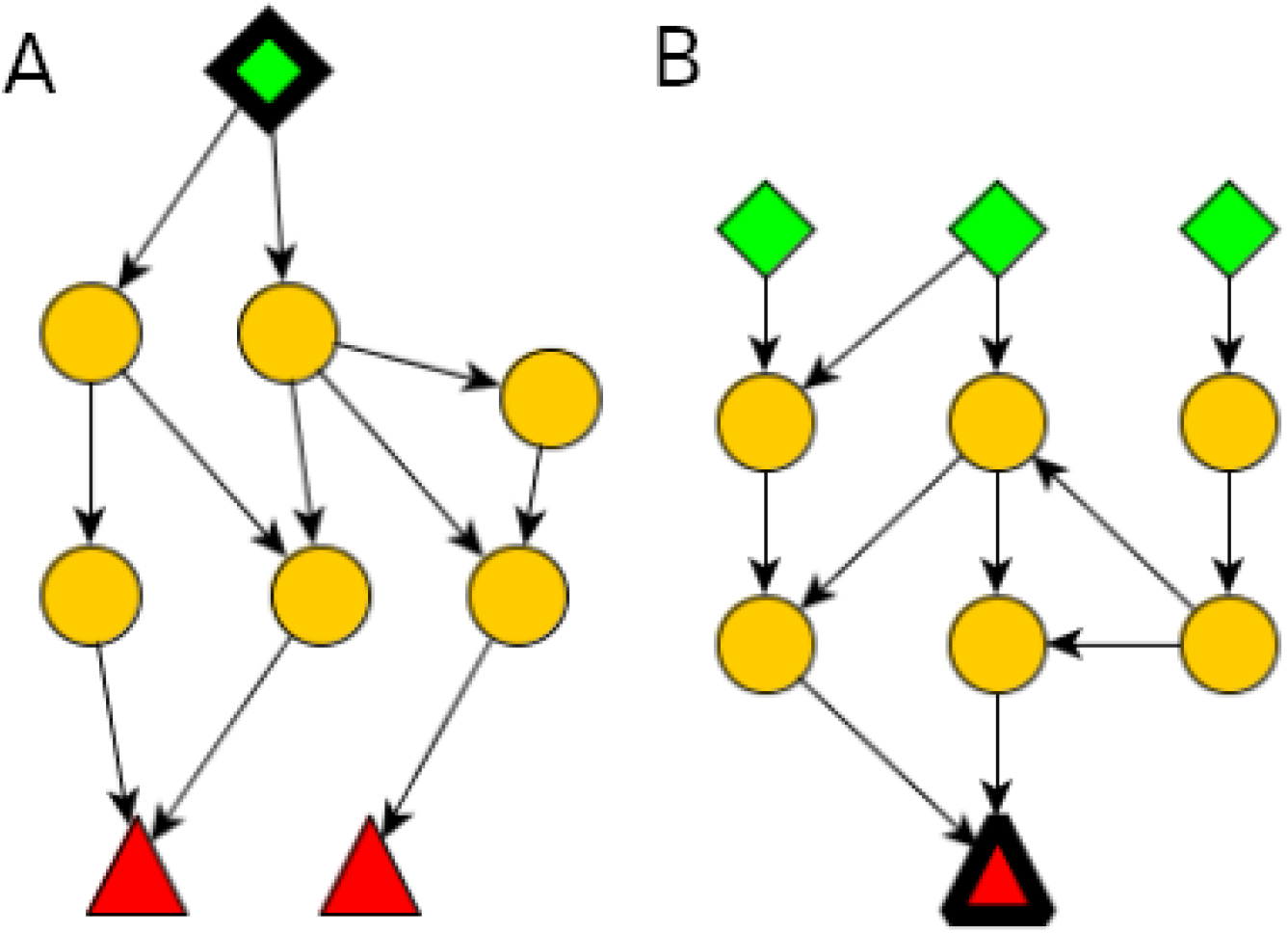
Conceptual view of subgraphs extracted by DeRegNet. **(A)** From a receptor node/root node (green cube) one can reach any node in the subnetwork. Nodes without any edges leading to other nodes (red triangles) of the subnetwork need to be elements of the so called terminal nodes. Generally, all nodes in the subgraph can be reached from the root node. **(B)** By reversing the orientation of the underlying network before applying DeRegNet, one can find subgraphs with only one terminal “root” node and multiple receptor nodes such that the terminal node can be reached from any other node in the subgraph. See *Additional File 1: Supplementary Material and Methods* for further details on applying DeRegNet in reverse mode.

**Algorithm 1:**
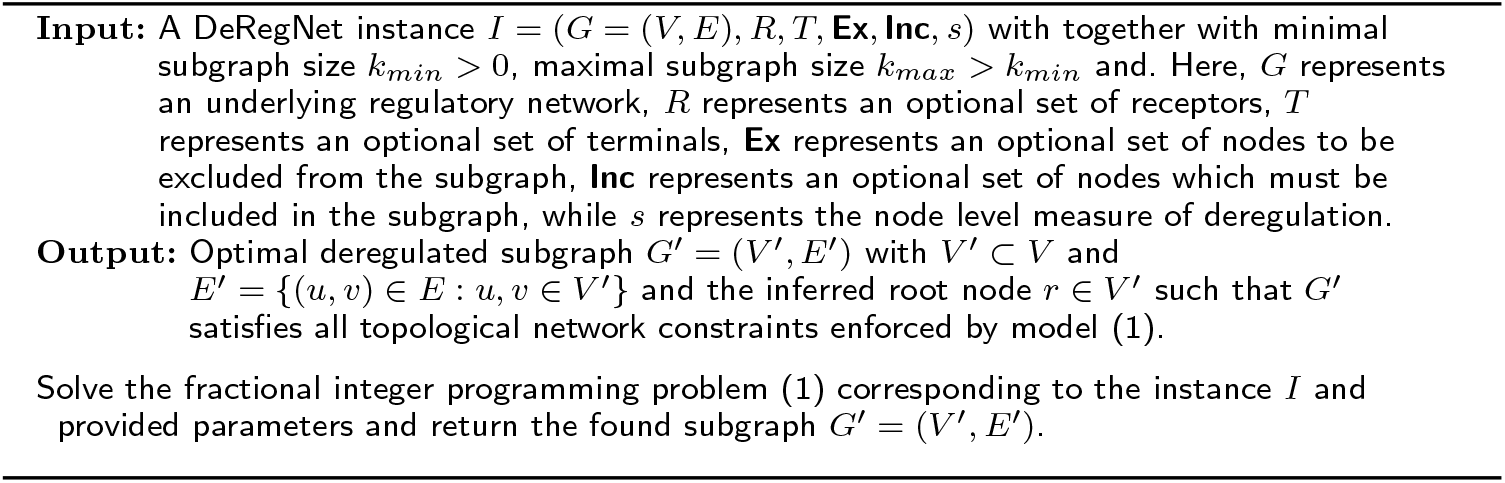
DeRegNet deregulated subnetwork detection. Overall high-level algorithmic procedure underlying de-novo pathway/subnetwork detection with DeRegNet. Note that resulting subgraph can additionally scored with a P-Value obtained by a GSE method interpreting the found subgraph as a gene set.

#### Solving the fractional integer programming model

We solve the integer fractional linear programming problems introduced in the previous sections by one out of two implemented methods. Firstly, a generalization of the Charnes-Cooper transformation [49] for fractional linear programs described by [50] and secondly an iterative scheme as introduced generally by Dinkelbach [51, 52] and subsequently applied in the context of integer fractional programming by [53, 54]. While the Dinkelbach-type algorithm solves the problem by iteratively solving certain non-fractional versions of the original problem until some convergence criterion is met, the generalization of the Charnes-Cooper method is based on reformulation of the entire fractional model to a quadratic problem and requires subsequent linearization of artifically introduced quadratic constraints. The latter is implemented in terms of the methods described by [55, 56, 57].

As in [40] the exponentially many constraints forbidding any strongly connected components not containing the root and with no incoming edges are handled by lazy constraints. Every time an integer solution is found the Kosaraju-Sharir algorithm [58] is employed (as implemented by the Lemon graph library) to check for violating components and, in the case of violating components, the corresponding constraints are added to the model. Both solution approaches, the generalized Charnes-Cooper method and the Dinkelbach-type algorithm, allow for the lazy constraints to be handled in terms of the original formulation since both retain the relevant variables of the model within the transformed model(s).

For more details on the theoretical underpinnings and the practical implementation of DeRegNet’s solution algorithms consult *Additional File 1: Supplementary Material and Methods*.

### Assessment of inference quality for known ground truths

The evaluation and benchmarking of *de novo* pathway enrichment or deregulated subnetwork detection algorithms and implementations remains a big challenge. While many of the methods cited in the introduction can be applied to reveal useful biological insight, there are limited studies concerning the comparison of formal and statistical properties of the methods. The two main obstacles are a lack of well-defined gold standard datasets as well as the differences concerning the exact output of the methods. For example, it is not immediately clear how to compare algorithms which produce undirected subnetworks to those which elicit directed networks of a certain structure. An important first step toward atoning the issue in general is described in [19] which focuses on benchmarking approaches for undirected networks. For the purposes of this paper, we designed and performed benchmarks of DeRegNet relative to its closest relative, namely the algorithm described in [40], henceforth referred to as *Backes et al.*. Note however, while we are comparing the integer programming based algorithm of Backes et al. to the fractional integer programming algorithm of DeRegNet, we are using the former as implemented in the DeRegNet software package. This renders the benchmark less dependent on implementation technology since both algorithms have been implemented with the same general stack of languages and libraries. For the benchmark we always utilize the human KEGG network as the underlying regulatory network. We then repeatedly simulate subgraphs which match the structure of both models (DeRegNet and Bckes et al.). The simulation procedure is described more formally in *Additional File 1: Supplementary Material and Methods*. Initially, the simulated subgraph consists of one randomly selected root node, to which we iteratively add a random ”outgoing” neighbor of a randomly selected current node in the subgraph until the size of the subgraph matches a randomly chosen value. The latter is uniformly chosen to be an integer between a given lower and an upper bound. ”Outgoing” neighbors of *v* ∈ *V* are any nodes from the set *δ*^+^(*v*) = {*u* ∈ *V*\{*v*}: (*v,u*) ∈ *E*}. All nodes in the simulated ”real” subgraph are assigned a node score of 1 with a certain probability *p*′ > 0, while all nodes which are not contained in the subgraph are assigned a node score of 1 with probability *p* where 0 < *p* < *p*′. In summary, we obtain random “real” subgraphs and simulated scores where the latter reflect the different likelihood of a node being deregulated given whether it is contained in the subgraph or not. In terms of the probabilistic interpretation of DeRegNet presented above, the simulation scheme corresponds directly to a deregulation probability of *p*′ for nodes in the ”real” subgraph and of *p* for nodes not part of the ”real” subgraph. The appendix in *Additional File 1: Supplementary Material and Methods* provides further details on the simulation of benchmark instances.

Given a sequence of 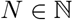 of these simulated instances, the algorithms are run in order to find subgraphs which can then be compared to the known simulated real subgraph. Here, a *hit* (*true positive, tp*) is defined as a node appearing in a subgraph calculated by some algorithm which is also an element of the real subgraph. A *false positive* (*fp*) is a node which appears in a subgraph calculated by an algorithm but is not part of the real subgraph. A *false negative* is defined as a node which is part of the true subgraph but not part of the subgraph detected by an algorithm. Furthermore, we can compare the sizes of the calculated subgraphs with the size of the real subgraph. In general, given an algorithm 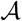, which on a given instance with true subgraph *V*′ ⊂ *V* finds a subgraph 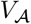, one can adopt all standard evaluation metrics for predictive classification models with the understanding that nodes in 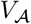 are *predicted positive* and nodes in *V*′ are *true positive.* Examples are the true positive rate (sensitivity) 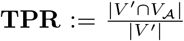, i.e. the number of actual hits divided by the number of possible hits, or the Jaccard index (intersection over union) 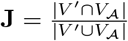. Specifically, we utilize the Matthews correlation coefficient (MCC), the F1 score, the Jaccard index, precision and sensitivity to compare subgraphs found by DeRegNet and Backes et al. The only non-standard metric we employ compares the closeness of an inferred subgraph to a real subgraph and is referred to as size efficiency 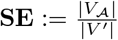, i.e. the proportion algorithm subgraph size to real subgraph size. Another comparison metric is the running time of the algorithms.

Furthermore, the benchmark is based on the realistic assumption that we do not know the exact size of the real subgraph and that one can only assume lower and upper bounds on the subgraph size instead. Since the Backes et al. algorithm does need a fixed a priori specified subgraph size we employ a strategy suggested in [40] to circumvent that fact. Namely, we iterate from the lower to the upper bound, find a subgraph for each subgraph size and then regard the union graph of all found subgraphs as the one subgraph emitted by the algorithm. DeRegNet natively requires only a lower and an upper bound on subgraph size as parameters. All benchmarks have been carried out with the following setup: software: Ubuntu 18.04, Gurobi 9.5.0, hardware: 12x Intel i7-8750H @ 4.1 GHz, 32 GB RAM, Samsung SSD 970 EVO Plus. See *Additional File 1: Supplementary Material and Methods* for more formal details. Finally, in order to assess the comparative advantage of deregulated subgraphs to pre-defined pathways (gene sets) we calculated GSE P-values for optimal DeRegNet subgraphs (interpreted as gene sets) based on the simulated scores, as well as for the standard KEGG gene sets and compared the distribution of subgraph P-values with those of significant KEGG gene sets across all simulation runs.

### Network and omics data

#### KEGG network

While many sources for directed biomolecular networks are available, e.g. [59], in this paper we here utilize a directed gene-level network constructed from the KEGG database with the KEGGgraph R-package [60]. The script used to generate the network as well as the network itself can be found in the DeRegNet GitHub repository. See the subsection on Software Availability for details.

#### RNA-Seq and 450k methylation array derived node scores for TCGA-LIHC

Gene expression and methylation data was downloaded for hepatocellular carcinoma TCGA project from the Genomic Data Commons Portal (https://portal.gdc.cancer.gov/projects/TCGA-LIHC). Raw quantified RNA-Seq counts were normalized with DESeq2 [61] which was also used for calculating log2 fold changes for every gene with respect to the entire cohort. We also used DESeq2 to calculate P-Values for differential expression between tumor and control samples. Personalized log2 fold changes were calculated by dividing a patients tumor sample expression by the mean of all available control samples (adding a pseudo count of 1) before taking the log. The following node scores are defined:

- *Global RNA-Seq score s: s_v_* = RNA-Seq log_2_-fold change for a gene *v* ∈ *V* as calculated by DESeq2 for the TCGA-LIHC cohort
- *Trinary global RNA-Seq score*:

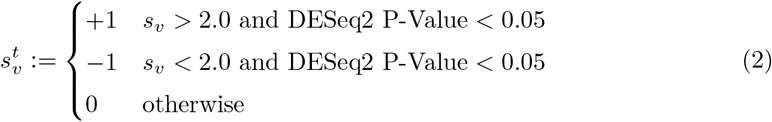
- *Trinary personalized RNA-Seq score s^c^* for case *c*:

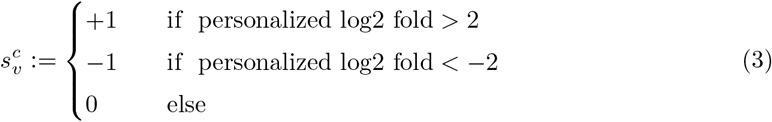

From the 450k methylation array [62, 63] data available for the TCGA-LIHC cohort we derive (global) methylation node/gene scores *m_v_* ∈ { –1,0,1} for every gene *v* ∈ *V* representing binary methylation status as follows. First signed differentially methylated probes (DMPs) were inferred using Subset Quantile Normalization (SQN) [63] between tumor and control samples. With *signed* we express the fact, that we keep track whether the median difference between tumor and control was positive or negative. Correspondingly, median(*β*_1_,…, *β_T_*) – median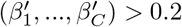 defines a upregulated DMP while median(*β*_1_,…,*β_T_*) – median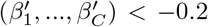 defines a downregulated DMP. Here, *β*_1_, …,*β_T_* ∈ [0,1] denote all beta values from tumor samples for a given array probe while 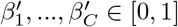 denote all beta values from control samples for that same probe. From the DMPs’ metadata contained in the TCGA-LIHC 450k data one obtains a mapping from any DMP to genes to whose promoter region the DMP lies close to (up to 1500 base pairs upstream of a genes transcription start site). Any gene *v* ∈ *V* which is indicated by at least one upregulated DMP and no downregulated DMPs is considered upregulated and we set *m_v_*:= 1. Any gene *v* ∈ *V* which is indicated by at least one downregulated DMP but no upregulated DMPs is considered downregulated and we set *m_v_*:= −1. For genes which are not up- or downregulated we set *m_v_*:= 0.

### Global and personalized deregulated subgraphs

We refer to subgraphs found with the global RNA-Seq score s as *global subgraphs*. A global subgraph can further be subdivided as being *upregulated* or *downregulated* depending on whether the subgraphs were found by employing a maximization or minimization objective respectively. For (any) node score *s*: *V* → ℝ we define |*s*|: *V* → ℝ by |*s*|(*v*):= |*s*(*v*)| for all *v* ∈ *V*. *Dysregulated* global subgraphs are those which were found by using the score |*s*| under a maximization objective. Similarily subgraphs found with any of the scores *s^c^* with a maximization objective are called *upregulated* while those found with minimization objective are called *downregulated* (personalized subgraphs for case/patient *c*). Subgraphs found with a |*s^c^*| score under maximization are called *dysregulated* (personalized subgraphs for case/patient *c*). Any of the above subgraph types is called a *deregulated* subgraph. Subgraphs were inferred with minimal subgraph size of *k_min_* = 10 and maximal subgraph size of *k_max_* = 50 as this represents a reasonable range of expected pathway sizes, compare *Supplementary Figure S24*. The optimal and four next best suboptimal global subgraphs were calculated for every modality. The subgraphs were then summarized as a subgraph of the union graph of optimal and suboptimal subgraphs in order to allow streamlined interpretation. See the supplementary figures referenced in the respective figures for references to the direct output of DeRegNet.

### Network-defined cancer genes

Genes, gene products or biomolecular agents are likely to bring about their various phenotypic effects only in conjunction with other agents via their shared biomolecular network context. By that token, one can search for genes which convey phenotypic differences by means of some defined network context. Here, we propose DeRegNet subgraphs as network context for a given case/patient in order to find genes whose inclusion into a case’s deregulated subgraph associates with a significant difference in overall survival. Algorithm 2 describes the procedure more formally. Genes implicated by the outlined procedure are termed *network-defined cancer genes*. The next section provides details on a specific network-defined cancer gene obtained by application of the procedure to personalized upregulated subgraphs in the TCGA-LIHC cohort.

**Algorithm 2:**
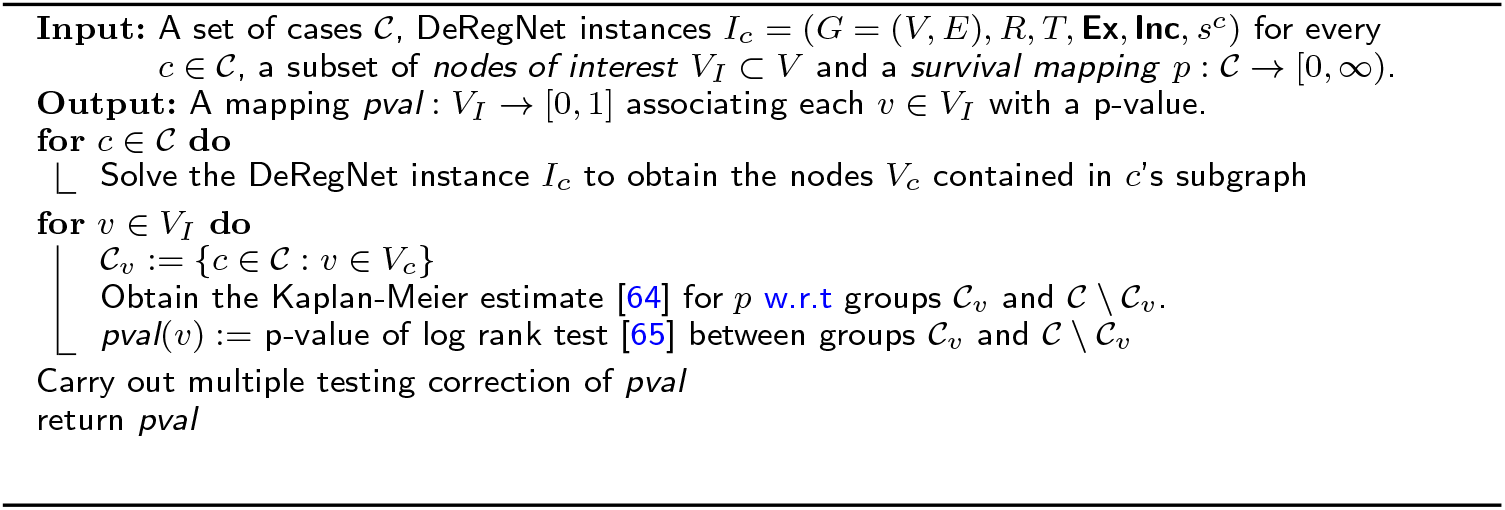
Finding subnetwork-defined cancer genes. After finding subgraphs for individual cases/patients the procedure partitions a set of cases/patients according to whether they contain a given gene in their determined subnetwork and tests whether the thus defined partition conveys a significant survival difference. Note, that in the described setting, the DeRegNet instances only differ in terms of their case-dependent node score *s^c^*.

### Nodes scores representing consistent methylation and transcription patterns

In general one considers a *consistent methylation and transcription pattern* for a given gene a situation where one observes increased methylation (hypermethylation) close to/in the gene’s promoter region and decreased transcription of the gene or decreased methylation (hypomethylation) close to/in the gene’s promoter region and increased transcription of the gene [66], [67]. For a node/gene *v* ∈ *V* we define a node score 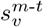 which captures these patterns by 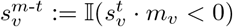, i.e.

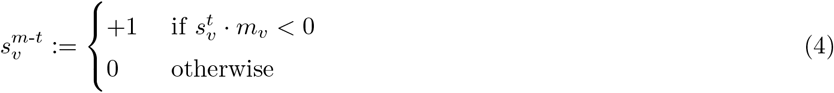

We then infer deregulated subgraphs with nodes scores 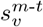 for *v* ∈ *V* in order to capture subnetworks which consist largely of nodes which show consistent methylation-transcription patterns, thus representing de-novo pathways which may be largely regulated by epigenetic modulation of transcription.

## Results and discussion

In the following we present multiple results relating to the application of the DeRegNet algorithm. Firstly, we present benchmark results for synthetic data which compares DeRegNet to its closest methodological relative [40]. Next, we present applications of DeRegNet on a TCGA liver cancer dataset. More specifically, we present global subgraphs for the TCGA representing deregulated subnetworks summarizing the cohort under study as a whole, as well as a personalized application of DeRegNet, i.e. the derivation of patient-specific subgraphs.

### Performance comparison on data with a known ground truth

As outlined in the introduction, the field of statistical functional annotation needs adequate known ground truths (gold standards) against which one can evaluate corresponding methods, see for example [19]. Since actual ground truths are hard to come by for fundamental reasons, research for functional annotation algorithms justifiably focuses on simulated/synthetic ground truths. The latter are then generated such that they represent the assumed or postulated data-generating process. We compared DeRegNet to its closest methodological relative introduced in [40] based on simulated instances as described in *Material and Methods*.

Figures 4 and 5 show results of simulation runs carried out according to the described procedure. As can be seen in Figure 4, DeRegNet outperforms [40] in terms of Matthews Correlation Coefficient (MCC), F1 Score, Jaccard index, Precision, subgraph size efficiency (i.e. closeness to true subgraph size) and running time. Backes et al. has almost perfect sensitivity but DeRegNet generally also performs close to optimal with only a few outliers with lower sensitivity compared to Backes et al.. The Backes et al. algorithm achieves these slight sensitivity advantages with considerable cost with respect to precision as can be seen in Figure 5. In order to assess the dependence of these simulation results on certain simulation parameters, in particular the noise level *p, Supplementary Figures S18-S23* can be consulted. For a wide range of noise settings, DeRegNet outperforms Backes et al. with the described patterns for the evaluation metrics. With increasing noise levels both algorithms start to perform less convincing.

**Figure 4:**
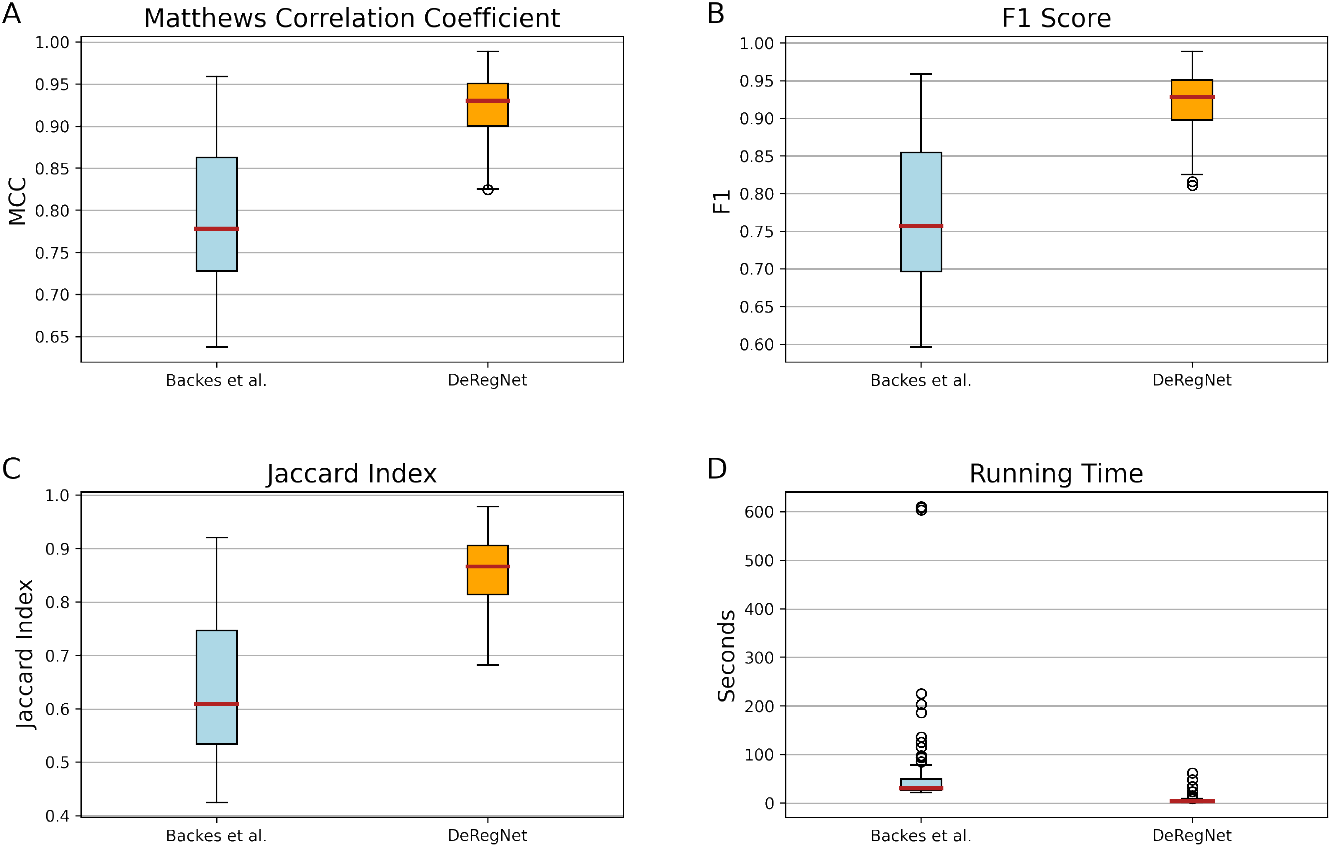
Benchmark patterns for DeRegNet and Backes et al. [40]. **(A)** Distributions of the Matthews correlation coefficients of DeRegNet and Backes et al. subgraphs. **(B)** Distributions of the F1 scores of DeRegNet and Backes et al. subgraphs. **(C)** Distributions of the Jaccard indices of DeRegNet and Backes et al. subgraphs. **(D)** Running time (in seconds) of DeRegNet (Dinkelbach algorithm) and *k_max_* – *k_min_* + 1 (*k_max_* = 50, *k_min_* = 25) runs of the Backes et al. algorithm [40]. Benchmark parameters: in-subgraph deregulation probability *p*′ = 0.99, out-of-subgraph deregulation probability p = 0.1, *k_min_* = 25, *k_max_* = 50, minimal size of simulated true subgraph = 30, maximal size of simulated true subgraph = 45, number of simulated instances = 100, time limit = 600 seconds. See *Additional File 1: Supplementary Material and Methods* for further formal details on the simulation procedure and *Supplementary Figures S18-S24* for simulation results for different parameter value settings.

**Figure 5:**
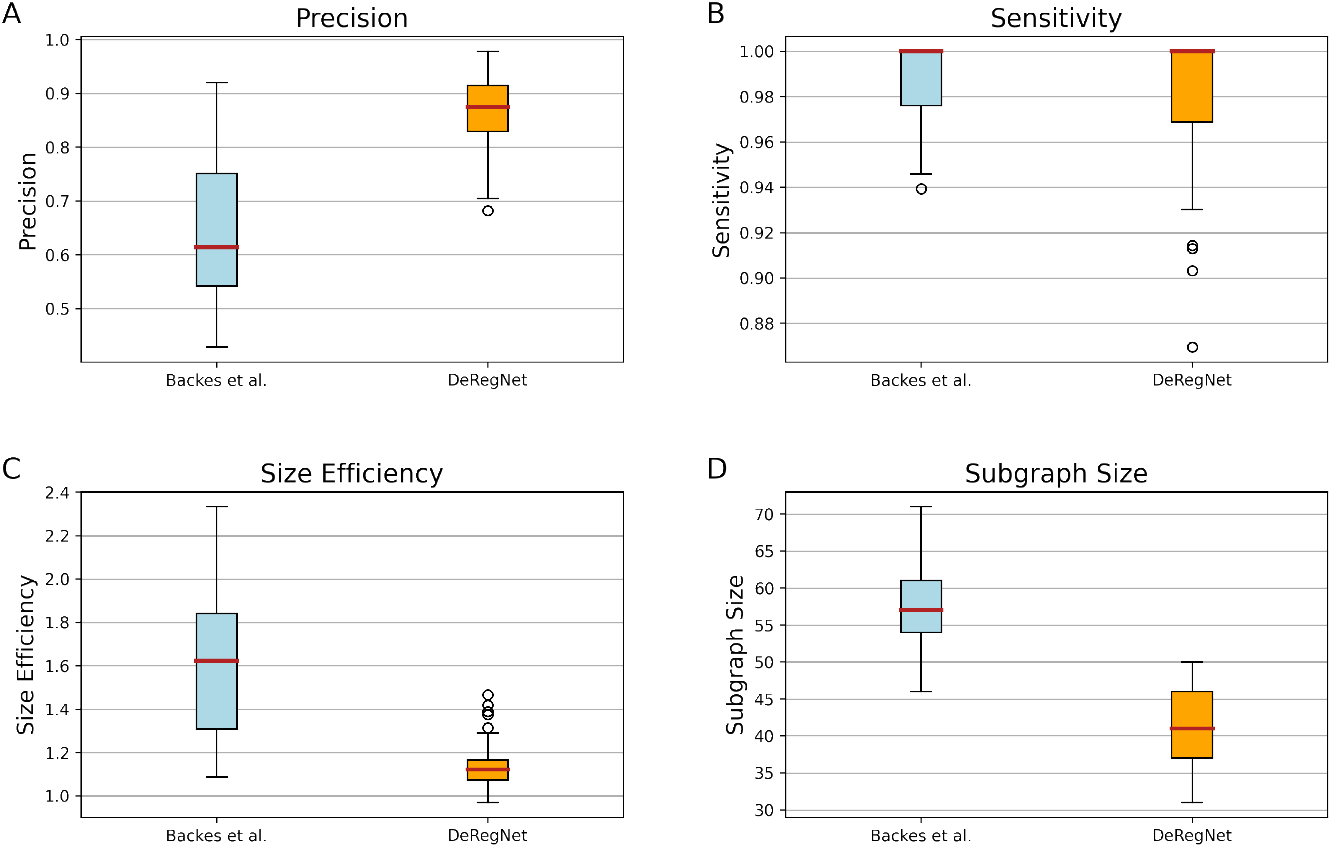
Benchmark patterns for DeRegNet and [40]. **(A)** Distributions of the precision of DeRegNet vs. Backes et al. subgraphs. **(B)** Distributions of the sensitivity of DeRegNet vs. Backes et al. subgraphs. **(C)** Distributions of the size efficiency of DeRegNet vs. Backes et al. subgraphs. **(D)** Distributions of subgraph sizes of DeRegNet vs. Backes et al. subgraphs. Benchmark parameters: As in Figure 4. See *Additional File 1: Supplementary Material and Methods* for further formal details on the simulation procedure.

Furthermore, all optimal subgraphs (interpreted as gene sets) found by DeRegNet (or Backes et al.) are significant w.r.t classical GSE analysis and considerably more so than pre-defined KEGG gene sets. This underlines the suitablility of denovo subnetwork/pathway detection algorithms to find significant data-dependent ”pathways” in the context of the outlined simulation studies. See Figure 6.

**Figure 6:**
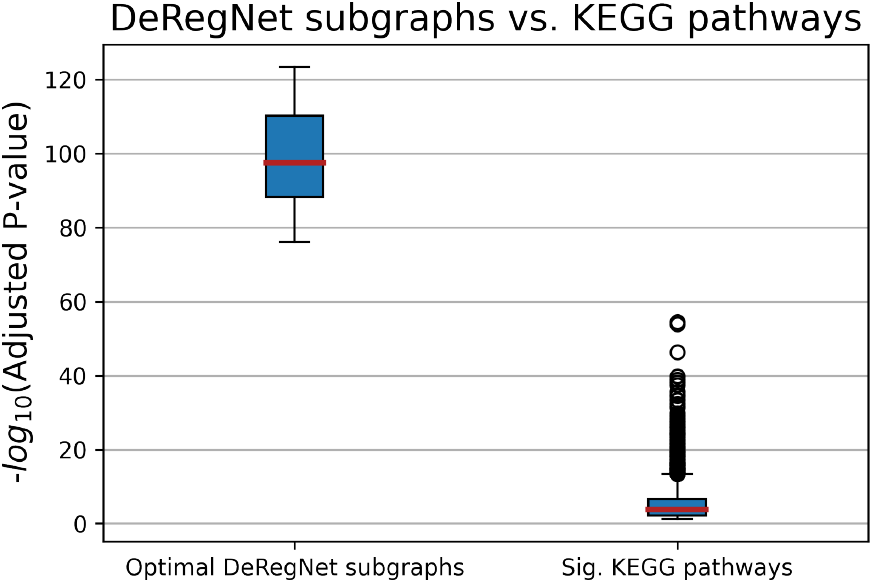
DeRegNet subgraphs vs. pre-defined KEGG subgraphs. Distributions of significant P-Values for optimal deregulated subgraphs found by DeRegNet and pre-defined KEGG gene sets. Note that DeRegNet subgraphs (interpreted as gene sets) are considerably more significant than pre-defined (non data-dependent) KEGG gene sets in light of classical GSE. Additionally, all DeRegNet subgraphs were significantly enriched without exception.

Less quantitatively, note that DeRegNet allows for subgraphs which originate from so called source (root, receptor) nodes and *end* at so called terminal nodes. This is not readily possible with the Backes et al. algorithm due to the necessity to specify a fixed subgraph size *a priori* and the resulting lack of flexibility to connect receptors to targets. Also note that DeRegNet is available as open-source software and also provides an open-source implementation of the Backes et al. algorithm. Currently the implementation supports the commercial Gurobi ILP solver as a solver backend. Gurobi readily provides free academic licenses though. Furthermore, given the statistical model introduced in *Material and Methods*, Backes et al. solves only a special case of the maximum likelihood estimation problem which is solved by DeRegNet in its general form.

### Global deregulated subgraphs TCGA-LIHC

Using the DeRegNet algorithm we determined the upregulated global subgraphs obtained from running the algorithm with the global RNA-Seq score defined above. The optimal and four next best suboptimal subgraphs were calculated for every modality. The subgraphs were then summarized as a subgraph of the union graph of optimal and suboptimal subgraphs in order to allow streamlined interpretation. The global subgraph comprised of upregulated genes as nodes is shown in Figure 7.

**Figure 7:**
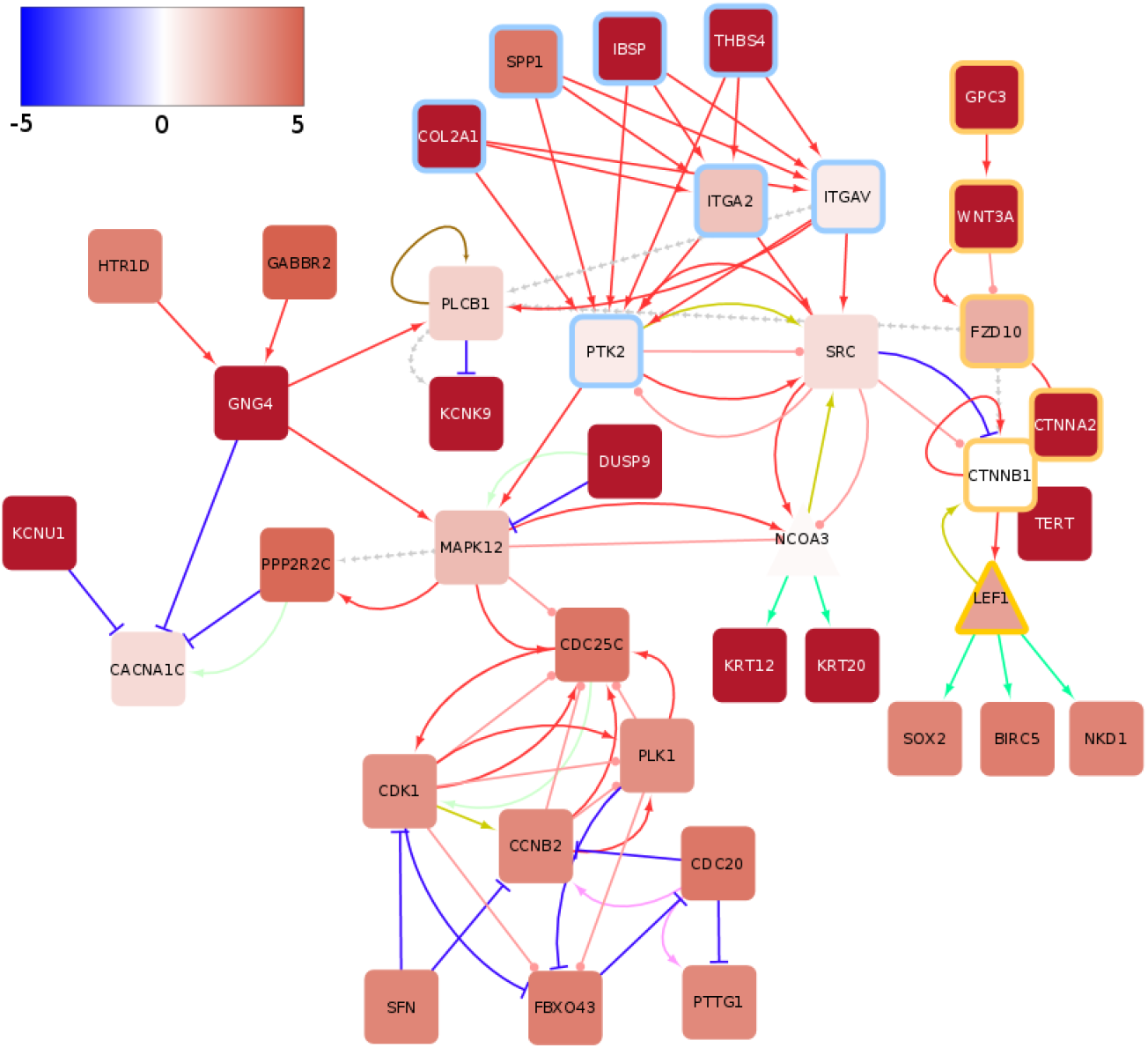
Global upregulated subgraph for TCGA-LIHC reconstructs transcriptional activation of *WNT* signaling. The Color of nodes indicates the average log_2_ fold change of tumor samples compared to controls as represented in the color bar. The color of rims around nodes indicates genes contained in the integrin pathway (blue), the *WNT* pathway (yellow) and diverse other pathways (no rim). The color of edges indicates following interactions: activation (red), inhibition (dark blue), compound (brown), binding/association (yellow), indirect effect (dashed grey), phosphorylation (pink), dephosphorylation (light green), expression (green) and ubiquitination (light purple). See also supplementary figures S6-S10

#### Reconstruction of transcriptional activation of WNT signaling

The subgraphs shows the activation of the *WNT* signaling pathway by means of over-expressed Glypican-3 (*GPC3*), which represents a membrane-bound heparin sulphate proteoglycan [68]. *GPC3* has been extensively researched as a early biomarker and potential therapy target in HCC [69, 70, 71, 72, 73, 74] (See Figure S5 in Additional File 2: Supplementary Figures).

Genomic analysis conducted over the past decade have identified mutations affecting Telomere Reverse Transcriptase (*TERT*), *β*-catenin (*CTNNB1*) and cellular tumor antigen *p53* (*TP53*) [75] as common driver mutations in HCC. Mutations in the *TERT* promoter are a well-studied factor in liver cancer development [76, 77] and lead to *TERT* overexpression while mutations in *CTNNB1*, activate *CTNNB1* and result in activation of *WNT* signaling. Previous studies have determined that *TERT* promoter mutations significantly co-occur with *CTNNB1* alternation and both mutations represent events in early HCC malignant transformation [78]. In agreement, the DeRegNet algorithm recatures the importance of a *CTNNB1*:*TERT* connection on a transcriptional level.

The subgraphs further show a possible alternative mechanism of *CTNNB1* activation through upregulated *GPC3*, an early marker of HCC, as well as Wnt Family member 3a (*WNT3A*) and Frizzled 10 (*FZD10*). *WNT3A* promotes the stablization of *CTNNB1* and consequently expression of genes that are important for growth, proliferation and survival [79] through activity of transcription factor Lymphoid Enhancer-Binding Factor 1 (*LEF1*). As shown in the subgraph figure 7, *LEF1* ‘s known targets SRY-box 2 (*SOX2*) (Sex-Determining Region Y (*SRY*)) and Baculoviral IAP Repeat Containing 5 (*BIRC5*) are likely important contributers to *WNT* pathway driven *WNT* proliferation. *SOX2* is a pluripotency-associated transcription factor with known role in HCC development [80, 81, 82] and *BIRC5* (survinin) is an anti-apoptotic factor often implicated in chronic liver disease and liver cancer [83, 84, 85].

In summary, our algorithm reconstructed important components of the canonical *WNT* signaling pathway activation in liver cancer [86, 87, 88, 89, 90] from TCGA-LIHC RNA-Seq data and pairwise gene-gene interaction information from KEGG.

#### Crosstalk between integrin and WNT signaling

Another interesting pattern emerging in the upregulated subgraphs is the crosstalk between the *WNT* signaling cascade and integrin signaling. Over-expression of Secreted Phosphoprotein 1 (*SPP1*) has been shown to be a common feature for most known human malignancies and it is commonly associated with poor overall survival [91]. The binding of *SPP1* to integrins (e.g. integrin *α*V*β*3) leads to further activation of kinases associated with proliferation, epithelial-mesenchymal-transition, migration and invasion in HCC, such as Mitogen Activated Kinase-like Protein (*MAPK*), Phosphatidylinositol-4,5-bisphosphate 3-kinase (*PI3K*), Protein Tyrosine Kinase (*PTK2*), and SRC proto-oncogene/Non-receptor tyrosine kinase (*SRC*) [92]. Further captured by the subgraphs is that elevated expression of *PTK2* and *MAPK 12* are accompanied with elevated expression of cell cycle related genes (Cell Division Cycle 25 Homolog C / M-phase inducer phosphatase 1 (*CDC25C*), Cyclin-dependent Kinase 1 (*CDK1*) and Polo-like Kinase 1 (*PLK1*)), thus connecting over-expression of kinases with cell proliferation.

Although KEGG lists the interaction between *SRC* and *CTNNB1* as inhibitory in nature, other studies have concluded that activated Src enhances the accumulation of nuclear beta-catenin and therefore through their interaction contributes to an oncogenic phenotype [93].

In conclusion, the upregulated subgraphs capture the interaction of *SPP1* with integrin and consequent activation of *PTK2* and *SRC* together with their connection to the *WNT* signaling pathway (via *CTNNB1*) and cell cycle genes.

#### Downregulated oncogenes FOS and JUN and drug metabolism

The global downregulated subgraphs are centered around down-regulation of transcription factors *FOS* and *JUN*. The subgraph summary is depicted in figure 8. *FOS* and *JUN*, which form *AP-1* transcription complex are considered to be oncogenic factors and necessary for development of liver tumors [94]. Considering their prominent role in liver tumorigenesis, further experimental study of the significance of Jun and Fos downregulation on HCC development could be of great interest. Interestingly, RNA-seq data show that all *FOS* (*FOS*, *FOSB, FOSL1, FOSL2*) and *JUN* (*JUN, JUN B, JUN D*) isoforms are downregulated in a majority of liver tumors of the TCGA cohort (See Figure S11 in Additional File 2: Supplementary Figures).

**Figure 8:**
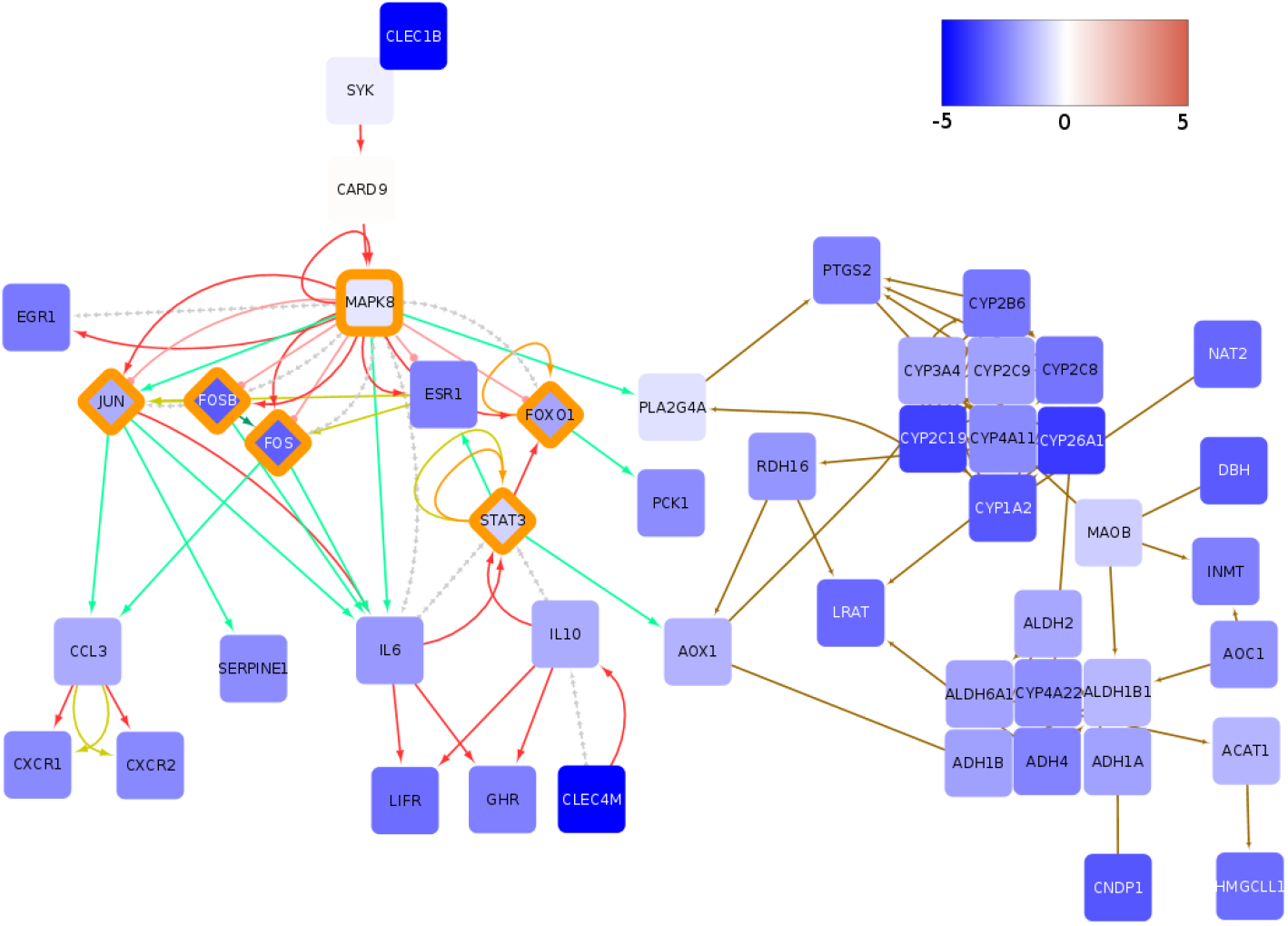
Global downregulated subgraph for TCGA-LIHC are centered on *FOS* and *JUN* transcription factors and drug metabolism. Color of nodes indicates the average log_2_ fold change of tumor samples compared to controls as represented by the color bar. The color of edges indicates the following interactions: activation (red), compound (brown), binding/association (yellow), indirect effect (dashed grey) and expression (green). Also noteworthy it the general connection of transcriptional activators and inhibitors to signaling as well as metabolic networks. Transcription regulators have been highlighted with an orange rim. See also supplementary figures S12-S16

Furthermore, the subgraphs show a number of downregulated Cytochrome P450 (*CYP*) enzymes as part of the most downregulated network of genes. *CYP 3A4* is mainly expressed in the liver and has an important role in the conversion of carcinogens, such as aflatoxin B_1_ toward their ultimate DNA-reactive metabolites [95], as well as, in detoxification of anticancer drugs [96]. Although the downregulation of *CYP* enzymes could potentially render HCC tumors sensitive to chemotherapy, liver tumors are notoriuosly irresponsive to chemotherapy [75]. Therefore, it is unclear how the gene pattern of *CYP* enzymes captured by the presented subgraphs could influence the HCC response to therapy and which compensatory mechanism is employed to counteract *CYP* downregulation.

### Personalized deregulated subgraphs for TCGA-LIHC

Finding deregulated subgraphs in a patient-resolved manner enables steps toward personalized medicine. In this section we introduce a case study where we employed our algorithm to find an upregulated subgraph for every TCGA-LIHC patient. Stratifying patients according to whether their subgraph contains a gene or not, one can identify genes whose inclusion into a patient’s inferred subgraph provides a survival handicap or advantage. Supplementary Figure S17 shows the survival effect for further identified *network-defined cancer genes.* Here, we concentrate on one particular such gene, namely Spleen Tyrosine Kinase (*SYK*).

#### Spleen tyrosine kinase (SYK) as a network-defined cancer gene

Patients whose subgraph contained the spleen tyrosine kinase (SYK) showed comparatively bad survival outlook (see Figures 9, 10).

**Figure 9:**
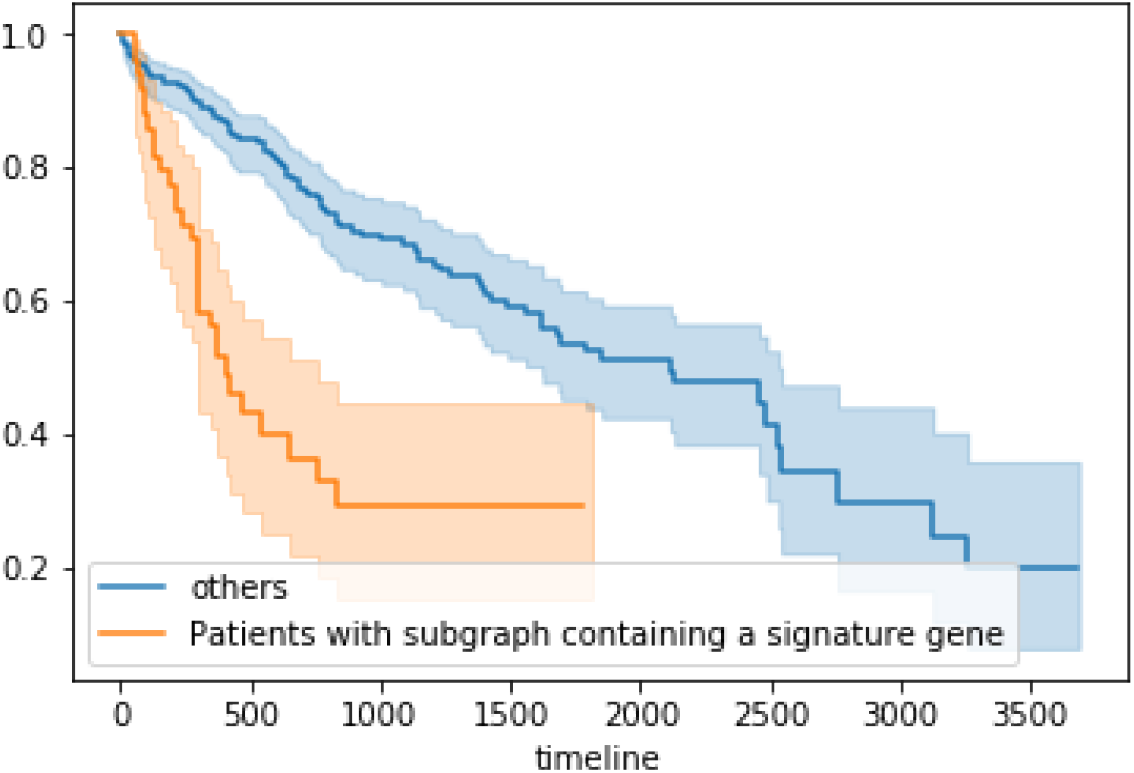
SYK signaling indicates poor survival. TCGA-LIHC cases TCGA-5C-AAPD, TCGA-CC-A3MA, TCGA-ED-A5KG, TCGA-DD-AACH, TCGA-YA-A8S7, TCGA-CC-5261, TCGA-CC-A3M9 show activated SYK signaling and poor survival. Survival difference is significant at p=0.0002 (Kaplan-Meier estimates and log-rank test).

**Figure 10:**
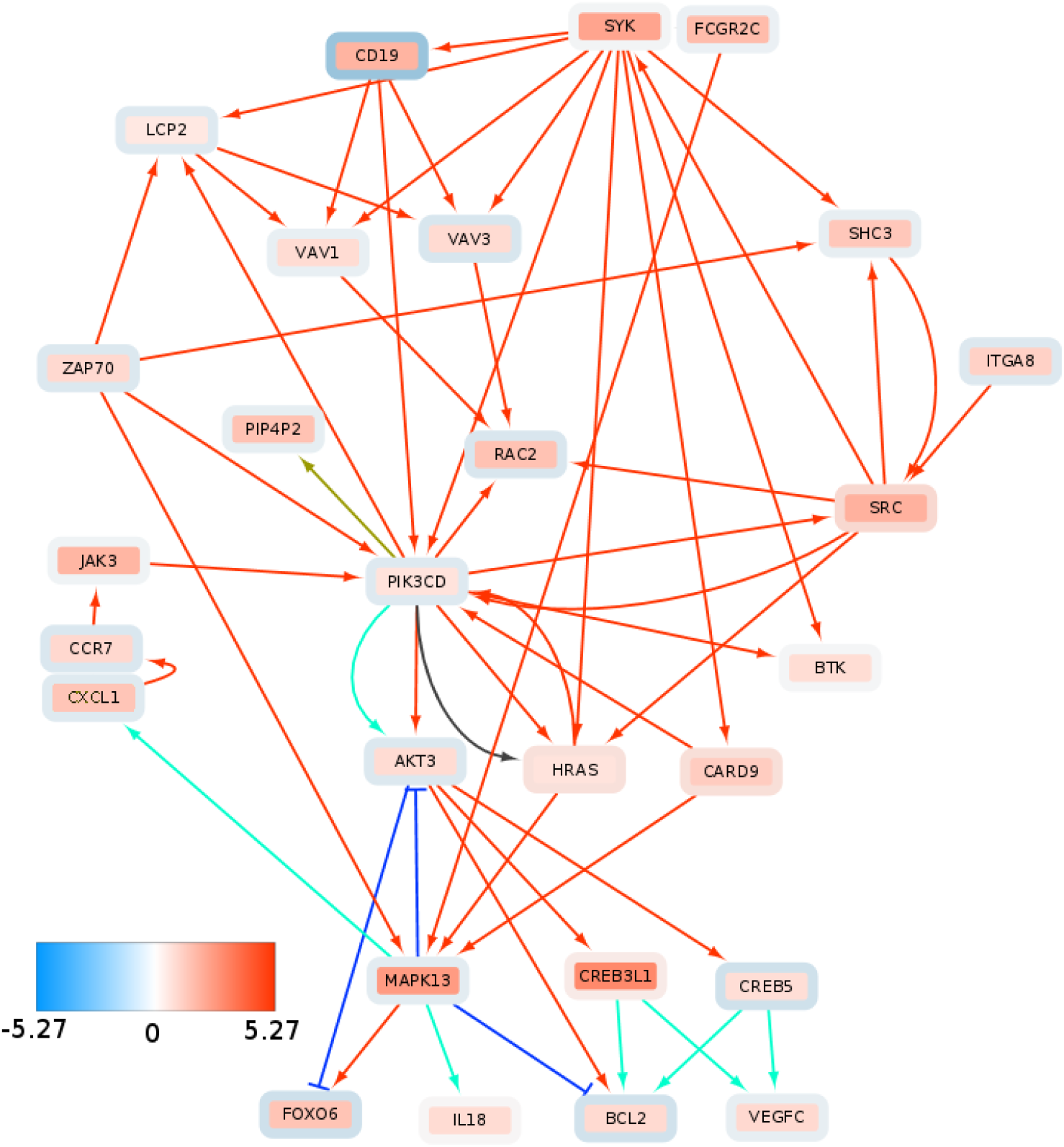
Consistent upregulation of SYK signaling components and downstream targets in subgraph of patients with poor survival. Inner node color represents the average log_2_ fold change across the “SYK-positive” patients and node rim color represent average log2 fold change across the rest of the TCGA-LIHC cohort. Color of edges indicates following interactions: activation (red), inhibition (dark blue), compound (brown), indirect effect (dark grey) and expression (blue green).

*SYK* is most commonly expressed in immune cells and its deregulation has been originally associated with hematopoietic cancers [97, 98, 99]. However, it has been shown that *SYK* plays a role in various other cancer types and its respective roles seem to vary significantly depending on the molecular (i.e. ultimately network) context [98]. *SYK* comes in the form of two splice variants, *SYK* (*L*) and *SYK* (*S*) [100]. In the context of liver cancer, *SYK* promoter hypermethylation and corresponding *SYK* downregulation has been associated with poor survival [101]. Furthermore, Checkpoint Kinase 1 (*CHK1*) mediated phosphorylation of *SYK* (*L*) and associated *SYK* degradation has been considered an oncogenic process [102], associating low levels of *SYK* as a factor of poor survival. On the other hand, [100] *SYK* (*S*) expression promotes metastasis development in HCC and thus leads to poor survival outcome. Furthermore, high *SYK* expression has been shown to promote liver fibrosis [103]. The development of HCC is closely related to formation and progression of fibrosis. Fibrosis represents excessive accumulation of extracellular matrix (ECM) and scarring tissue in an organ. A fibrotic environment promotes development of dysplastic nodules which can gradually progress to liver tumors [104]. In short, a somewhat inconsistent role of *SYK* as a tumor supressor or oncogene can be observed in many cancers [98], including liver cancer.

By employing DeRegNet, we identified by means of the approach defined as algorithm 2 a subgroup of HCC patients from the TCGA-LIHC cohort which show poor survival and a distinguished *SYK*-signaling pattern shown in Figure 10. The depicted network is manually extracted from the union graph of all the patient’s subgraphs which contained *SYK*. The network shows *SRC-SYK*-mediated activation of PI3K-Akt signaling via B-lymphocyte antigen CD19 (*CD19*) and Phosphatidylinositol 4,5-bisphosphate 3-kinase catalytic subunit delta (*PI3KCD*) (p110*δ*) [105]. Furthermore, *SYK* also feeds into mitogen-activated protein kinase 11-13 (p38) signaling (only *MAPK* 13 shown) through GTPase Hras (*HRAS*) and aspase recruitment domain-containing protein 9 (*CASP9*). p38 signaling promotes cytokine expression via Growth-regulated alphaprotein (*CXCL1*). Increased cytokine expression and activation is another canonical effect of *SYK* signaling [99]. This in turn, activates JAK signaling through Januskinase 3 (*JAK3*) activity, thereby reinforcing PI3K activation. Interestingly, *SYK* signaling is consistently linked to the upregulation of the guanine nucleotide exchange factors *VAV 1* and *VAV 3* [99, 97] (Guanine nucleotide exchange factor (*VAV*)). The proto-oncogene *VAV 3* is associated to adverse outcomes in colorectal [106] and breast cancer [107, 108]. Furthermore *VAV 3* mutations have been profiled to be potential drivers for liver cancer [109]. *VAV* signaling is mediated by forming a complex with Lymphocyte cytosolic protein 2 (*LCP2*) (SLP-76) upon activation of *SYK* signaling. *VAV*-meditated Ras-related C3 botulinum toxin substrate 2 (*RAC2*) activation may play a role in intravastation and motility [110]. Additionally, the subgraph shows upregulation of the B-cell lymphoma 2 (*BCL2*) gene, a known regulator of apoptosis [111], and vascular endothelial growth factor-C (*VEGGC*) which can promote metastasis [112] and angiogenesis [113, 114].

### Multi-omics subgraphs with consistent methylation and transcription patterns

To demonstrate a multi-omics application of DeRegNet (i.e. simultaneously using different omics layers) we have utilized RNA-seq and methylation data of the TCGA-LIHC cohort. With the transcriptome-methylome node scores defined in *Materials and Methods* we inferred a deregulated subgraph showing consistent patterns of methylation and transcription. In mammals, hyporegulation of the gene promoter typically leads to downregulation of gene expression and hypermethylation to upregulation of gene expression and hence the optimal subgraph we found represents a functional module which shows consistent patterns of gene regulation by means of promoter methylation [66]. The optimal subgraph depicted in Figure 11 is centered around protein kinase cAMP-activated catalytic subunit beta (PRKACB) gene. This gene is a catalytic subunit of cAMP (cyclic AMP)-dependent protein kinase. As such, it regulates signalling though cAMP. cAMP signaling is crucial to a large number of processes involved in carcinogenesis, including cell proliferation and differentiation [115]. As visible from the subgraph PRKACB gene is connected to a large number of downstream proteins, that could be potentially regulated through promoter methylation.

**Figure 11:**
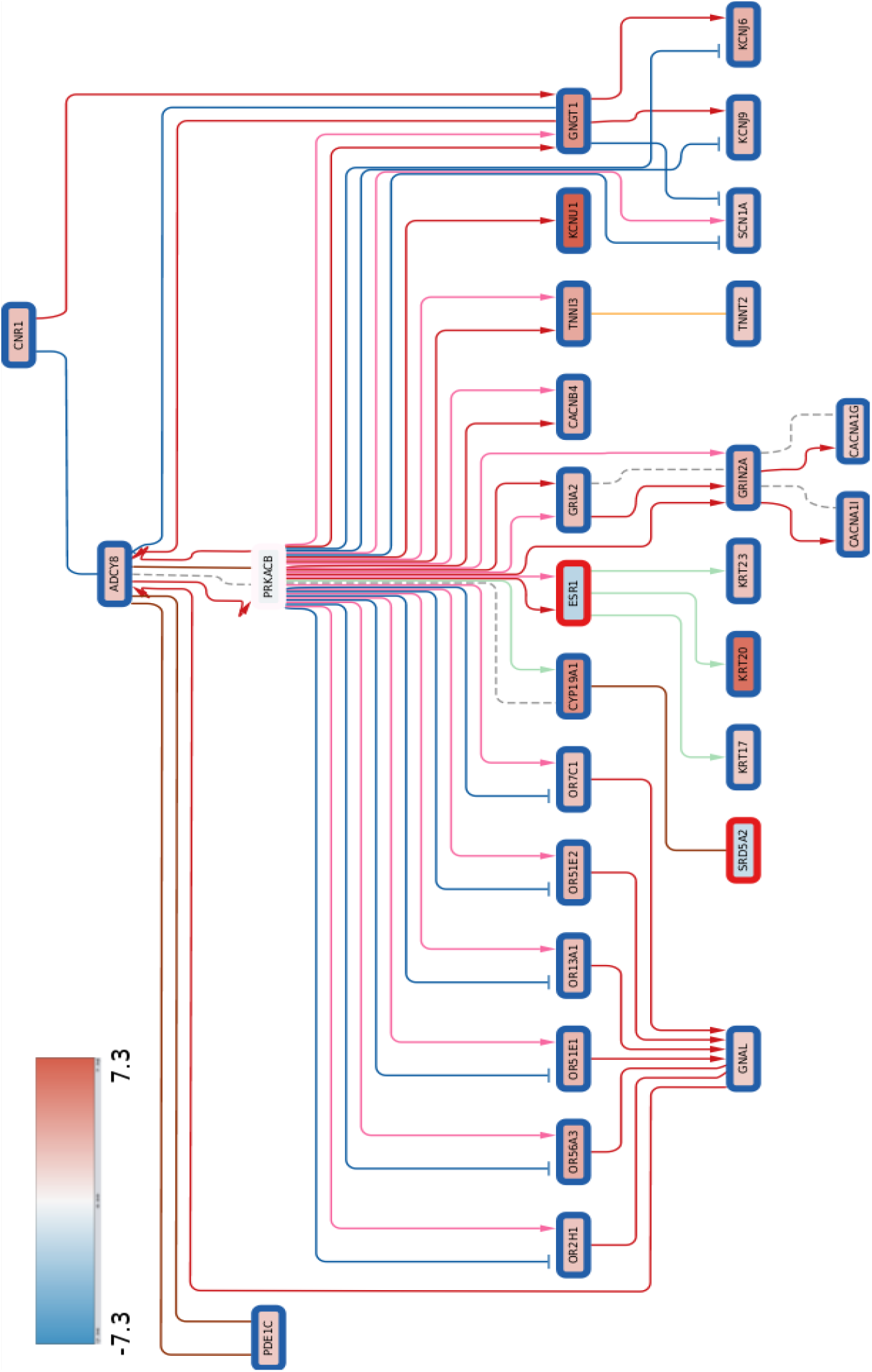
Multi-omics deregulated subgraph showing consistent methylation-transcription patterns. Inner node color represents RNA-Seq log2 fold change while the rim color of a node represents the methylation score *m_v_* where *red* corresponds to upregulated methylation and *blue* corresponds to downregulated methylation. The color of edges indicates the following interactions: activation (red), compound (brown), binding/association (yellow), indirect effect (dashed grey) and expression (green).

## Conclusion

We have shown DeRegNet’s capability to infer relevant patterns to a high degree of accuracy based on simulation benchmarks and showed that it compares favorably to related algorithms. Furthermore, application of DeRegNet to publically available data in a global fashion identified driving factors of liver cancer such as a transcriptionally activated WNT-pathway, thus showing that DeRegNet can provide valuable insight into a given omics experiment and may lead to novel and so far uncharacterized discoveries of gene/pathways involved in carcinogenesis and other biological contexts. An example of such discovery is the outlined insights into the global interaction of integrin and WNT signaling, as well as drug metabolism in liver cancer. In fact, profiling of such interaction between pathways is one of the main strengths of our algorithm over classical gene enrichment methods. Additionally, the application of our subgraph algorithm in a patient-specific manner could identify a consistent subgroup of patients showing poor prognosis potentially due to aberrant SYK signaling and therefore can generate meaningful hypotheses suitable for further experimental follow-up. Given that the SYK example is just one example case of a network-defined cancer gene, this indicates that DeRegNet is a useful hypothesis generation tool for network-guided personalized cancer research. In addition, further modes of application of the DeRegNet algorithm increase the spectrum of meaningful exploratory directions. Note, for example, that we only presented and discussed network-defined cancer genes (i.e. SYK in our subgraph example) for up-regulated subgraphs, while we have not presented the results of an analysis based on downregulated or generically deregulated (either up- or downregulated) subgraphs which would lead to similar opportunities. Another venue of further research is the utilization of deregulated subnetworks as features for phenotype prediction tasks. See *Additional File 1: Supplementary Material and Methods* for some computational experiments regarding survival predictions within the TCGA-LIHC cohort based on subnetwork features. Furthermore, DeRegNet promises to be usable in single cell data analysis. One example of such application can be a construction of cell-type specific subgraphs. For example, genes up- or downregulated in one cell type in comparison to other cell types can be used to define suitable node scores leading to identification of the most active subnetwork in a given cell type relative to other cell types. In conclusion, together with a solid underlying statistical model for which DeRegNet is shown to infer Maximum Likelihood estimates and its open-source implementation, this makes DeRegNet a viable option for any researcher interested in network interactions in an high-throughput omics context.

## Declarations

### Ethics approval and consent to participate

Not applicable.

### Consent for publication

Not applicable.

### Availability of data and materials

- Project name: DeRegNet
- Project home page: https://github.com/KohlbacherLab/deregnet
- Operating system: Linux, OSX and Windows via Docker
- Programming language: C++, Python
- Other requirements: Lemon Graph Library 1.3.1, Gurobi ≥ 8
- License: BSD-3-Clause
- Any restrictions to use by non-academics: Gurobi license required

Our implementation is written in C++ and Python and utilizes the Gurobi optmization libary (http://www.gurobi.com/index) and the Lemon graph library (https://lemon.cs.elte.hu/trac/lemon). Our software is open source under a BSD-3-Clause OSI-approved license and is available at https://github.com/KohlbacherLab/deregnet where you can also find installation instructions and usage examples. The algorithm can be run either by using a Python package or a command line tool via Docker images. The Docker images *sebwink/deregnet* are available at Docker Hub (https://hub.docker.com/r/sebwink/deregnet) and bundle all necessary dependencies. Additionaly Docker images are also provided via https://github.com/orgs/KohlbacherLab/packages?repo_name=deregnet. Furthermore, in order to run DeRegNet, a license for the Gurobi optimization library is required. For academic purposes these licences are readily obtained at https://www.gurobi.com/downloads/. The applications of DeRegNet to TCGA data appearing in this paper can be found at https://github.com/KohlbacherLab/deregnet-tcga. DeRegNet depends on a C++ library called *libgrbfrc* (https://github.com/KohlbacherLab/libgrbfrc) to solve fractional integer programs with Gurobi which was implemented by the authors of DeRegNet which is also available under the BSD-3-Clause open source license. Finally, to run the synthetic benchmarks presented in this paper, one can follow the instructions at https://github.com/KohlbacherLab/deregnet/tree/master/examples/custom-python-script. The benchmark code and results as obtained by the authors and presented in figure 4 are available here: https://github.com/KohlbacherLab/deregnet/tree/0.99.999/benchmark.

For more information, see Additional File 1: Supplementary Material and Methods.

### Authors’ contributions

SW designed, formalized and implemented the algorithms and overall research. SW, IW and MF visualized networks. IW, TT, OK contributed to the conceptual development of the presented methods. IW, SW and AN interpreted the resulting deregulated subnetworks biologically. OK and AN provided general supervision.

## Acknowledgements

We would like to thank all members of the Chair for Applied Bioinformatics at the University of Tubingen for valuable discussions and comments. Also we especially would like to thank Fabian Aicheler, Marc Rurik and Nico Weber for testing various components of the software at various stages and providing useful feedback. SW, IW, AN and OK acknowledge support from the *International Max Planck Research School (IMPRS) “From Molecules to Organisms”*.

## Abbreviations

**Acronyms**

AP-1: Activator protein 1. 19
BCL2: B-cell lymphoma 2. 22
BIRC5: Baculoviral IAP Repeat Containing 5. 18
CASP9: aspase recruitment domain-containing protein 9. 22
CD19: B-lymphocyte antigen CD19. 22
CDC25C: Cell Division Cycle 25 Homolog C / M-phase inducer phosphatase 1. 18
CDK1: Cyclin-dependent Kinase 1. 18
CHK1: Checkpoint Kinase 1. 21
CTNNB1: *β*-catenin. 17, 18
CXCL1: Growth-regulated alphaprotein. 22
CYP: Cytochrome P450. 19, 20
FOS: AP-1 transcription factor subunit / Fos proto-oncogene. 19
FZD10: Frizzled 10. 18
GPC3: Glypican-3. 17, 18
HRAS: GTPase Hras. 22
JAK3: Januskinase 3. 22
JUN: AP-1 transcription factor subunit / Jun proto-oncogene. 19
LCP2: Lymphocyte cytosolic protein 2. 22
LEF1: Lymphoid Enhancer-Binding Factor 1. 18
MAPK: Mitogen Activated Kinase-like Protein. 18, 22
PI3KCD: Phosphatidylinositol 4,5-bisphosphate 3-kinase catalytic subunit delta. 22
PI3K: Phosphatidylinositol-4,5-bisphosphate 3-kinase. 18
PLK1: Polo-like Kinase 1. 18
PTK2: Protein Tyrosine Kinase. 18
RAC2: Ras-related C3 botulinum toxin substrate 2. 22
SOX2: SRY-box 2. 18
SPP1: Secreted Phosphoprotein 1. 18
SRC: SRC proto-oncogene/Non-receptor tyrosine kinase. 18, 22
SRY: Sex-Determining Region Y. 18
SYK: Spleen Tyrosine Kinase. 20–22
TERT: Telomere Reverse Transcriptase. 17, 18
VAV: Guanine nucleotide exchange factor. 22
VEGGC: vascular endothelial growth factor-C. 22
WNT3A: Wnt Family member 3a. 18
WNT: Wingless-related integration site. 17, 18
DMP: differentially methylated probe. 12
ECM: Extracellular Matrix. 21
GSE: Gene Set Enrichment. 9, 11, 14, 16
HCC: Hepatocellular Carcinoma. 17–22
KEGG: Kyoto Encyclopedia of Genes and Genomes. 11, 18
LIHC: Liver Hepatocellular Carcinoma. 12, 13, 17–19, 22
RNA: Ribonucleic Acid. 11, 12, 18, 19, 23
SQN: Subset Quantile Normalization. 12
TCGA: The Cancer Genome Atlas. 11–13, 17–19, 22
w.r.t: with respect to. 13, 14

## Supplementary Material

### Additional Files

#### Additional File 1: Supplementary Material and Methods

Provides additional details and formalized exposition of many aspects of DeRegNet. In particular, it provides details on directions on how to run the DeRegNet software, definition and derivation of the probabilistic model underlying DeRegNet, as well as the proof that DeRegNet corresponds to maximum likelihood estimation under outlined model, DeRegNet in the context of the general optimization problem referred to as the *Maximum Average Weight Connected Subgraph Problem* and its relatives, proofs of certain structural properties of DeRegNet solutions, different application modes of the DeRegNet algorithms, fractional mixed-integer programming as it relates to the solution of DeRegNet instances, lazy constraints in branch-and-cut MILP solvers as it relates to DeRegNet, further solution technology employed for solving DeRegNet instances, DeRegNet benchmark simulations and use of DeRegNet subgraphs as a basis for feature engineering for survival prediction on the TCGA-LIHC dataset.

#### Additional File 2: Supplementary Figures

This document contains supplementary figures associated to the main text.

## Supplementary Material and Methods

### Abstract

This document contains details concerning the Material and Methods outlined in the main paper **de novo identification of maximally deregulated subnetworks based on multi-omics data with DeRegNet**. It provides details about the following topics:

- Directions on how to run the DeRegNet software
- Definition and derivation of the probabilistic model underlying DeRegNet, as well as the proof that DeRegNet corresponds to maximum likelihood estimation under outlined model
- DeRegNet in the context of the general optimization problem referred to as the *Maximum Average Weight Connected Subgraph Problem* and its relatives
- Proofs of certain structural properties of DeRegNet solutions
- Different application modes of the DeRegNet algorithms
- Fractional mixed-integer programming as it relates to the solution of DeRegNet instances
- Lazy constraints in branch-and-cut MILP solvers as it relates to DeRegNet
- Further solution technology employed for solving DeRegNet instances
- DeRegNet benchmark simulations
- Use of DeRegNet subgraphs as a basis for feature engineering for survival prediction on the TCGA-LIHC dataset

### How to use the DeRegNet software

#### DeRegNet Docker images and Gurobi setup

The main source code repository for DeRegNet is available here: https://github.com/sebwink/deregnet. DeRegNet is licensed under the BSD 3-clause OSI-approved open source license. The primary route to run DeRegNet is via Docker images which package DeRegNet and all its dependencies. Hence, in terms running DeRegNet on a Linux host there are only two dependencies: Docker and a Gurobi license. The official Docker images for DeRegNet can be found here: https://hub.docker.com/repository/docker/sebwink/deregnet. For instructions for setting up a Gurobi license for running DeRegNet it is referred to the source code repository where one can always find up-to-date information. The *sebwink/deregnet* Docker images support basically two modes of usage: command-line and Python package.

#### Running DeRegNet via Docker

Assuming you have Docker and a named-user Gurobi license configured, running DeRegNet in command-line mode is as easy as running

~~~
git clone https://github.com/sebwink/deregnet && **cd** deregnet
docker/named-user/run sebwink/deregnet: 0.99.999 avgdrgnt.py **—help**
~~~

which would display all available command-line options for avgdrgnt.py (i.e. DeRegNet’s main script):

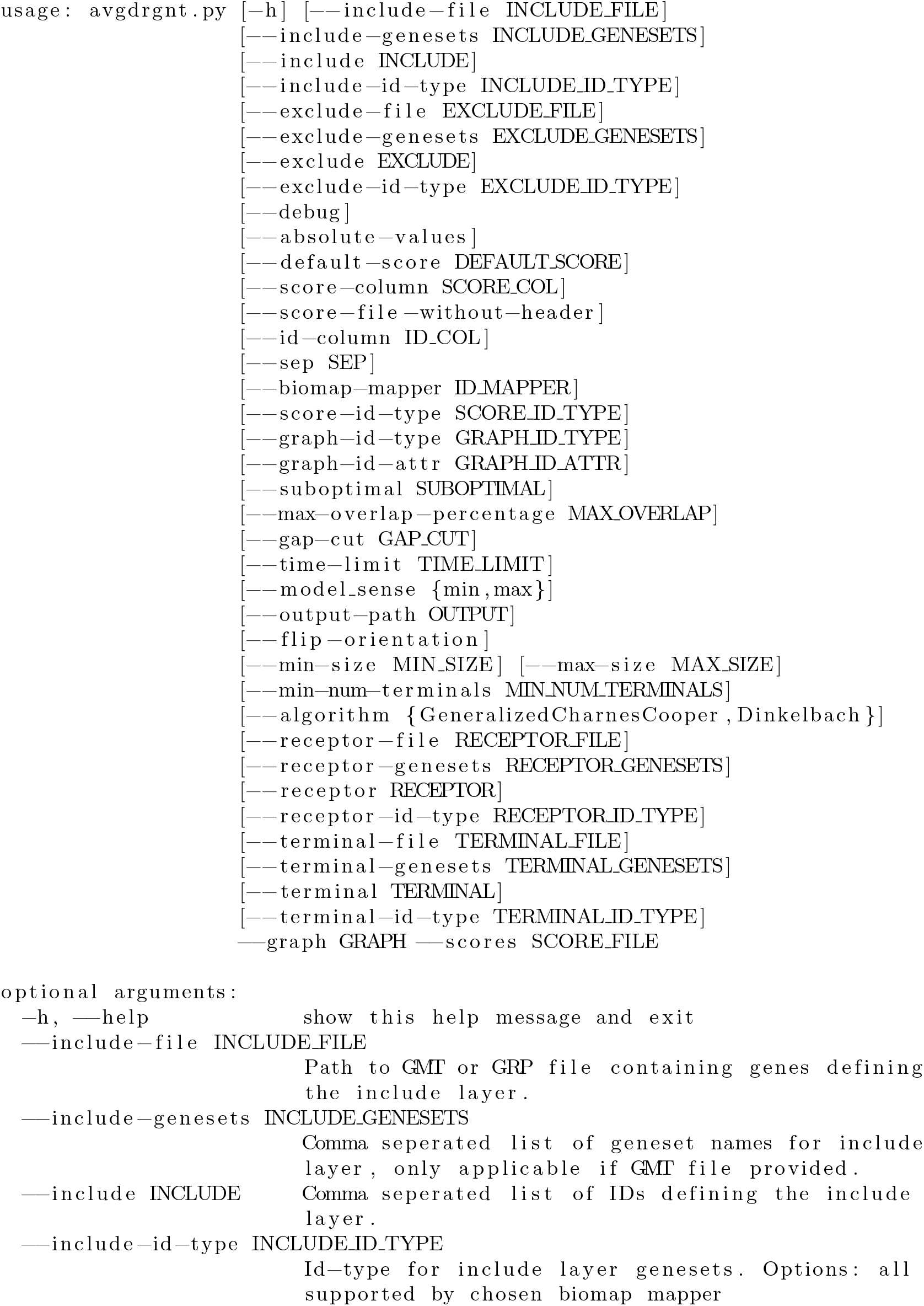
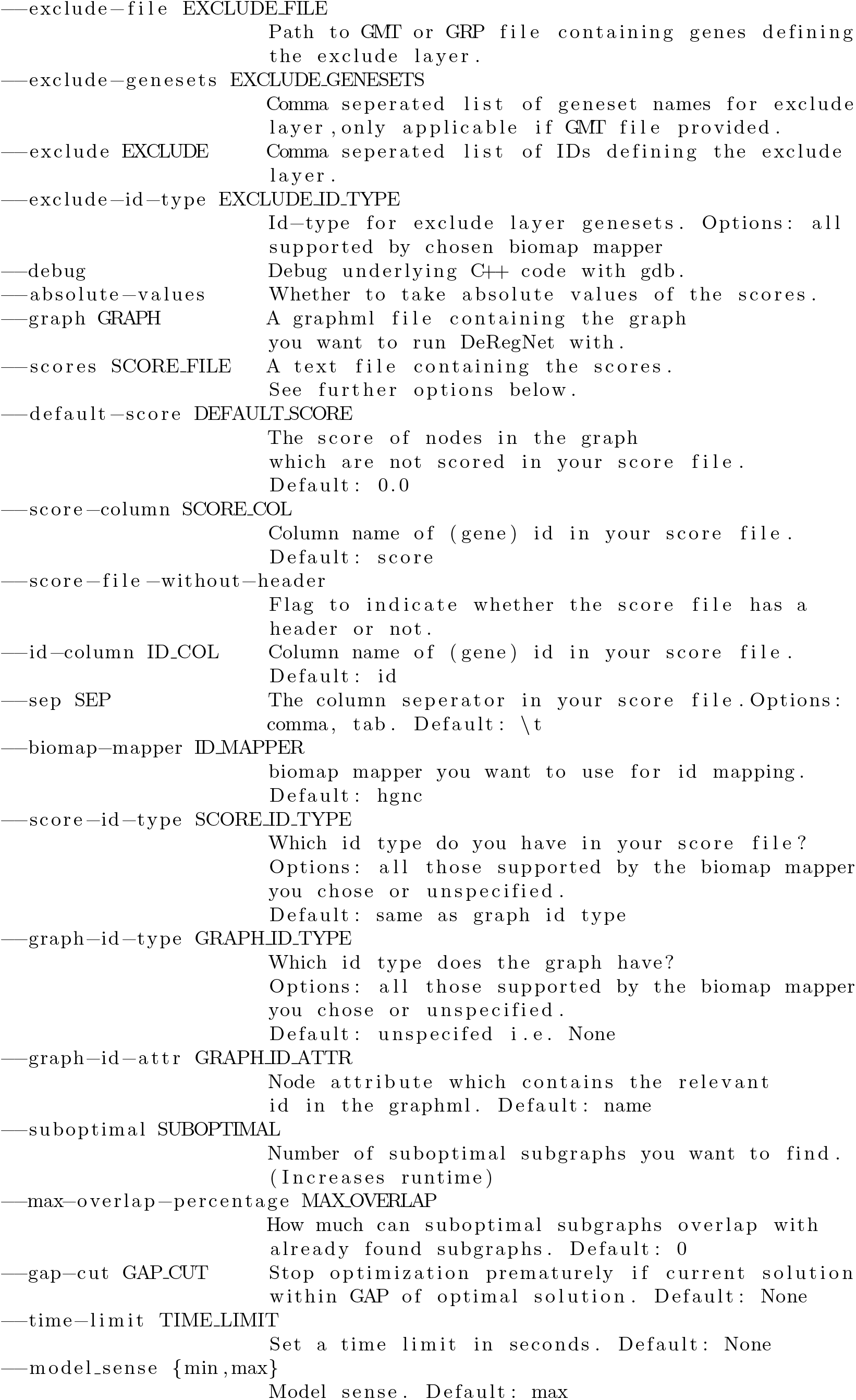
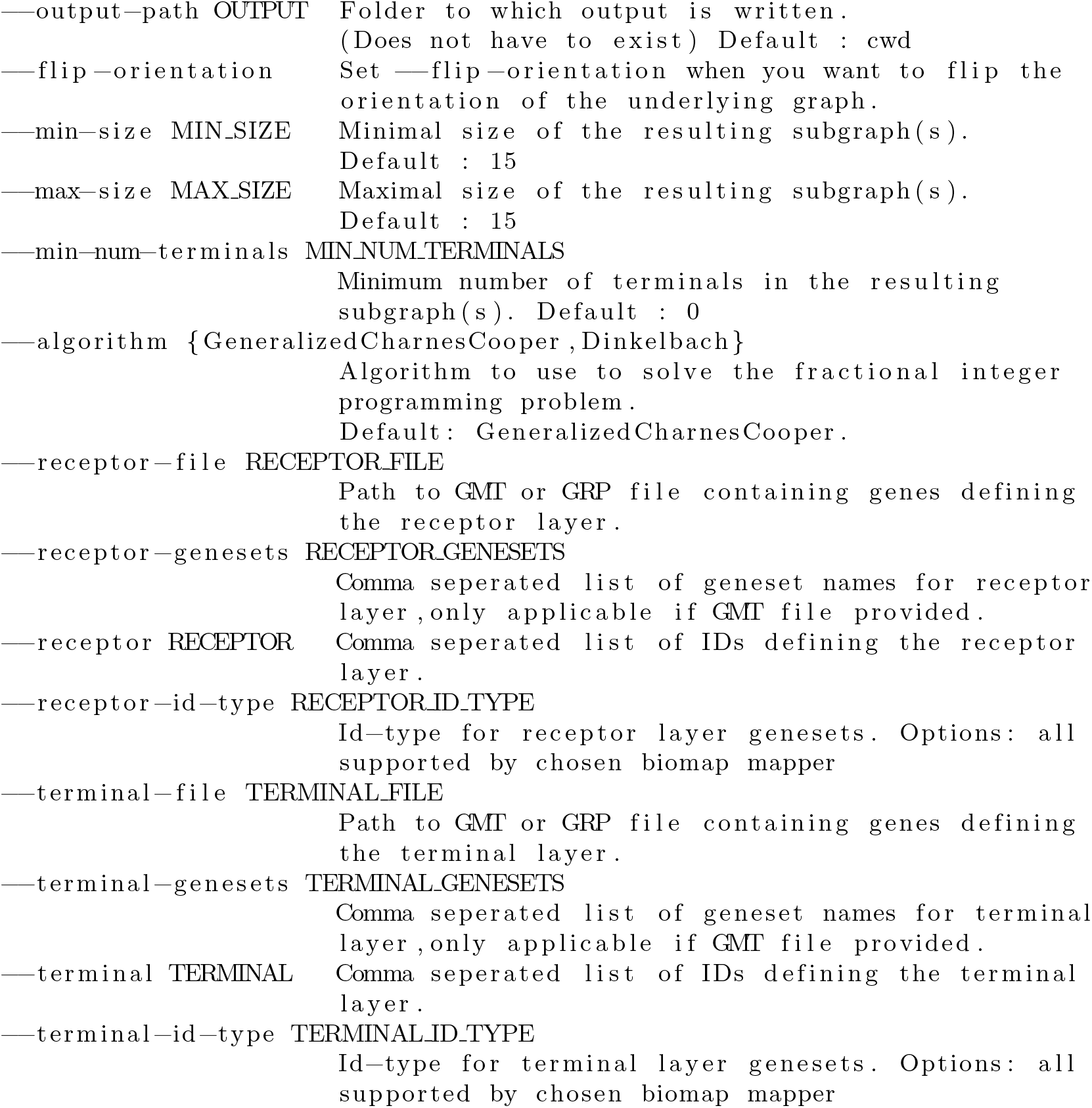

Still in the top-level of the repository, you can find your first subgraph like so:

~~~
docker/named—user/run sebwink/deregnet:latest avgdrgnt.py \
  ——graph **test**/kegg_hsa.graphml \
  ——scores **test**/data/score.csv \
  ——sep, \
  ——graph–id–attr ensembl
~~~

This will generate *deregnet.log* and finally *optimal.graphml* where the former is a log of the optimization procedure carried out by DeRegNet and the latter the resulting optimal subgraph in GraphML format. For more information on GraphML, the most prominent graph serialization format supported by DeRegNet, see: http://graphml.graphdrawing.org/.

#### DeRegNet Python package via Docker

For more custom analyses it is often necessary to work with the deregnet Python package directly. This is also supported by the *sebwink/deregnet* Docker images which come with all the relevant packages pre-installed and properly configured. E.g. in order to run the benchmarks presented in the main text, you can follow the directions given here: https://github.com/sebwink/deregnet/tree/master/examples/custom-python-script. Running any Python script which uses the *deregnet* Python package is then as easy as:

~~~
docker/named–user/run sebwink/deregnet: 0.99.999 python3 any_script.py
~~~

### Results concerning the probabilistic model for DeRegNet

This subsection formalizes the notion that a *deregulated* subgraph satisfying given topological constraints should have higher/maximal probability of deregulation with respect to all possible subgraphs of that particular topological class. We present a basic probabilistic model yielding one possible formal probabilistic rationale for optimizing a model of form given in the main paper. Furthermore we provide a suitable interpretation of the model proposed in [1] in terms of that model, showing that DeRegNet solves a more general problem in the statistical sense necessitated by the probabilistic model introduced in the main text.

For sake of locality of exposition we restate the statistical model as introduced in the main text. The model assumes binary node scores *s*: *V* → {0,1} which are realizations of random variables **S** = (*S_v_*)_*v*∈*V*_. Further it is assumed the existence of a subset of vertices *V*′ ⊂ *V* such that *S_v_* |*v* ∈ *V*′ ~ *Ber*(*p*′) and *S_v_*|*v* ∈ *V*\*V*′ ~ *Ber*(*p*) with *p,p*′ ∈ (0,1) denoting probabilites of deregulation outside and inside of the deregulated subgraph respectively. It is assumed that *p*′ > *p* to reflect the idea of *higher* deregulation (probability) in the *deregulated* subgraph. The network context (dependency) is introduced via the restriction that 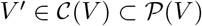. Here, 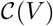 denotes the set of feasible substructures and should (can) reflect topologies inspired by known biomolecular pathway topologies like the one described in [1] and the last subsection. Furthermore it is assumed, that the (*S_v_*), given a network context and deregulation probabilities *p,p*′, are independent. We further introduce the notation 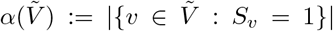 and considering *V*′,*p,p*′ to be parameters, and a subgraph determined by indicator variables *x* as outlined in the previous subsection, we can state:

#### Proposition 1

*The log-likelihood* 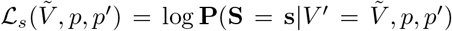 *under above model is given by*:

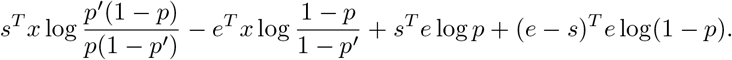

*Proof*

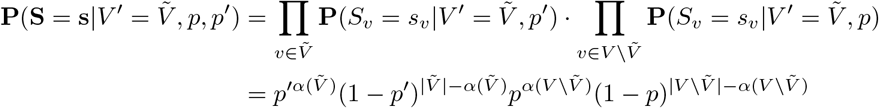

Employing decision variables 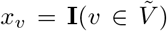, we can write 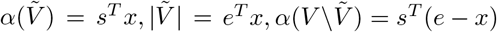. and 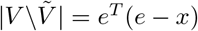. It follows that the log-likelihood 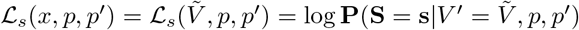 can be written as:

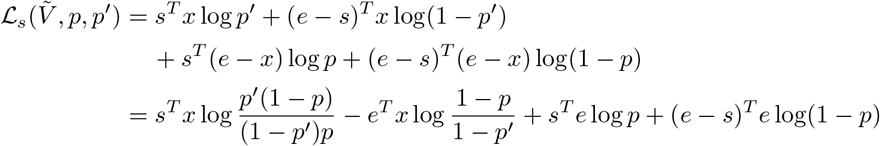

We call an optimization model maximizing the objective *s*^T^*x* subject to any constraints on x (the subgraph topology) a *model of Backes-type* [1]. Note that the DeRegNet model reduces to a Backes-type model in case of *k_min_* = *k_max_*.

#### Proposition 2

*Any subgraph model of Backes-type enforcing a fixed subgraph size can be interpreted as maximum likelihood estimation with respect to subgraph structure given the above model.*

*Proof* Given the log-likelihood as determined by proposition 1, ignoring the constant term with respect to x, a maximum likelihood estimator V* with respect to subgraph structure can be determined as follows:

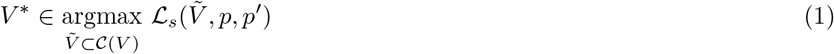

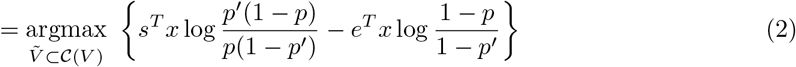

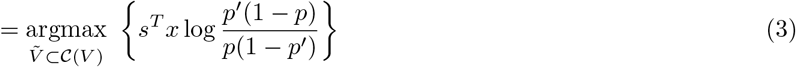

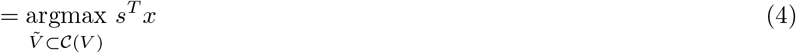

Here, equality (2.4) follows from the assumption that the topological constraints of the optimization model enforce a constant subgraphs size (i.e. *e^T^x* = *k* for some fixed 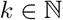). The last equality follows (by assumption *p*′ > *p*) because 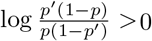. Overall, a maximum likelihood estimator is given by a solution to a given Backes-type optimization model max *s^T^x* with subgraph topology restricted to subgraphs from 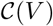.

In particular, the specific model proposed by [1] lends itself to the just justified interpretation:

#### Corollary 1

*The optimization model suggested by [1] can be interpreted as maximum likelihood estimation with respect to subgraph structure given the above probabilistic model.*

We now proceed to provide a maximum likelihood interpretation for the DeRegNet model. Since the DeRegNet model does not assume a fixed subgraph size, above conclusions do not apply. Under the assumption that the parameter *p* is estimated external to the model and represents some general base level of deregulation one can by (conceptual) reduction from the full log-likelihood 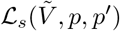 to 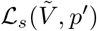 state the following proposition.

#### Proposition 3

*Solving a DeRegNet instance amounts to maximum likelihood estimation under above model with respect to subgraph structure and deregulation probability p′ (assuming p′ > 0*).

*Proof* Given the log-likelihood as in proposition 1, one can differentiate with respect to *p*′:

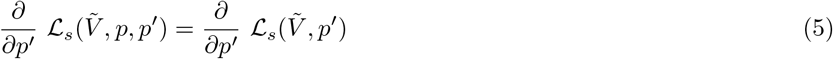

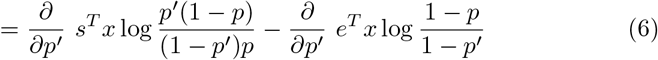

By computing

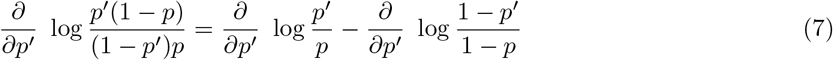

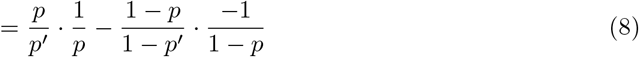

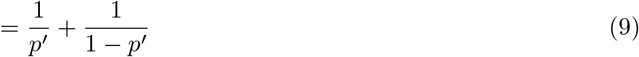

and

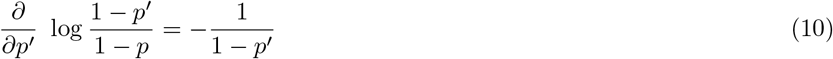

one obtains

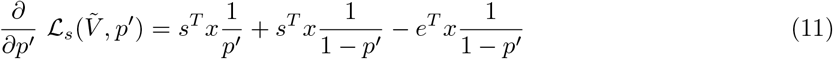

Requiring 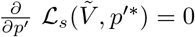 and with

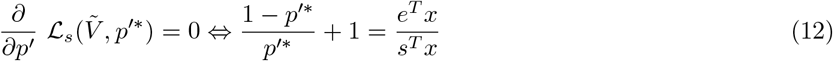

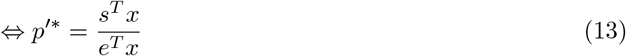

and since *s^T^x* ≤ *e^T^x* and *s^T^x* > 0 under the assumption that there is at least one node deregulated in the found subgraph and *p*′ > 0:

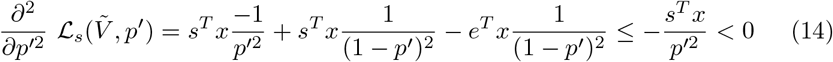

one arrives at

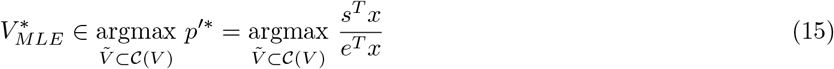

since no terms involving *x* were dropped in the derivation for *p*′*.

The propositions of this subsection show, that, given the introduced statistical model, solving a DeRegNet instance instead of an instance of the optimization model proposed in [1] allows to carry out maximum likelihood estimation without the need to fix the subgraph size in advance. Given the assumptions of the model, these results hold regardless of further topological constraints and only relate to the respective objective functions.

### Maximum Average Weight Connected Subgraph Problems

#### (Rooted) Maximum (Average) Weight Connected Subgraph Problems

In terms of mathematical optimization and up to minor modifications, for example the requirement of the subgraphs to be of a certain predefined size 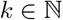, [1] solve instances of the so called (Rooted) Maximum Weight Connected Subgraph Problem.

#### Definition 1

(Maximum Weight Connected Subgraph Problem (MWCSP)) *Given a directed graph G* = (*V, E*) *and node scores s*: *V* → ℝ, *find a set of nodes V*′ ⊂ *V whose induced subgraph* (*V*′,*E*′) *maximizes* 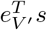 *such that there is a node r* ∈ *V*′ *such that there is a directed path from r to every other node v* ∈ *V*′.

By fixing the root node in the MWCSP to a particular node in the underlying graph one arrvies at the so called **Rooted Maximum Weight Connected Subgraph Problem (RMWCSP)**:

#### Definition 2

(Rooted Maximum Weight Connected Subgraph Problem (RMWCSP)) *Given a directed graph G* = (*V, E*), *node scores s*: *V* → ℝ, *and a node r* ∈ *V called the root node, find a set of nodes V*′ ⊂ *V with r* ∈ *V′ whose induced subgraph* (*V*′, *E*′) *maximizes 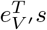 such that there is a directed path from r to every other node v* ∈ *V*′.

The (R)MWCSP has found applications in network biology [2], [1]. It also attracted general computational and theoretical research in recent years [3], [4], from different integer programming formulations and problem-specific branch-and-cut strategies [5], [6], [7], [8], to more recent research on computational strategies for addressing large-scale instances [9] and problem reduction techniques and heuristics [10], [11].

#### The Maximum Average Weight Connected Subgraph Problem (MAWCSP)

Analogously to the (R)MWCSP one can define versions which strive to optimize the average score in the subgraph.

#### Definition 3

(Maximum Average Weight Connected Subgraph Problem (MAWCSP)) *Given a directed graph G* = (*V, E*) *and node scores s*: *V* → ℝ, *find a set of nodes V*′ ⊂ *V whose induced subgraph* (*V*′, *E*′) *maximizes* 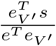 *such that there is a node r* ∈ *V*′ *such that there is a directed path from r to every other node v* ∈ *V*′.

#### Definition 4

(Rooted Maximum Average Weight Connected Subgraph Problem (RMAWCSP))

*Given a directed graph G* = (*V, E*), *node scores s*: *V* → ℝ, *and a node r* ∈ *V called the root node, find a set of nodes V*’ ⊂ *V with r* ∈ *V*′ *whose induced subgraph* (*V*′, *E*′) *maximizes* 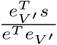 *such that there is a directed path from r to every other node v ∈ V*′.

DeRegNet solves extended versions of the (Rooted) Maximum Average Weight Connected Subgraph Problem.

### Some formal properties of DeRegNet solutions

In terms of the notation and exact formulation provided in the main text, we will here formally specify certain topological characteristics of solutions of the above model which were hinted at before. For similar proofs and also alternative formulations for the MWCSP it is referred to [1], [7], [6], [5], [8]. I first formally recapture the defining topological feature of problems of (R)M(A)WCS flavour for DeRegNet.

#### Proposition 4

*A feasible subgraph V* of a DeRegNet instance has the property that any node in the subgraph can be reached from the root of the subgraph.*

*Proof* Any given node *v* ∈ *V*\* of the subgraph is contained in a strongly connected component. By constraints (2.1e) and (2.1f) this strongly connected component either contains the root node or is reachable from some node *u* ∈ *V\** in the subgraph which is not in that strongly connected component: Let *S* ⊂ *V* be the vertex set inducing the strongly connected component. If the root is not in *S* we have 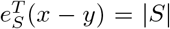 and hence it need to hold 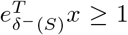, otherwise one would have 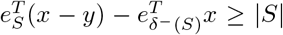 in violation of constraints (2.1e) and (2.1f). If the root node is in *S*, it holds that 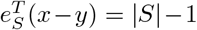 and hence constraints (2.1e) and (2.1f) always hold due to 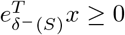. In the case, that the root node is in *V*′s component, *v* is reachable from the root node. In the case the component does not contain the root, repeat the argument with *u* instead of *v*. Again, the root is in the strongly connected component of *u* or the component is reachable from some *u*′ ∈ *V*\*, and so on. Since the subgraph has a finite number of strongly connected components, one ultimately will encounter the component containing the root in the above argument which proves the the existence of a path to any arbitrary *v* ∈ *V*\* from the root node.

The terminals from the terminal set *T* represent terminals of a subgraph in the following sense.

#### Proposition 5

*A feasible subgraph V* of a DeRegNet instance has the property that a node v* ∈ *V* in the subgraph with v* ∉ *T has to have an outgoing edge into the subgraph, i.e. only terminal nodes are allowed to have no outgoing edges within the subgraph.*

*Proof* Given a non-terminal node *v* ∉ *T* one has constraint (2.1h): 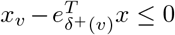, i.e. if *x_v_* = 1 it has to hold that 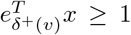. The latter inequality means that there exists another node *u* ∈ *V*\* such that (*v,u*) ∈ *E, E* being the edge set of the underlying graph.

### Further application modes of DeRegNet

#### Fixing the root node

Instead of the *root* being determined by the algorithm as outlined in the previous paragraph, one can also specify a given node *r* ∈ *V* as root [1]. In this case, one does not need the *y* variables anymore and, since the constraint logic can be carried over analogously, we can write the corresponding fractional integer problem as:

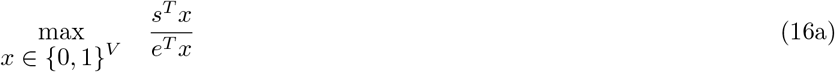

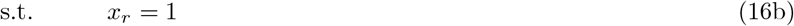

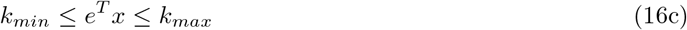

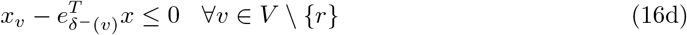

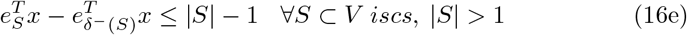

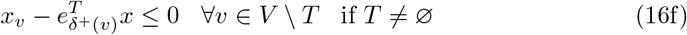

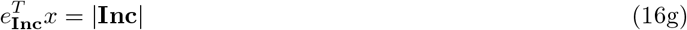

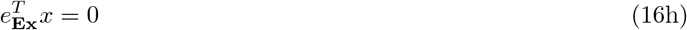

Note, that the above formulation is a special case of the more general formulation of the previous section, namely *R* = {*r*}. It is nonetheless convenient to sometimes refer to the tuple (*G, r, T,* **Ex**, **Inc**, *s*) as a *rooted DeRegNet instance*. All other terminology from the general case carries over without modification.

#### Reversing the orientation

The default version of the just outlined algorithm will find subnetworks which possess a ”root” node from which one can reach any other node in the subnetwork. This can be interpreted as the subnetwork being deregulated downstream of that root. As outlined in the previous sections, this root can either be determined by the algorithm or pre-determined by biological curiosity or insight. By reversing the orientation of the graph one can easily obtain subnetworks where the ”root” can be reached from any node in the subnetwork. Such a subgraph can be interpreted as deregulated upstream of the either algorithmically determined or user-defined ”root” node. In that case a more intuitive name for the ”root” is ”terminal” or ”destination”. Formally this difference in the structure of the output can be achieved by substituting the original graph *G* with the transposed graph 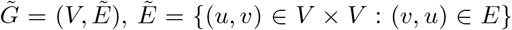, and defining the models as before with the roles of receptors and terminals exchanged.

#### Definition 5

*A **reverse solution** of a DeRegNet instance I* = (*G, R, T, **Ex**, **Inc**, s*) *with underlying graph G* = (*V, E*) *is the (graph) transpose of an optimal subgraph of the DeRegNet instance* 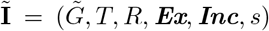. *The latter is called the reverse instance of* **I**. *Here*, 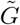 *denotes the transposed graph of G, i.e*. 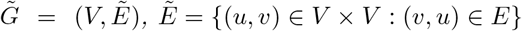.

After the algorithm found subnetworks with respect to the reversed graph the resulting subnetworks have to be re-reversed to reflect physical reality. Also note, that the reversed instance exchanges the roles of receptors and terminal nodes to keep the intuitive notions associated with these terms in line with the topology of the just defined reverse solutions.

#### Extracting suboptimal subnetworks

Although the strategy to optimize seems like a sensible heuristic, it is nonetheless just an heuristic. There is no intrinsic need for a biological system at hand to behave consistently with this optimization objective in the sense that it is not granted that the patterns found by the algorithm actually correspond to what is biologically important in the given situation. Vice versa, something (nodes, a particular pattern of nodes) not showing up in any subgraph does not mean that they may not be important in the given context. While this cannot be mediated completely, it is sensible to find at least possible suboptimal patterns along with the optimal one. This can be seen as a step to capture mathematically speaking slightly less optimal but biologically potentially similarily or even more important patterns. I implement this notion by following the appproach found in [2] and adapt it to DeRegNet. Given a specified *maximal overlap α* ∈ [0,1) and a (induced) subgraph *V*\* ⊂ *V* one adds to the DeRegNet model as stated in the main part of the paper the suboptimality constraint 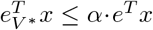 and reoptimizes, forcing any corresponding subgraph to be found to maximally have 100 ± *α* % node overlap with the the nodes of the previously found subgraph. One can iterate this theme. For example, given a set of subgraphs *V*^(1)^,…, *V*^(*k*)^ for some 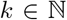 one can add the constraints 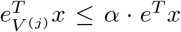 for all *j* = 1,…, *k* to the DeRegNet instance to obtain a optimal subgraph of that modified DeRegNet instance which is guaranteed to have node overlap ≤ *α* with any of the *V*^(*j*)^. With *V*^(1)^ = *V*\* being the original optimal subgraph of a DeRegNet instance one thus obtains a series of suboptimal subgraphs *V*^(2)^,…, *V*^(*k*)^. The question which *k* to choose can be for example decided such that one chooses the *k* for which 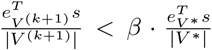 for the first time for some *β* ∈ [0,1]. Here, *β* quantifies the degree of suboptimality one is willing to accept.

### Fractional mixed-integer programming

#### Definition 6

(Fractional mixed-integer linear program; FMILP)

*A **Fractional mixed-integer linear program (FMILP)** is an optimization problem of the following structure*:

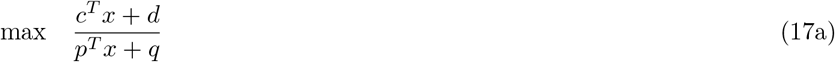

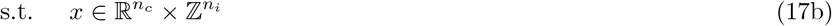

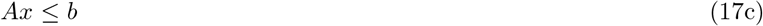

*Here, c, p* ∈ ℝ^*n*^, *d,q* ∈ ℝ *define the objective, A* ∈ ℝ^*m*×*n*^, *b* ∈ ℝ^*m*^ *define* 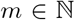 *linear constraints and* 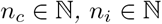 *denote the number of continuous and discrete (integer) variables*.

We assume 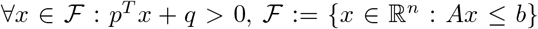. Fractional mixed-integer linear problems are hence mixed-integer problems except for the objective which is a rational function with linear enumerator and denominator instead. While a FMILP is non-convex, it turns out that a FMILP is pseudolinear and hence quasilinear, rendering local optima to be globally optimal [12].

#### Proposition 6

*A FMILP is pseudoconvex and pseudoconcave.*

#### Proposition 7

*A FMILP is strictly quasiconvex and strictly quasiconcave.*

#### Proposition 8

*A local optimum of a FMILP is also a global optimum.*

The latter facts render FMILP solvable by any generic mixed-integer nonlinear programming (MINLP) solver which can handle pseudolinear objective functions [12]. Empirically, it was shown that iterative schemes [12] or linearization-reformulation approaches [13] outperform generic MINLP solvers with respect to computing time and memory footprint. These approaches rely on a mixed-integer linear programming (MILP) solver as their optimization kernel, hence unlocking the power of modern MILP software, and rely on transforming the original problem into a (sequence of) MILP problem(s). The DeRegNet software package discussed in the main text implements a Dinkelbach-type algorithm [12] and a reformulation-linearization method [13] resembling the Charnes-Cooper method [14] for solving fractional linear programs (FLP). The following sections provide algorithmic details on the these methods.

#### Dinkelbach-type algorithm (Dinkelbach algorithm)

Originating in the 1960’s [15, 16] and studied in the context of FMILP problems [17, 12] later on, the Dinkelbach algorithm relies on the iterative solution of linear problems only containing the original variables and an auxiliary iteration parameter. *Algorithm 1* details the procedure. In the following, as well as in the entire thesis, *Dinkelbach algorithm* and *Dinkelbach-type algorithm* are used synonymously to refer to *Algorithm 1*.

**Algorithm 1:**
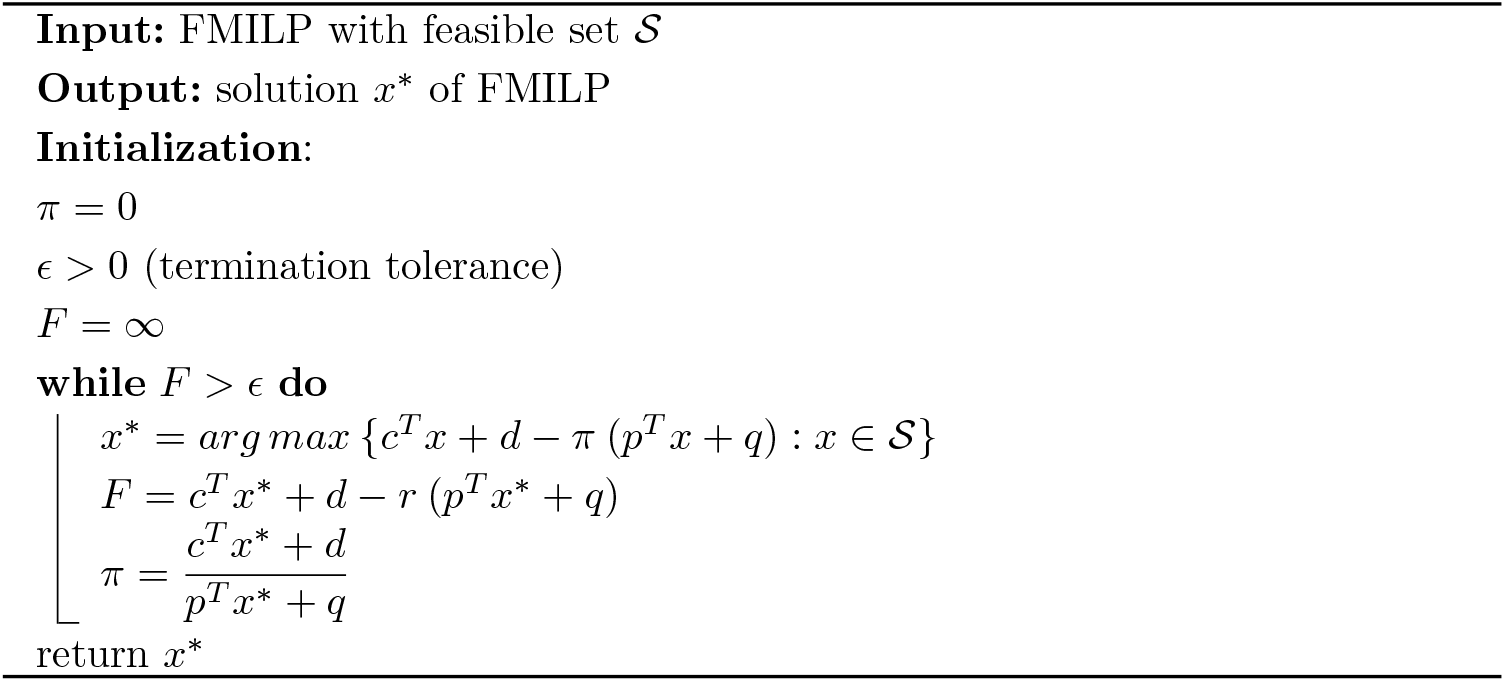
Dinkelbach-type algorithm

The mixed-integer linear program appearing in the *while*-loop of *algorithm 1* is called a *Dinkelbach iteration problem.* Dinkelbach’s algorithm iteratively solves a sequence Dinkelbach iteration problems until some convergence criterion is met. The follwing subsection shows that this procedure indeed solves the original FMILP.

##### Correctness of Dinkelbach’s Algorithm (1) - based on You et al.[12]

In order to facilitate the following exposition the functions 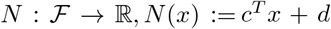 for the nominator and 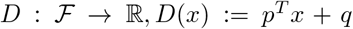 for the denominator of the objective function are introduced. Without loss of generality one can set *d* = *q* = 0 since one can introduce dummy variables *x_d_* and *x_q_* with linear constraints *x_d_* = *x_q_* = 1 and corresponding coefficients *c_d_* = *p_q_* = 1 leading to *N*(*x*) = *c^T^x* + *c_d_x_d_* and *D*(*x*) = *p^T^x* + *p_q_x_q_*. Furthermore, define *L_π_*(*x*):= *N*(*x*) – *πD*(*x*) and *F*: ℝ → ℝ, 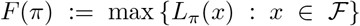 be the optimal objective value of a Dinkelbach iteration problem as a function of the auxiliary parameter *π*. Without loss of generality we assume *D*(*x*) > 0 for all 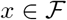.

The two main results concerning Dinkelbach’s algorithm are the following:

#### Proposition 9

(Optimality criterion, [13] Proposition 1)

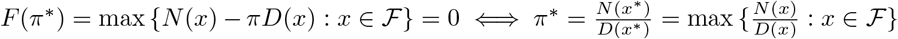 *where* 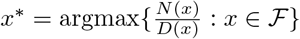

#### Proposition 10

(Convergence (rate), [13] Proposition 2)

*Dinkelbach’s algorithm converges superlinearly to π*\* *in where* 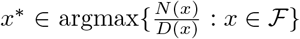 *and* 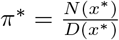.

We follow [13] in proving the above propositions via a series of lemmas.

#### Lemma 1

([13] Appendix, Lemma 4)

*F is convex*.

*Proof* For λ ∈ [0,1], let 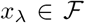 be 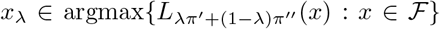 with *π*′,π” ∈ ℝ. Then:

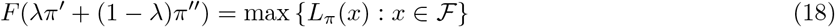

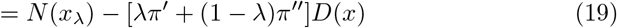

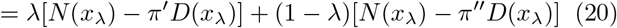

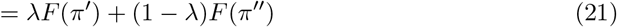

#### Lemma 2

([13] Appendix, Lemma 5)

*F is strictly monotonically increasing, i.e. π*′ < *π*″ ⇒ *F*(*π*′) < *F*(*π*″).

*Proof* Given *π*′ < *π*” one obtains with 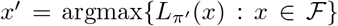 and 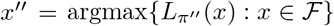:

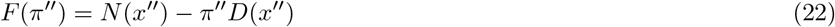

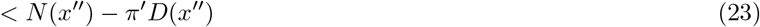

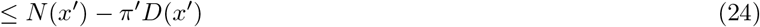

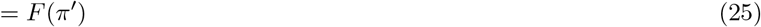

#### Lemma 3

([13] Appendix, Lemma 6)

*F*(*π*) = 0 *has a unique solution*.

*Proof* Follows from 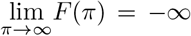 and 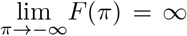 and *F* being strictly monotonically increasing (Lemma 2).

#### Lemma 4

([13] Appendix, Lemma 7)

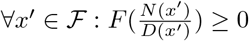

*Proof* For any 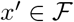 one has:

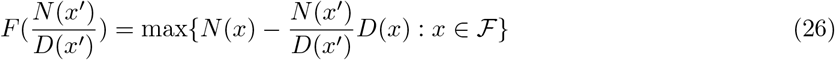

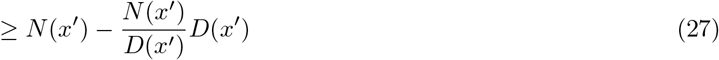

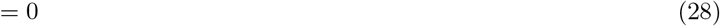

One can now prove proposition 1:

#### Proof of proposition 1

We have to show: 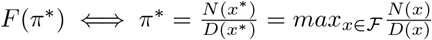.

⇒: Given 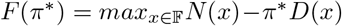 it follows with 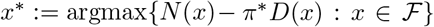 for all 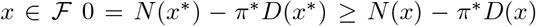. Hence 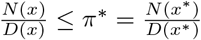, i.e. 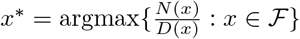.

⇐: With 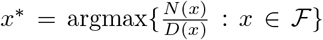 one has 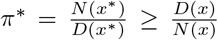. Under our general assumption *D*(*x*) > 0 for all 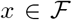 it follows *N*(*x*) – *π*\**D*(*x*) ≤ 0 = *N*(*x*\*) – *π*\**D*(*x*\*) for all 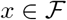 which shows 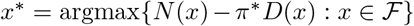.

From now onward, let *π*\* be the unique solution of *F*(*π*) = 0 and let 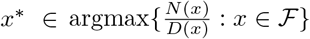 with 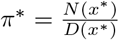.

#### Lemma 5

([13] Appendix, Lemma 8)

*Let x*′ ∈ argmax{*N*(*x*) – *π*′*D*(*x*)} and 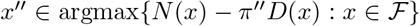 with *π*′ < *π*″, then *D*(*x*′) ≥ *D*(*x*”).

*Proof* Adding the inequalities *N*(*x*″) – *π*′*D*(*π*′) ≥ *N*(*π*″) – *π*′*D*(*π*′) and *N*(*π*″) – *π*″*D*(*π*″) ≥ *N*(*π*′) – *π*″*D*(*π*′) leads to (*π*″ – *π*′)*D*(*π*′) ≥ (*π*″ – *π*′)*D*(*π*″), i.e. *D*(*π*′) ≥ *D*(*π*″) since *π*″ ≥ *π*′ by assumption.

#### Lemma 6

([13] Appendix, Lemma 9)

*Let π*′ ∈ argmax{*N*(*x*) – *π*′*D*(*x*)} *and* 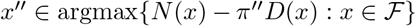, *then* 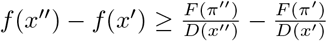

*Proof* From *F*(*π*″) = *N*(*π*″) – *π*″*D*(*π*″) ≥ *N*(*π*′) – *π*″*D*(*π*″) it follows 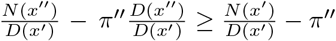. This implies:

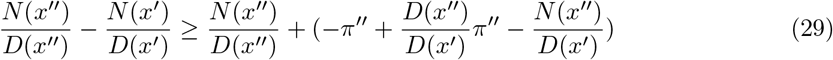

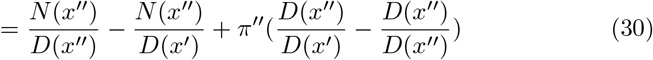

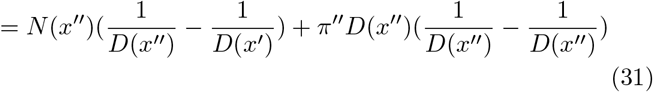

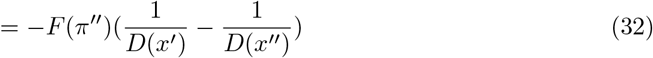

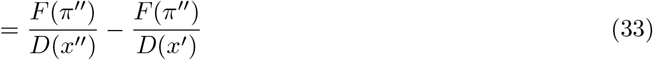

#### Lemma 7

([13] Appendix, Lemma 10)

*Let π*′ ∈ argmax{*N*(*x*) – *π*′*D*(*x*)} *and* 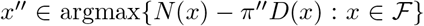 *and F*(*π*\*) = 0, *then if follows for π*′ ≤ *π*″ ≤ *π*\*, *that* 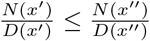.

*Proof*

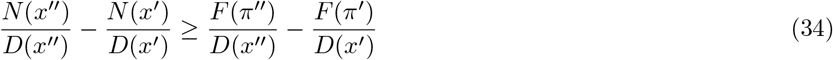

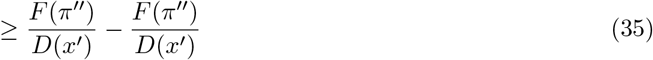

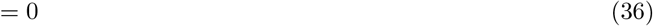

The first inequality follows from lemma 9, the second from lemma 7 and 8.

#### Lemma 8

([13] Appendix, Lemma 11)

*Let π*′ ∈ argmax{*N*(*x*) – *π*′*D*(*x*)} *and* 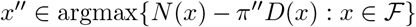, *then* 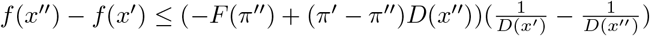.

*Proof* From *N*(*π*′) – *π*′*D*(*π*′) ≥ *N*(*π*″) – *π*″*D*(*π*″) it follows 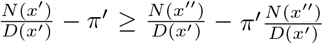 by dividing by *D*(*π*′) > 0. It then follows:

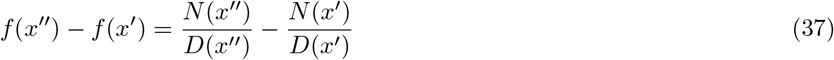

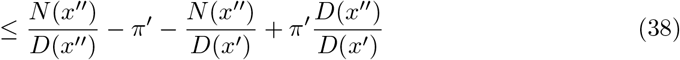

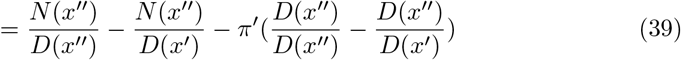

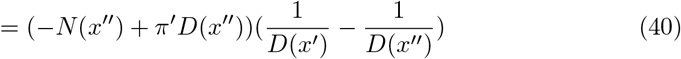

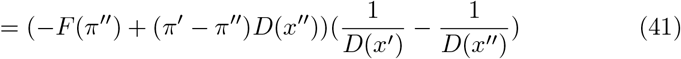

#### Lemma 9

([13] Appendix, Lemma 12)

*Let π*′ ∈ argmax{*N*(*x*) – *π*′*D*(*x*)} *and* 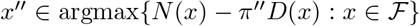 with *F*(*π*\*) = *N*(*x*\*) – *π*\**D*(*x*\*) = 0, *then* 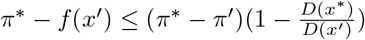.

*Proof*

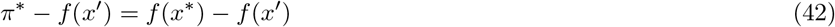

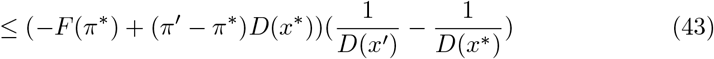

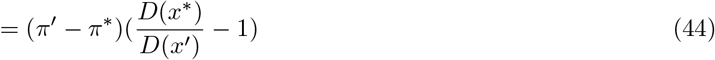

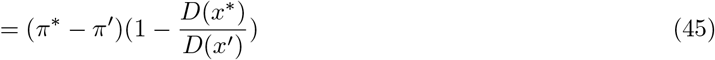

where the inequality follows from Lemma 11.

Proposition 2 can now be demonstrated as follows:

#### Proof of proposition 2

Let *F*(*π*\*) = 0, i.e. 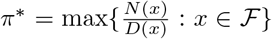. For 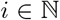, let 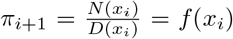 where 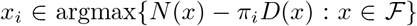 it follows with Lemma 9:

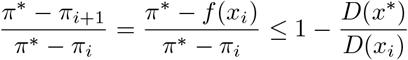

Since 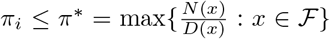 it follows with Lemma 5 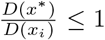 and since 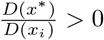 one obtains

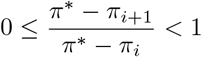

for all 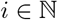. The latter inequality demonstrates superlinear convergence.

##### Correctness of Dinkelbach’s algorithm for solving the DeRegNet model

Here, we prove that the fractional integer programming model for finding deregulated subgraphs proposed in the main text can be solved via Dinkelbach’s algorithm. The only points to clarify are the suitability of Dinkelbach’s algorithm for models with lazy constraints, the suitability of an initial value for π of 0 and the positivity of the objective denominator, see last subsection.

#### Proposition 11

(Dinkelbach-type algorithm for DeRegNet)

*The Dinkelbach algorithm is correct for the fractional integer programming problem of DeRegNet.*

*Proof* The first point to observe is that the objective of DeRegNet is always ≥ 0 hence the initialization condition of the iteration parameter *π* = 0 statisfies *π* ≤ *π*\*. Furthermore, for subgraphs which are constrained to contain at least one node, the denominator of the objective is strictly positive. These two properties are enough to guarantuee convergence of Dinkelbach’s algorithm as detailed above. Also since the original decision variables are also part of the parameterized Dinkelbach iteration problems introducing lazy constraints is technically feasible. Since lazy constraints can only decrease the maximum objective, after every iteration *π* ≤ *π*\* where *π*\* is the optimal objective determined by the current constraints and hence lazy constraints do not interfere with the correctness of Dinkelbach’s algorithm since it requires a starting value of *π* which is a lower bound of the optimal objective value.

Note that lazy constraints effectively amount to restarting Dinkelbach’s algorithm (in a valid initialization state) every time a lazy constraint is added. Hence, convergence can also only be considered superlinear (see last subsection) with respect to the current optimal objective determined by the lazy constraints.

#### Reformulation-Linearization methods

##### Generalized Charnes-Cooper method

The so called Generalized Charnes-Cooper transformation [13] described in this subsection derives its name and general idea from the classical Charnes-Cooper transformation [14] used to solve continuous fractional linear problems. Consider the above general form of a FMIP in the following slightly more detailed format:

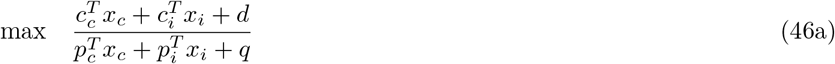

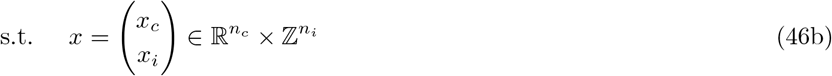

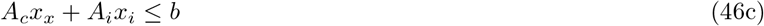

where we explicitly decomposed the variable *x* into its continuous and integer parts. Analogously we have 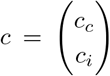 with *c_c_* ∈ ℝ^*n_c_*^, *c_i_* ∈ ℝ^*n_i_*^, and 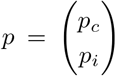 with *p_c_* ∈ ℝ^*n_c_*^, *p_i_* ∈ ℝ^*n_i_*^, and *A* = (*A_c_ A_i_*) with *A_c_* ∈ ℝ^*m*×*n_c_*^, *A_i_* ∈ ℝ^*m*×*n_i_*^. As detailed in [13] one can now define additional variables 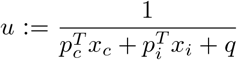 and 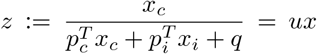. Note, since we assume that there exists some real *m* > 0 such that *p^T^x* + *q* > *m* for all feasible *x* ∈ ℝ^*n*^, it follows that *u* > 0. After incorporating the definition of *u* as a further constraint and multiplying all constraints with *u* one arrives at the following quadratic mixed-integer problem:

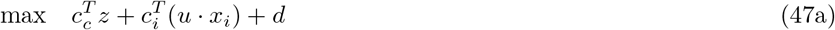

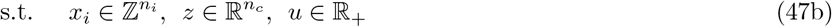

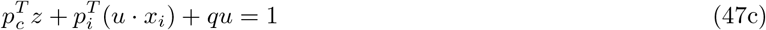

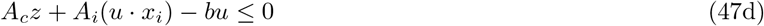

Note that the above problem is not a MILP but a quadratic mixed-integer problem due to the terms *ux_i_* in the transformed constraints. This is addressed in the next subsection. With the notation of this subsection one can formulate the following propositions formalizing the equivalence of the two model formulations [13]:

#### Proposition 12

(Feasible points of the generalized Charnes-Cooper transform) *A point* (*x_c_,x_i_*) *is a feasible solution of problem (A.30) if and only if* (*z,x_i_,u*) *is a feasible solution of problem (A.31*).

*Proof* Because of 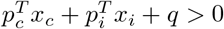 this is true by definition of *u* and *z*.

#### Proposition 13

(Equivalence of solutions of the generalized Charnes-Cooper transform)

*An feasible point* 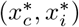 *of (A.30) is optimal if and only if* (*z*\*, 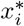, *u*\*) *is optimal for (A.31). It holds that* 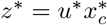 and 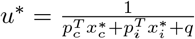.

*Proof* By definition of *u* and *z* the objectives of (A.30) and (A.31) have the same value for all feasible points. The relations for the optimal points are also true by definition.

With respect to lazy constraints involving the integer variables *x_i_* there do not arise any complications since they are part of both problem formulations. Lazy constraints for the continuous variables *x_c_* require more care due to the necessity to transform the constraints correspondingly. The DeRegNet model does only contain integer (in fact, binary) variables and hence it is straight-forward to incorporate lazy constraints in the solution process in terms of the original model formulation.

##### Linearization of binary-continuous quadratic constraints

In contrast to the iterative Dinkelbach scheme, the reformulation-linearization method described in the last section relies on the linearization of products of integer and continuous variables. Since we only deal with binary variables in this paper, we assume from now on that all integer variables are in fact binary. In case of a proper integer variable 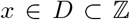, one can introduce auxiliary binary variables 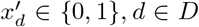 with 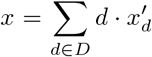 and 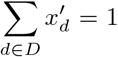 in order to transform its product with continuous variables into a sum of products between binary and continuous variables. There exist variations on the theme of linearization [18], [19], but here we will present the implemented most basic version going back to [20].

Given a continuous variable *v* ∈ ℝ and a binary variable *x* ∈ { 0, 1} one introduces a third (continuous) variable *z* ∈ ℝ corresponding to *z* = *vx* and substitutes any appearance of the product *vx* with *z*. Along with *z* one introduces the following constraints to ensure equivalence:

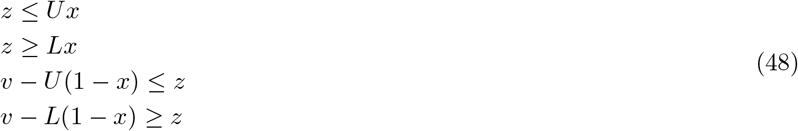

Here, *U* ∈ ℝ is an upper and *L* ∈ ℝ is a lower bound of v which are either given by the problem formulation itself, can be inferred from manual insight into the problem or by solving a certain MILP in some cases. See below.

#### Proposition 14

(Linearization binary-continuous products)

*Let* 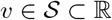 *with bounded* 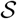 *and let x* ∈ {0,1} *and z* ∈ ℝ. *Furthermore U* ≥ sup 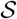 *and L* ≤ inf 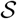. *Then, the constraints (A.15) are statisfied if and only if z* = *vx*.

*Proof* Let *z* = *vx*, then *z* = *vx* ≤ *Ux* since *U* is an upper bound of *v* and *z* = *vx* ≥ *Lx* since *L* is a lower bound of *v*. Also for the case *x* = 1 one has *v* – *U* (1 – *x*)= *v* = *vx* = *z* and *v* – *L*(1 – *x*) = *v* = *vx* = *z* and for the case *x* = 0 the two constraints *v* – *U*(1 – *x*) ≤ *z* and *v* – *L*(1 – *x*) ≥ *z* reduce to *v* ≤ *U* and *v* ≥ *L* respectively which is true by assumption. Conversely, let the constraints in (A.15) be satisfied. Then in the case *x* = 1, the constraints *v* – *U*(1 – *x*) ≤ *z* and *v* – *L*(1 – *x*) ∆ *z* imply *v* ≤ *z* ≤ *v* and hence *z* = *v* = *vx*. In the case *x* = 0 the first two constraints of (A.15) imply *z* = 0 = *vx*.

The lower bound L and the upper bound U can generally be obtained by solving suitable MILPs [13] involving the denominator of the original objective. To obtain the (tightest possible) lower bound one can solve the following problem:

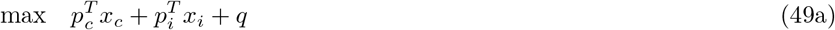

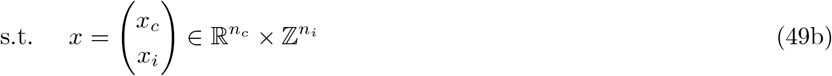

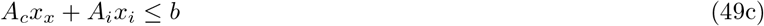

Analogously to obtain the (tightest possible) upper bound one can solve the following minimization problem:

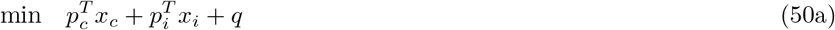

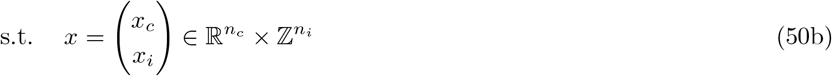

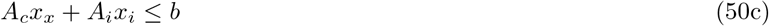

Note however, that any lower and upper bound would work. The trade-off between less tight bounds on the denominator variable and the necessity of solving up to two MILPs up front has to be decided for every model.

In case of DeRegNet, lower and upper bound on the objective denominator are explicitly set in the problem formulation in the form of minimal and maximal subgraph size. Hence one does not have to solve any MILPs up front and has (optimal) lower and upper bounds for the inverse denominator readily available due to the problem formulation.

#### Software for solving fractional integer programs: libgrbfrc

In order to solve the fractional integer programs formulated in the main text, a C++ library based on the commercial Gurobi solver was implemented. libgrbfrc (https://sebwink.github.io/libgrbfrc/) in particular implements the two solution methods from above: Dinkelbach’s algorithm and the generalized Charnes-Cooper transform. Due to the requirements of the developed optimization models (see main text) the implementations support lazy constraints. Academic licenses for Gurobi are readily obtained.

### Lazy constraints in branch-and-cut MILP solvers

For reference this section contains an high-level outline of how lazy constraints fit into branch-and-cut algorithms for solving mixed-integer programs. The exposition is adapted from [21].

Let a MILP with 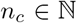 continuous and 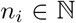 integer variables of the following form be given:

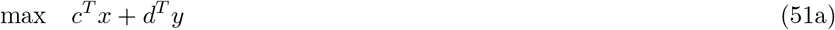

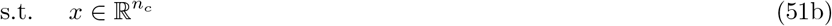

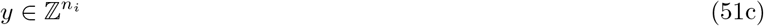

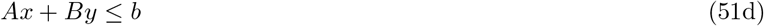

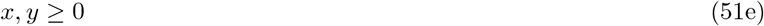

Here *c* ∈ ℝ^*n_c_*^, *d* ∈ ℝ^*n_i_*^, *A* ∈ ℝ^*m*×*n_c_*^ and *B* ∈ ℝ^*m*×*n_i_*^ for some 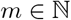. The (*natural) linear programming relaxation* of a MILP of the above form is the following:

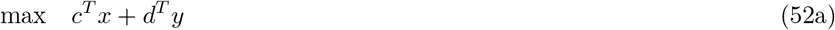

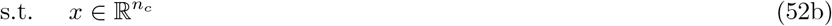

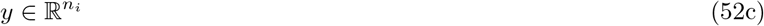

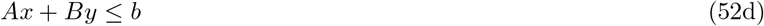

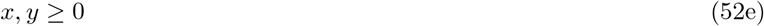

Lazy constraints are constraints which are not initially explicitly part of the model formulation, the reason usually being that it would require an infeasable exponential number of constraints (with respect to the number of variables).

The classical branch-and-cut strategy for solving MILPs with lazy constraints can then be formulated as the following algorithm 2.

**Algorithm 2:**
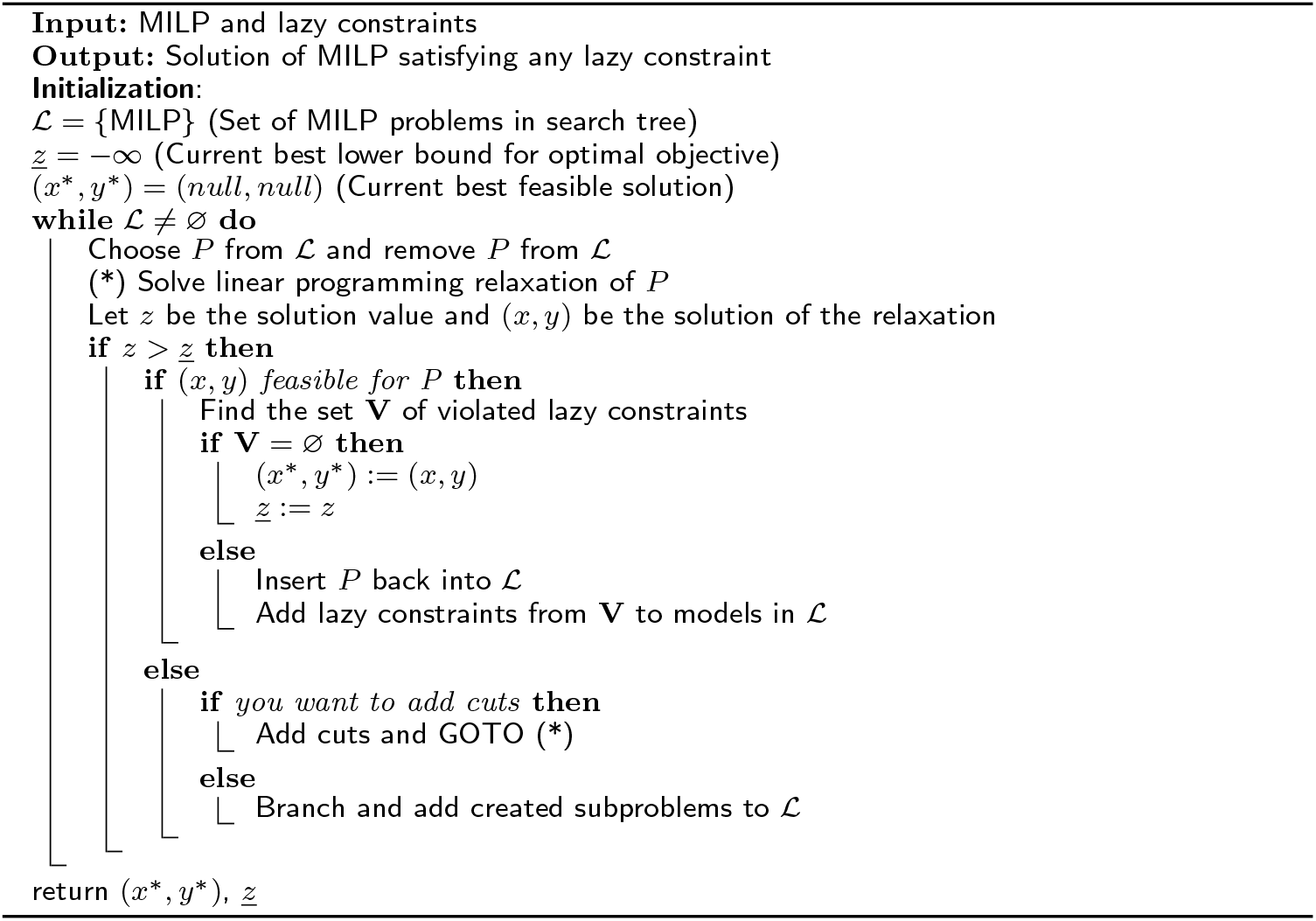
Branch-and-cut for MILPs with lazy constraints

#### Lazy constraints for the DeRegNet model

For DeRegNet the lazy constraint separation subroutine centers around finding the strongly connected components of the given solution. This is generally considered an efficiently solvable problem.

##### Strongly connected components

Given a directed graph *G* = (*V, E*) one says that *G* is strongly connected if and only if there is a directed path from every node *v* ∈ *V* to every other node *u* ∈ *V*. A strongly connected component of a directed graph is any maximal subgraph which is strongly connected, i.e. adding any node not in the subgraph would render the resulting subgraph to be not strongly connected anymore. Sometimes one refers to *V*’ ⊂ *V* as inducing a strongly connected component if the subgraph induced by *V*′ is a strongly connected component. One denotes the set of node sets inducing all strongly connected components of a graph *G* = (*V,E*) by 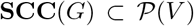. The three classical algorithms which can be used to solve the problem of finding a directed graph’s strongly connected components in *O*(|*V*| + |*E*|) time are the Kosarju-Sharir algorithm [22], Tarjan’s algorithm [23] and variants of the path-based strong component algorithm [24]. A strongly connected *subgraph* (in contrast to *component*) is a subgraph of a graph which is strongly connected.

##### Lazy constraint separation subroutine of DeRegNet

This subsection and algorithm 3 provide the details on the lazy constraint separation subroutine employed for the solution of the DeRegNet model. The formal details are given as algorithm 3. In short, given a (potential) incumbent solution to a DeRegNet instance not containing all strong-component constraints, the subroutine finds the strongly connected components of the corresponding subgraph and checks whether any such component either contains the root node itself or has at least one incoming edge from within the subgraph but from outside the component. If so, the (potential) incumbent is feasible, hence an actual incumbent solution. Otherwise the violated constraint is added to the model in while the (potential) incumbent is declared infeasible. The general implementation strategy employed is based on the one given by [1] where cycles are detected in order to avoid unconnected subgraphs.

**Algorithm 3:**
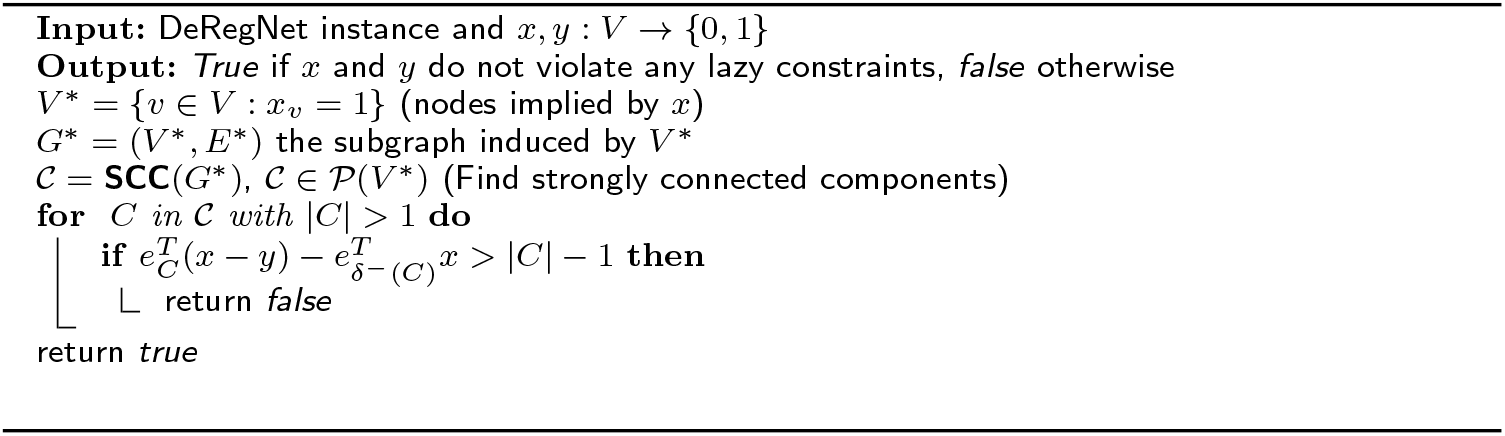
Lazy constraint subroutine for DeRegNet. In case a potential incumbent is found all strongly connected components are checked to assess feasibility. In case any strongly connected component does not contain the root node and has no incoming edges from another component, a (lazy) constraint enforcing the requirement is added. **SCC**(*G*) denotes the set of all strongly connected components of a graph *G*.

#### Further technical aspects of solving DeRegNet models

##### Primal heuristics for the DeRegNet model

Every feasible solution of a mixed-integer program provides a lower bound on the optimal solution value (for maximization problems). The feasible solution which currently gives the best lower bound on the optimal value during a branch-and-bound procedure is called the *incumbent* (*solution*). Branch-and-bound (and hence branch-and-cut) for mixed-integer programs relies on pruning parts of the search tree of LP relaxation subproblems by assessing whether the optimal solution value of a given LP relaxation is less than the best lower bound provided by the incumbent. Primal heuristics [25] aim at finding and/or improving feasible solutions during a branch-and-bound procedure. While some generic methods for primal heuristics exist [26], [27], [28], [29], [30], they tend to be highly problem-specific [25]. Of special interest in that context are primal heuristics for the MWCSP [11], [6], [7]. In the following I describe start and improvement heuristics useful during the solution of DeRegNet instances.

###### Start heuristics

A priori there is no feasible solution known at the beginning of a branch-and-bound procedure for solving a mixed-integer program. Heuristics which try to find initial feasible solutions are called *start heuristics*. I outline two start heuristics which can be employed at the beginning of the branch-and-bound search for the solution of the DeRegNet model.

###### Greedy start heuristic

The first start heuristic is called *greedy start heuristic* and basically starts with the highest scoring node and greedily adds neighbors of already added nodes until the average score of the thus defined subgraph starts decreasing. If the currently selected subgraph is feasible upon termination, one has found a feasible solution. The formal procedure is outlined in algorithm 4. There are a number subtleties attached to this start heuristic. First and foremost the procedure only assures the reachability constraints regarding the root node. Most other constraints may or may not be satisfied at any given time during the procedure, mostly: subgraph size constraints and constraints ensuring the necessity of leaf nodes to be from the subset of terminal nodes. While the subgraph size constraint is relatively easily manageable by stopping the procedure when the maximal subgraph size is reached and by restarting in case the minimal subgraph size can not be achieved in the first place. In the latter case, one can restart the procedure from the best scoring node not already selected during earlier attempts of the greedy start heuristic. The issue of the terminal node constraints is not easily handled and hence the greedy start heuristic is in effect only usable in case *T* = ∅. Also instances with **Inc** ≠ ∅ cannot be handled by this heuristic.

###### Receptor-terminal shortest path heuristic

The second start heuristic is more suitable in situations where there is a non-empty terminal set *T*. In short, it finds the shortest path between a pair of receptor and terminal nodes with high node scores. The **SHORTEST PATH** subroutine referenced in algorithm 5 can be an implementation of any of the canonical algorithms to find single-source shortest paths with unit edge weights in directed graphs in polynomial time [31], [32], [33]. Subject to **Ex** = **Inc** = ∅ all connectivity constraints will be satisfied by construction. If the subgraph size constraints are met is up to chance however. Again, running multiple times with the, say *K*, highest scoring pairs of receptors and terminals, can help in this situation. Note, that the restriction of **Ex** = **Inc** = ∅ could be lifted by formulating the corresponding shortest path problem by canonical means in terms of integer programming problems [34]. This possibility is not explored further however since solving integer programs to get initial feasible solutions to integer program may be a slippery slope. In particular in the case of DeRegNet, where the main problem to solve is formulated in terms of decision variables corresponding to nodes while shortest path integer programming formulations usually introduce decision variables corresponding to the edges of the graph.

**Algorithm 4:**
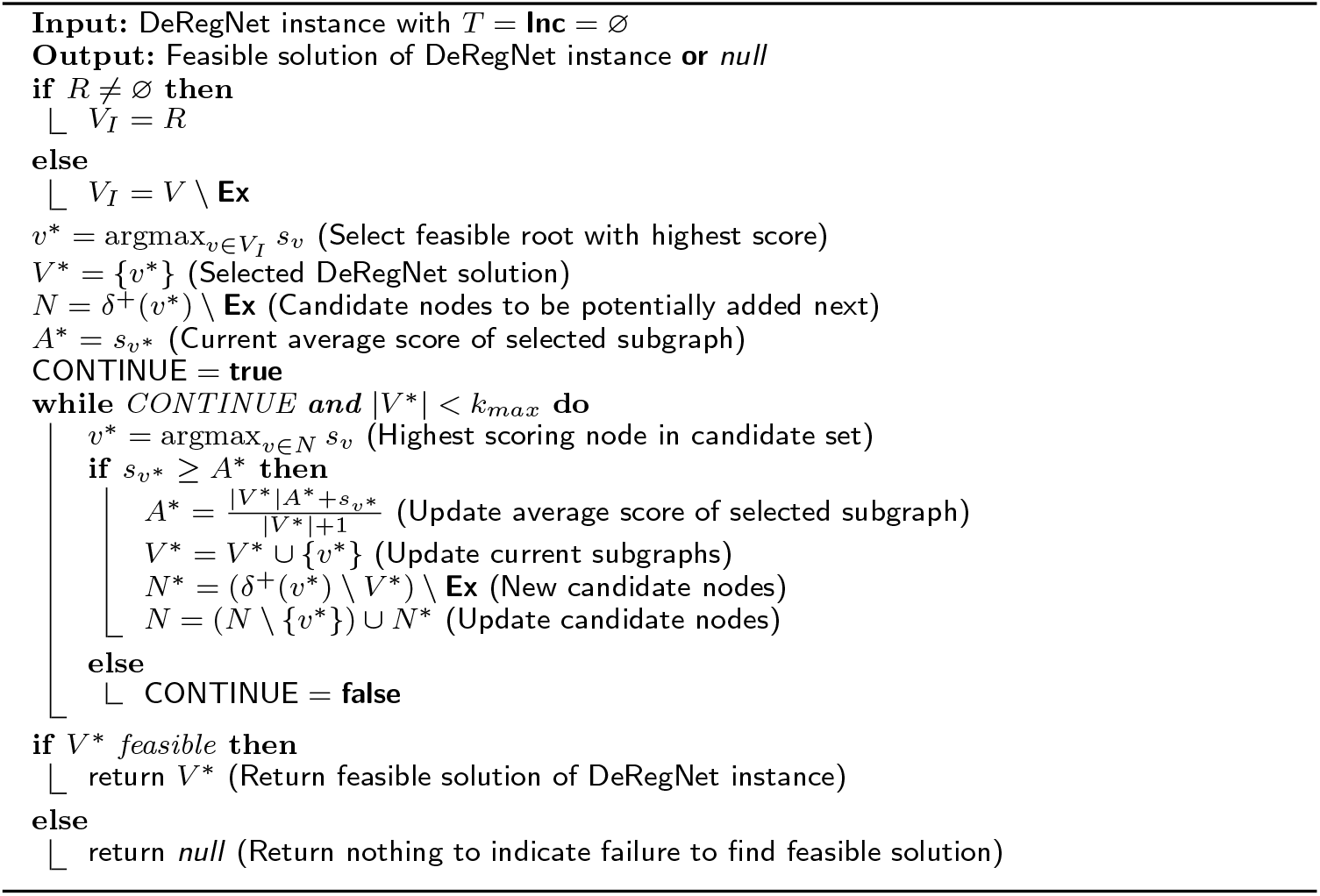
Greedy start heuristic for the DeRegNet model

##### Improvement heuristics

In case a feasible solution is found at a particular branch-and-bound node (which may be a new incumbent or not), heuristics which try to improve that given feasible solution are called *improvement heuristics*. Here I describe a simple greedy improvement heuristic which can be applied to any feasible solution, either found during the branch-and-cut procedure or otherwise. It works analogously to the greedy start heuristic (algorithm 4), the only difference being that one is already starting with a feasible solution. In particular, the heuristic can be applied to solutions constructed by the receptor-terminal shortest path start heuristic (algorithm 5) described in the previous section. Trying to improve the greedy start heuristic (algorithm 4) with the improvement strategy outlined below is futile however since by construction the former already added all potential subgraph nodes in a greedy fashion. During a branch-and-cut run any new feasible solution can potentially be improved by the heuristic. In case of an incumbent one can hope for an even better incumbent, in case of a feasible solution one can hope to improve it up to a point where it actually becomes a new incumbent. The description of the heuristic is provided as algorithm 6.

**Algorithm 5:**
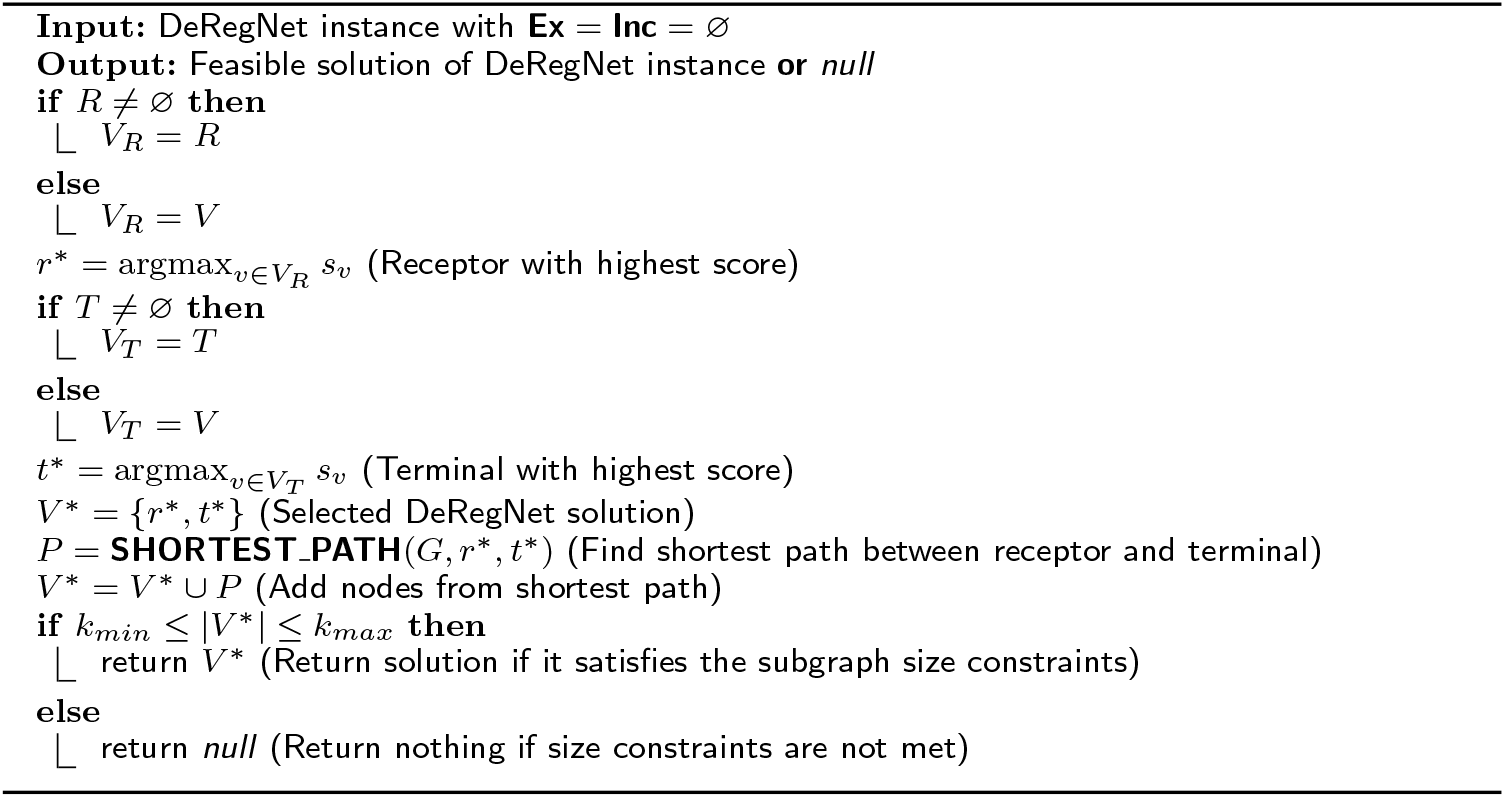
Receptor-terminal shortest path start heuristic for the DeRegNet model

**Algorithm 6:**
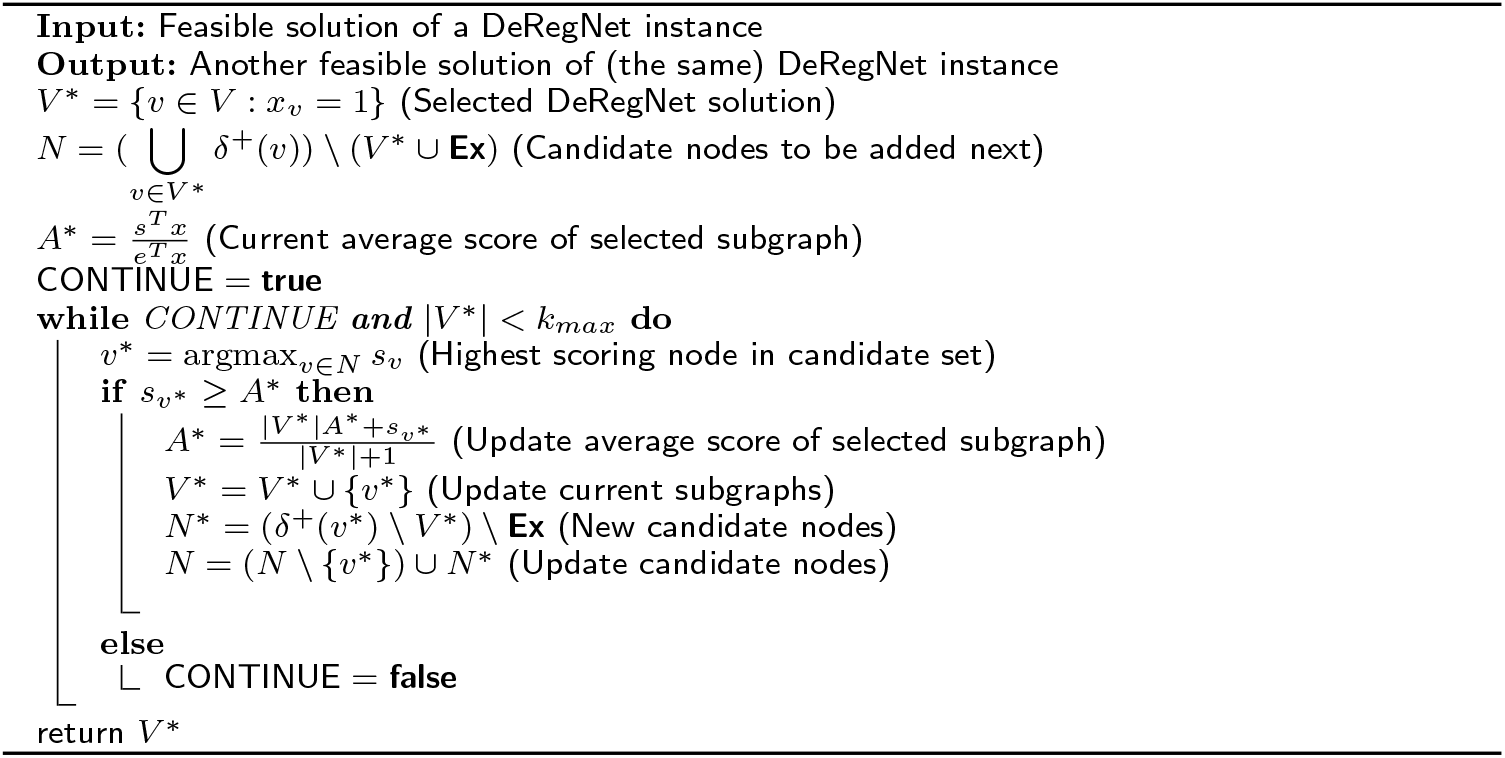
Greedy improvement heuristic for the DeRegNet model

#### Approximate solutions via branch-and-bound gap cut

One can use a mixed-integer programming solver generically to obtain suboptimal solutions to a given (maximization) MILP with optimal objective value *z*\*. During the branch-and-cut search one obtains lower bounds on the optimal value by feasible solutions to the problem and an upper bound by the solution value of the initial LP relaxation of the problem. Let 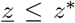 be the best available lower bound and let 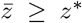 be the upper bound obtained by the relaxed problem. The *relative gap* λ_*rel*_ during a branch-and-cut search is defined as 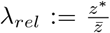. With the upper bound 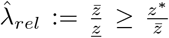 on the gap it follows that 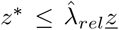 and hence 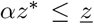 with 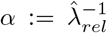. Stopping the branch-and-cut procedure at the given gap upper bound value hence provides an approximate solution of a posteriori approximation guarantee of 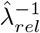. I refer to the strategy of stopping the branch-and-cut search once the gap upper bound is below a certain threshold as *gap cut* or *gap (cut) thresholding.* Employing the gap cut strategy can be useful in situations where the MILP solver can find reasonably good solutions in reasonable time but would take significantly more time to find the optimal solution. The option of to carry out gap cut thresholding is incorporated in the implementation of DeRegNet for this very reason.

#### Caching transformed model formulations

For DeRegNet’s use cases it is quite common to optimize DeRegNet instances which just differ in terms of their node scores, i.e. share the same underlying graph. For example, finding deregulated subgraphs for individual cases in a TCGA cohort with a fixed regulatory network derived from KEGG will require to solve a model with the same structural properties but with differing score data, for example a omics-readout for every case in the cohort. In particular, in such a situation the reformulation and linearization procedure of the generalized Charnes-Cooper transform only has to be carried out once and can be reused across cases since it does not depend structurally on the objective data vector *s*. While solution time of a DeRegNet instance with the generalized Charnes-Cooper transform tends to be dominated by the time to solve the resulting integer linear program, reuse of the transformed model structure can nonetheless result in significant computational savings.

### Further details on benchmarking DeRegNet

Algorithm 7 details the benchmark instance simulation algorithm. Algorithm 8 details the mode of application of [1] in the context of the benchmarks described in the main part of the paper. Figure 1 depicts the subgraphs simulation procedure conceptually.

### DeRegNet subgraph derived features for predicting survival

Predicting phenotypes based on clinical and molecular data is one of the big challenges on the road to personalized medicine. A frequently readily available phenotype for cancer patients is survival time (i.e. the time from disease onset/diagnosis to (possibly disease induced) death). Improving upon clinical predictors with molecular data often still poses significant challenges [35]. Here, we provide an example of the suitability of deregulated subgraph-derived features for predicting survival in the TCGA-LIHC dataset. In particular, we demonstrate that predictions based on subgraphs is at least as good GSEA-based predictions obtained in a comparable manner. Furthermore, subgraph derived features can improve upon predictions based on clinical features alone.

**Algorithm 7:**
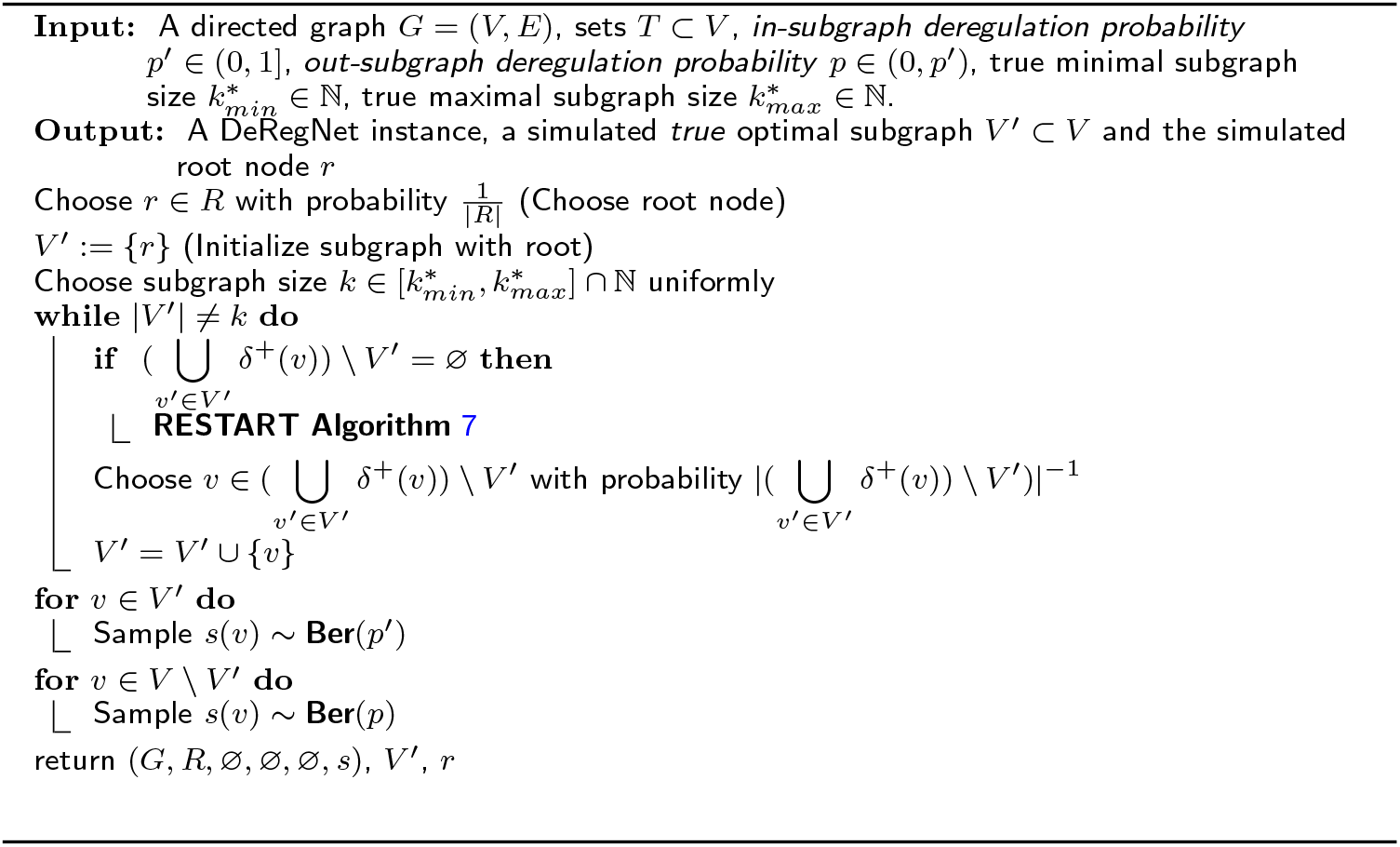
Simulating DeRegNet instances with known ”optimal” subgraph. Here, **Ber**(*p*) denotes a Bernoulli random variable with parameter *p* ∈ [0,1]. Note that the true minimal subgraph sizes 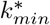 and 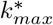 can be different than the minimal and maximal subgraph sizes *k_min_* and *k_max_* specified when solving related DeRegNet or Backes et al. instances.

**Algorithm 8:**
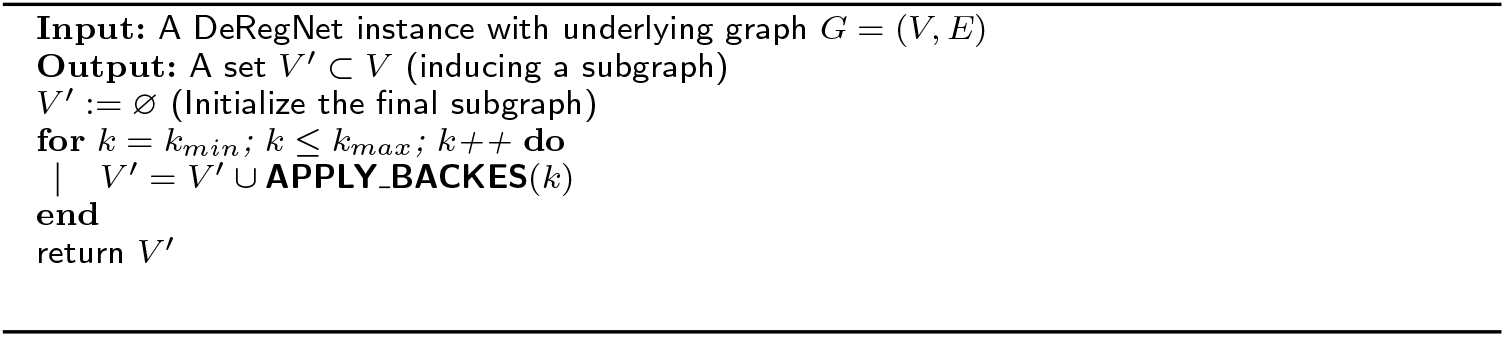
Applying [1] for benchmarking DeRegNet. Here, **APPLY_BACKES**(*k*) refers to applying the algorithm of [1] with fixed subgraph size *k*, understood to return a set of nodes corresponding to the induced subgraph found by the run.

**Figure A 1:**
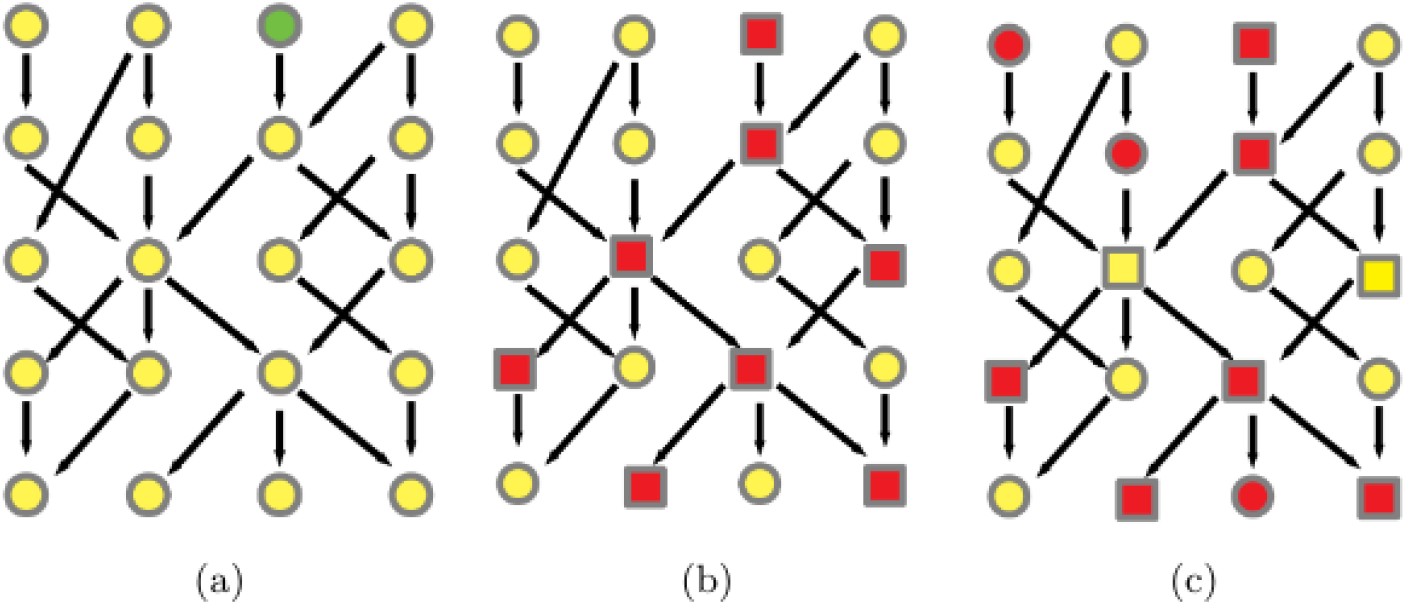
Simulating DeRegNet instances. See algorithm 7 for a formal outline of the simulation procedure. On a high level it proceeds like this: **(a)** Choose root node randomly. **(b)** Simulate a feasible subgraph by randomly choosing nodes maintaining the topological constraints of the model. Set the deregulation score of nodes in the subgraph to one. **(c)** Introduce noise by flipping node deregulation scores randomly.

#### Data preparation and feature engineering

Survival times were binarized by labeling all patients with survival less than three years (1095 days) as bad outlook patients (*y* = 0) and all patients with last followup time larger than three years as good outlook patients (*y* = 1). The resulting dataset consisted of 198 patients from the TCGA-LIHC cohort. For every case the following features are derived:

- **clinical**: Features from clinical data comprising age, *gender, body mass index (BM1), tumor stage* (!) and *tumor morphology.* Age (in years) and BMI were scaled via z-scores. Tumor stage and morphology where one-hot encoded.
- **gsea**: Features derived from (single sample) Gene Set Enrichment Analysis (GSEA) [36]. Two lists of significantly enriched pathways w.r.t good outcomes vs. bad outcomes and vice versa were computed by (standard) GSEA. From every list I retained pathways with adjusted p-value less than 0.1, which resulted in a total of 14 KEGG pathways. After performing ssGSEA, every sample received the corresponding personalized ssGSEA enrichment scores for these pathways as a 14-dimensional feature vector. The above steps were carried out with *gseapy* (http://gseapy.rtfd.io/). For more information on single-sample GSEA, see [37]. The obtained features were scaled via z-scores.
- **subgraph-overlap**: Features based on up- and downregulated subgraphs for the good and bad outcome subgroups. Subgraphs were computed based on the global deregulation score for the good outcome and bad outcome patients respectively (on the respective training sets only, see below). Every sample is then associated with the regulation-aware node overlap between its personalized de-, up- and downregulated subgraphs and up- and downregulated global subgraphs for the good and bad outcome subgroups respectively. The deregulation-aware node overlap is defined as follows. Given two (induced) subgraphs *V*′,*V*″ ⊂ *V* and node scores *s*′, *s*″: *V* → { −1, 0,1} the deregulation-aware node overlap is defined as 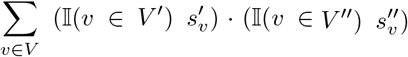. This amounts to a 12-dimensional feature vector. Again, z-scores were applied.
- **ndcg**: Subgraph features derived from network-defined cancer genes. After identifying network-defined cancer genes (see previous subsection) for de-, up- and downregulated subgraphs one obtains a binary indicator for every case representing whether it contains any given such gene or not, leading to 15-dimensional feature vectors corresponding to 15 network-defined cancer genes.
- **subgraph**: *subgraph-overlap* and *ndcg* combined (concatenated).

Under a *feature combination* it is understood the combination of two or more of the just defined features. In the following, I use a plus sign to indicate feature combinations, e.g. *subgraph* = *subgraph_overlap* + *ndcg.* As another example, *subgraph* + *clinical* then denotes *subgraph* features combined with *clinical* features.

#### Survival prediction with clinical, pathway and subgraph features

The experiments described in the following were carried out with scikit-learn (https://scikit-learn.org). Every feature/feature combination was tested by training a Support Vector Machine, a simple artificial neural network, a random forest and a logistic regression. For every algorithm we performed an algorithm-specific grid search for model selection. The grid search was equivalent for different feature combinations in order to be able to assess the comparative suitability of the features. Final models were evaluated with 6-fold cross validation estimating mean Receiver Operating Characteristic (ROC) curves and Area under the curve (AUC) scores.

Features *gsea* and *subgraph-overlap* are roughly equivalent with respect to the underlying logic, with subgraphs or pathways as contextual data inputs respectively. Hence, comparing these two features may give an indication of the suitability of subgraph vs. pathway methods for feature engineering for survival prediction. Figure 2 shows that the *subgraph_overlap* features hold promise w.r.t *gsea* features.

Furthermore, it has been shown that improving upon clinical features with molecular features for survival prediction is not an easy task [35]. The experiments conducted here show that for the given setting, prediction models combining clinical and subgraph features (based on molecular interactions and data) provide performance gains compared to a purely clinical model. Also, the subgraph features achieve parity with classifiers based on clinical data alone. Figure 3 represents these findings.

**Figure A 2:**
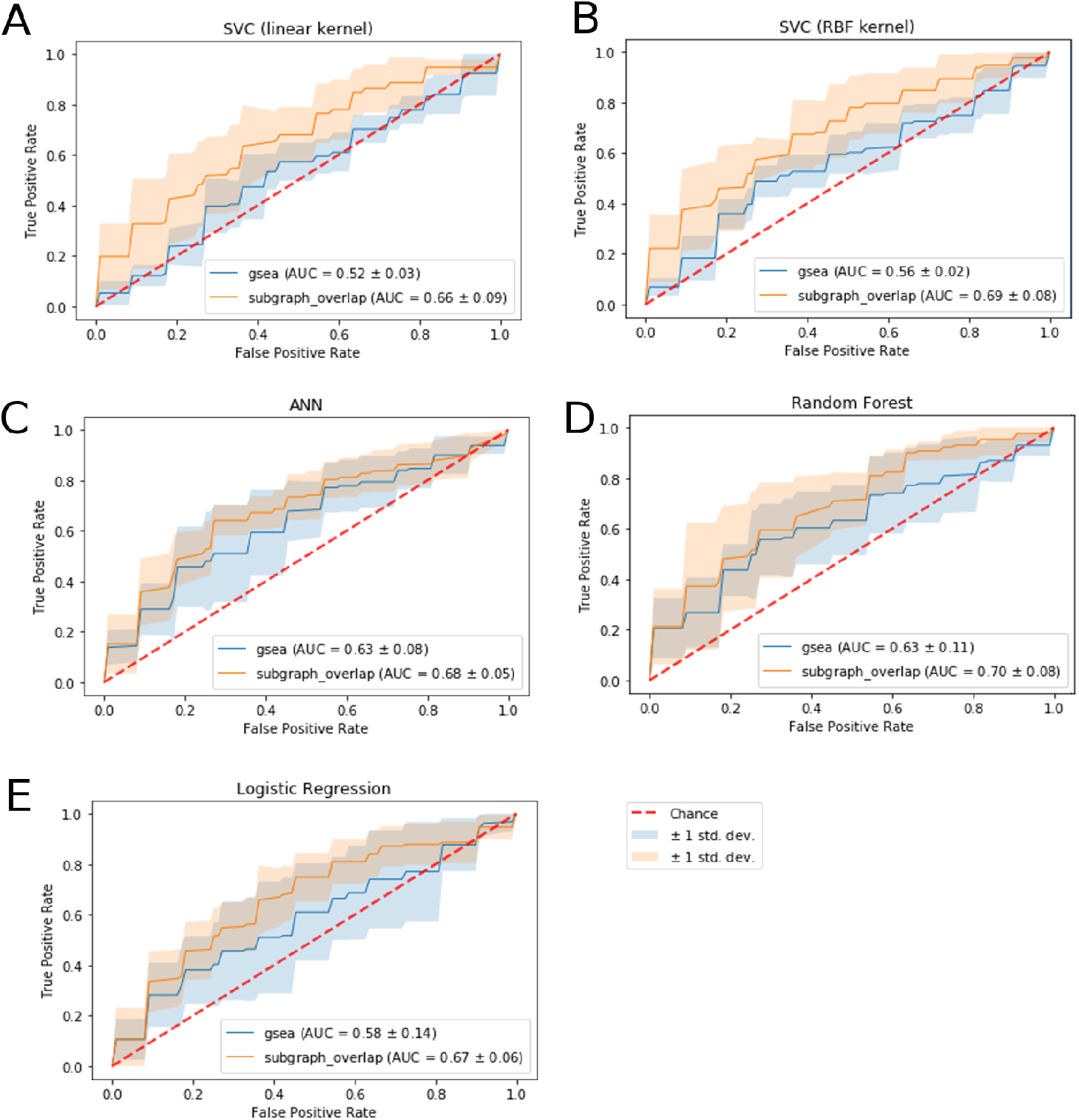
Subgraph features vs. GSEA features across models. (A) Support Vector Classifier (SVC) with linear kernel (B) Support Vector Classifier (SVC) with radial basis function (RBF) kernel (C) Artificial Neural Network (ANN) (D) Random forest (E) Logistic Regression.

---

**Figure A 3:**
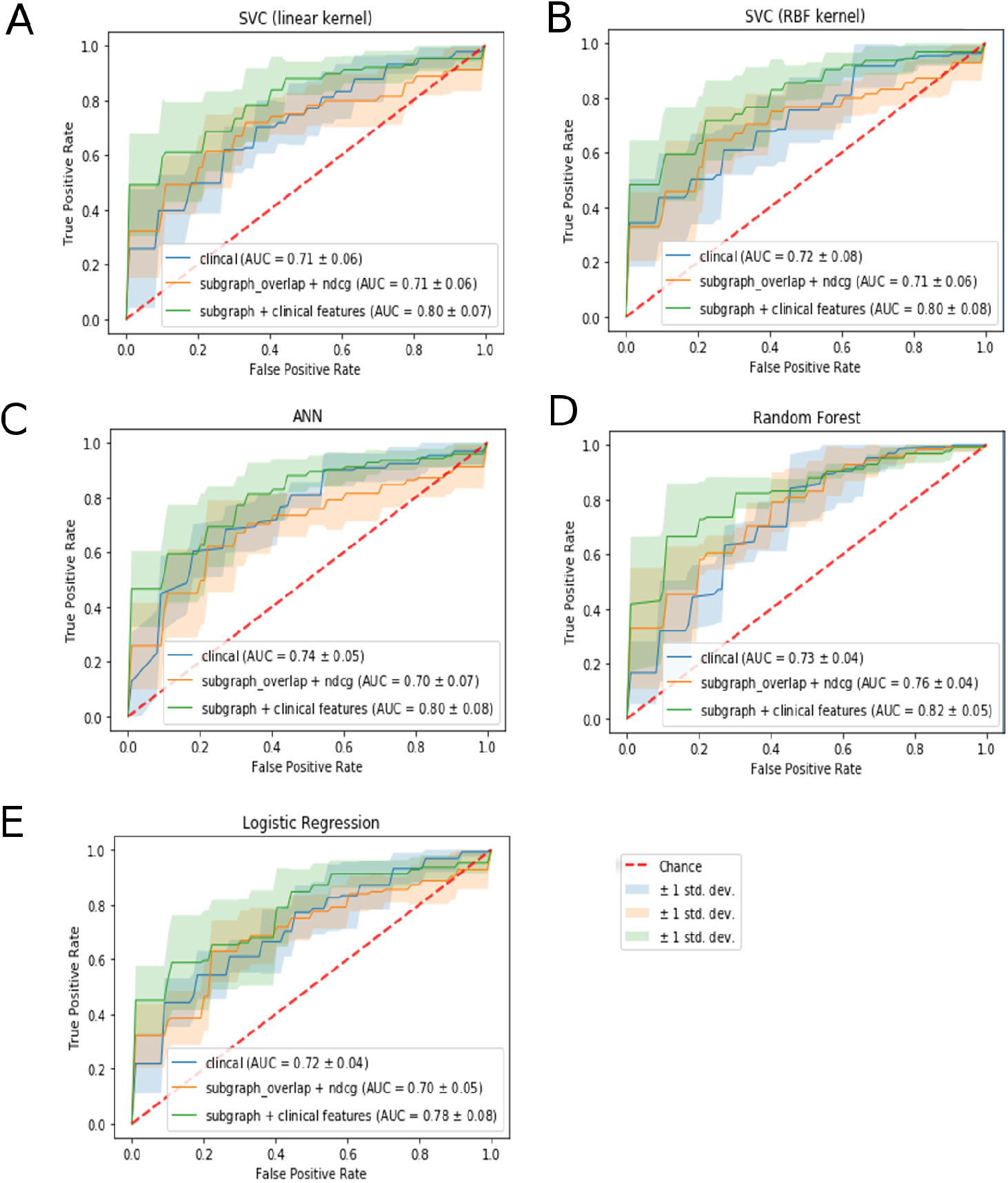
Subgraph features vs. clinical features across models. Clinical + subgraph features together outperform either in isolation. Subgraph features perform comparably to clinical features. (A) Support Vector Classifier (SVC) with linear kernel (B) Support Vector Classifier (SVC) with radial basis function (RBF) kernel (C) Artificial Neural Network (ANN) (D) Random forest (E) Logistic Regression.

## Abbreviations

**Acronyms**

ANN: Artificial Neural Network. 30, 31
AUC: Area under the curve. 29
BMI: Body Mass Index. 28
GSEA: Gene set enrichment analysis. 28, 30
KEGG: Kyoto Encyclopedia of Genes and Genomes. 26, 28
LIHC: Liver Hepatocellular Carcinoma. 28
MAWCSP: Maximum Average Weight Connected Subgraph Problem. 9
MWCSP: Maximum Weight Connected Subgraph Problem. 8
RBF: Radial Basis Function. 30, 31
RMAWCSP: Rooted Maximum Average Weight Connected Subgraph Problem. 9
RMWCSP: Rooted Maximum Weight Connected Subgraph Problem. 8
ROC: Receiver Operating Characteristic. 29
ssGSEA: Single Sample Gene Set Enrichment Analysis. 28
SVC: Support Vector Classifier. 30, 31
TCGA: The Cancer Genome Atlas. 26, 28
w.r.t: with respect to. 28, 29

## Supplementary Figures

**Figure S 1:**
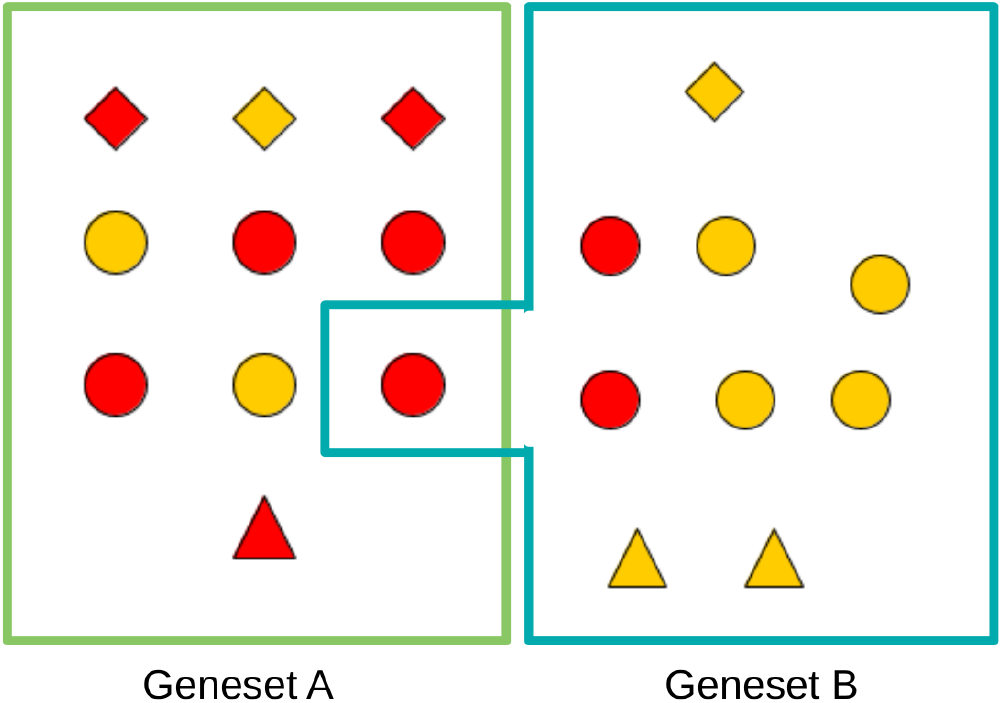
Conceptual view of classical pathway/gene set analysis. Gene sets/pathways are considered merely as sets of genes ignoring any explicit biomolecular interactions between the elements of a gene set/pathway. Here red nodes represent differentially regulated genes and a basic GSE analysis employing hypergeometric Over-representation analysis (ORA) would test for more red nodes than expected in any given gene set.

**Figure S 2:**
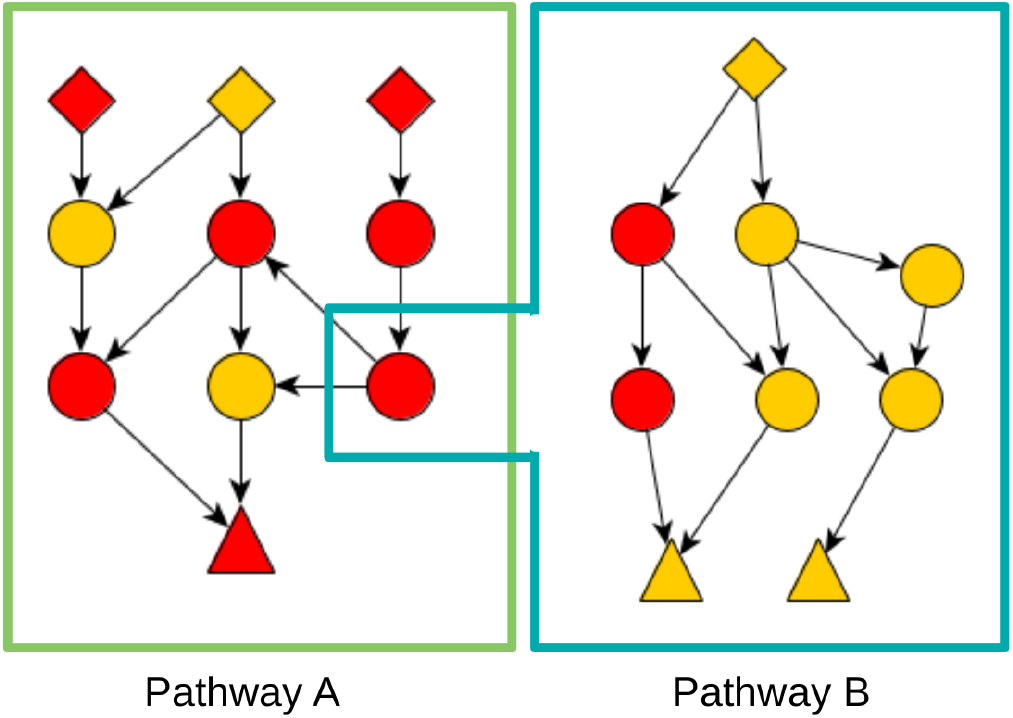
Conceptual view of topological pathway/analysis. Biomolecular interaction are taken into account when calculating enrichment for any given pathway. Gene sets/pathways are still predefined though and interactions between pathways are usually not taken into account

**Figure S 3:**
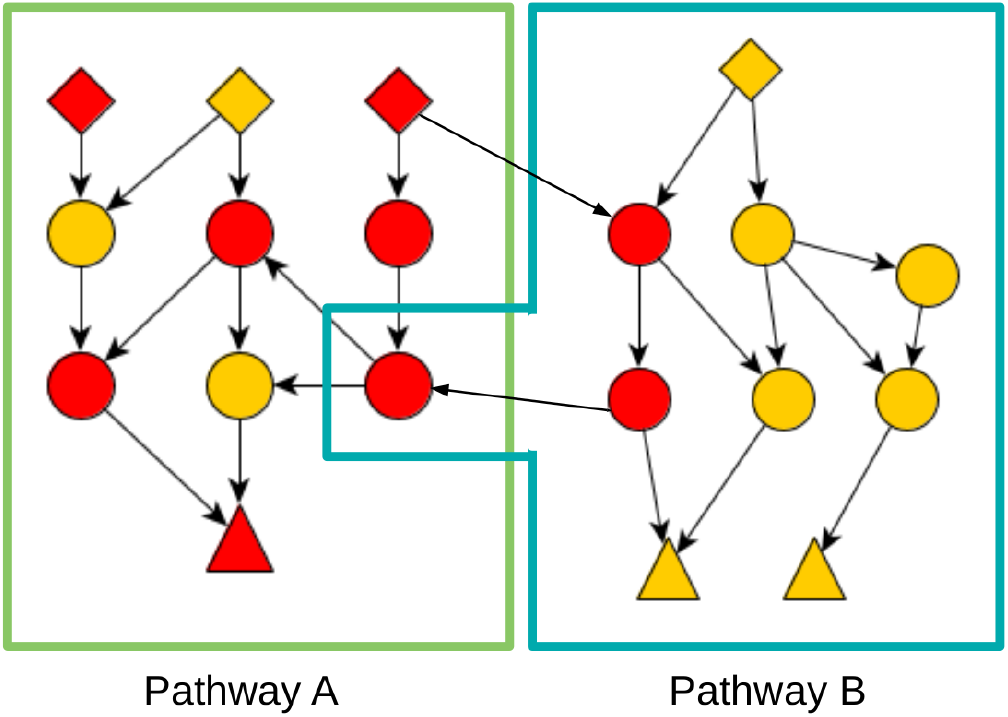
Conceptual view of topological pathway/analysis with pathway crosstalks. Pathway crosstalks happen when genes are part of multiple pathways. They can also happen if there are genes in two pathways with interactions between them from another pathway. Even with pathway crosstalks accounted for, the gene sets/pathways as such are still predetermined.

**Figure S 4:**
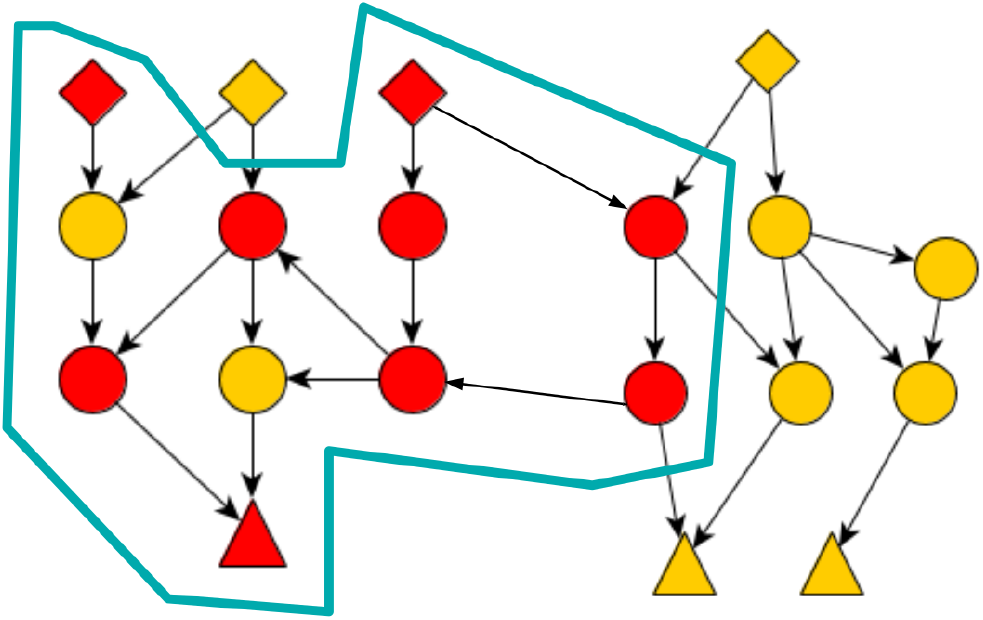
Conceptual view of de-novo pathway analysis. De-novo pathway identification / deregulated subnetwork discovery drops the predetermined pathways and defines enriched subnetworks/pathways from the omics data itself.

**Figure S 5:**
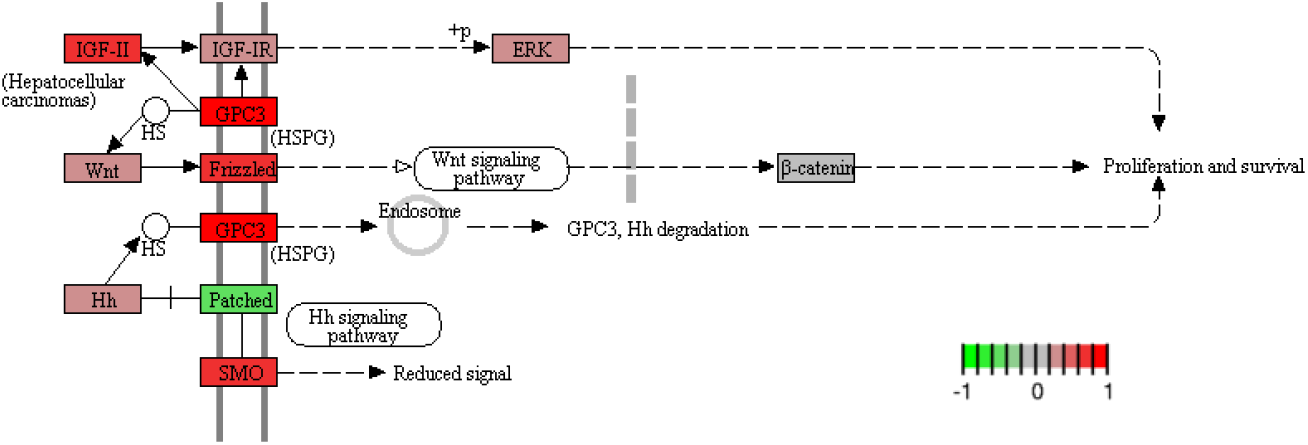
GPC3-mediated activation of WNT signaling is a well-documented process in liver cancer. The figure shows the relevent KEGG map (Proteoglycans in cancer: hsa05205) with TCGA-LIHC min-max-scaled log_2_ fold changes mapped onto the genes. This process was automatically recaptured by our upregulated subgraphs for TCGA-LIHC.

**Figure S 6:**
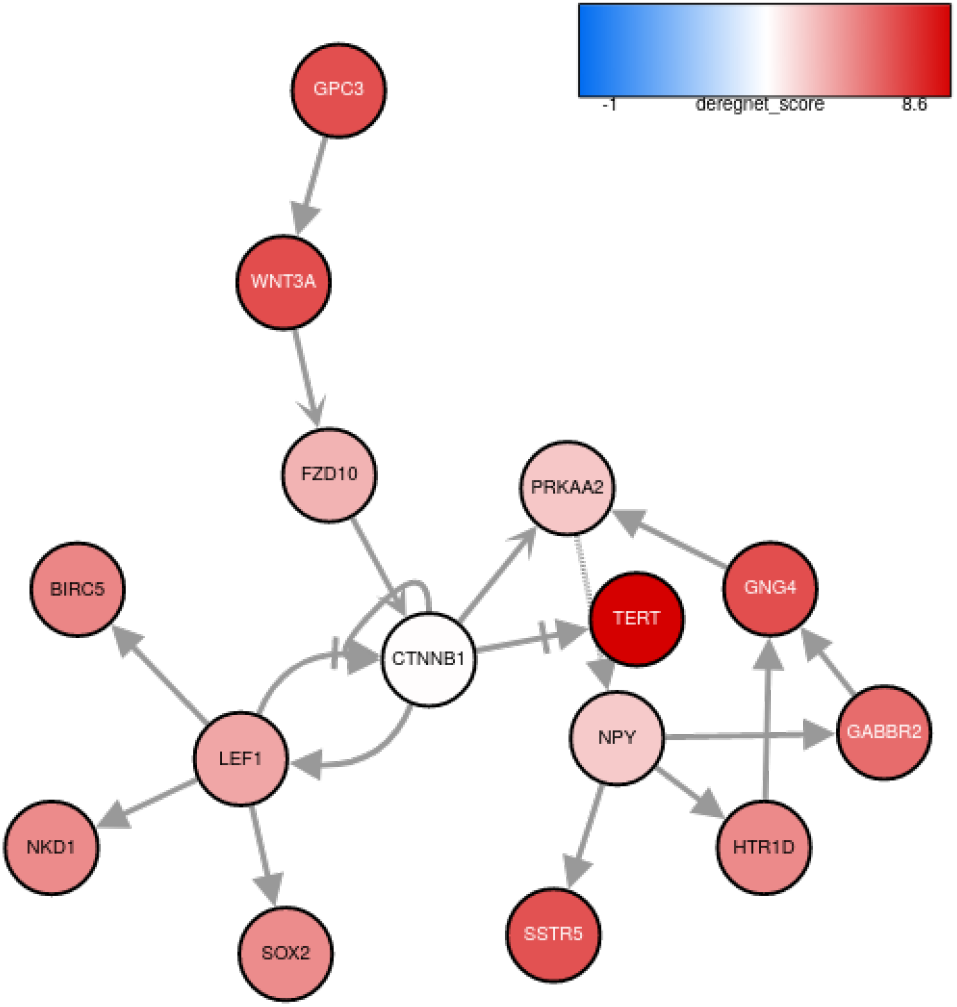
Optimal upregulated global subgraph for TCGA-LIHC.

**Figure S 7:**
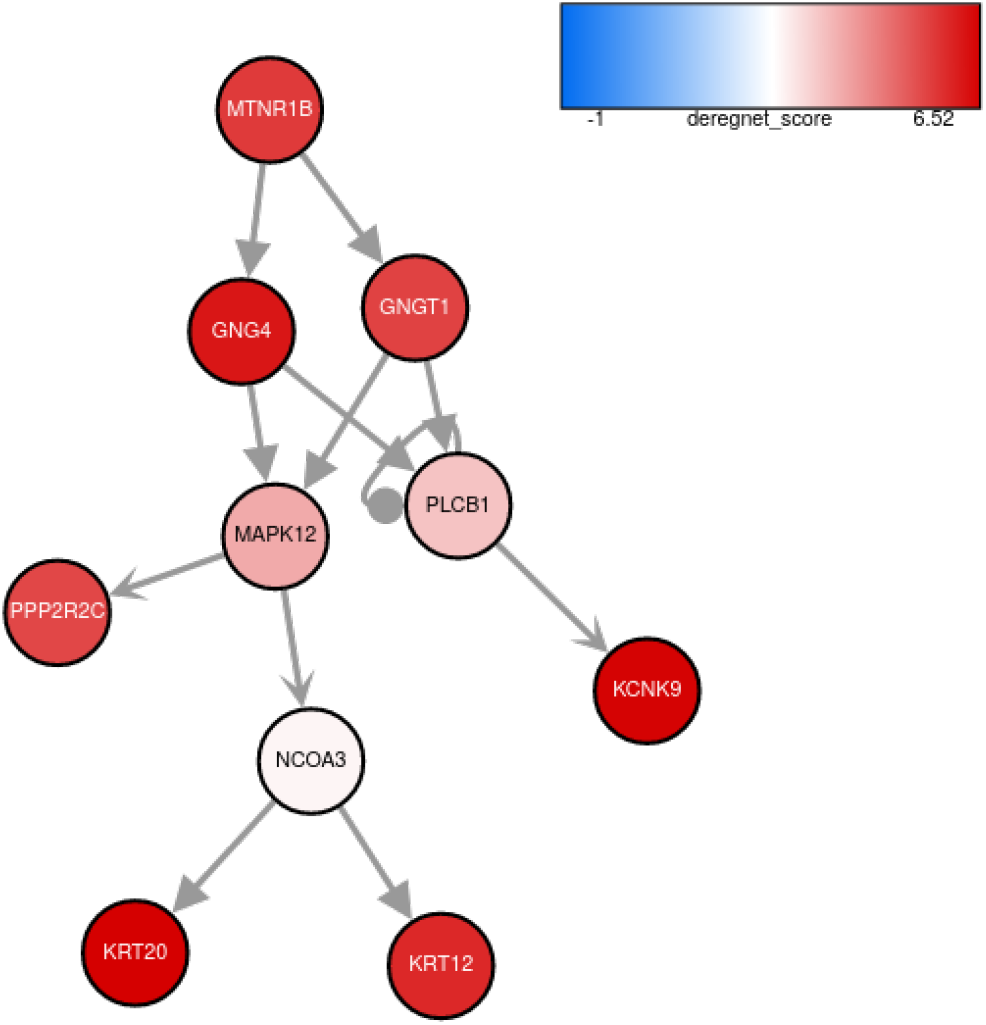
1^*st*^ suboptimal upregulated global subgraph for TCGA-LIHC.

**Figure S 8:**
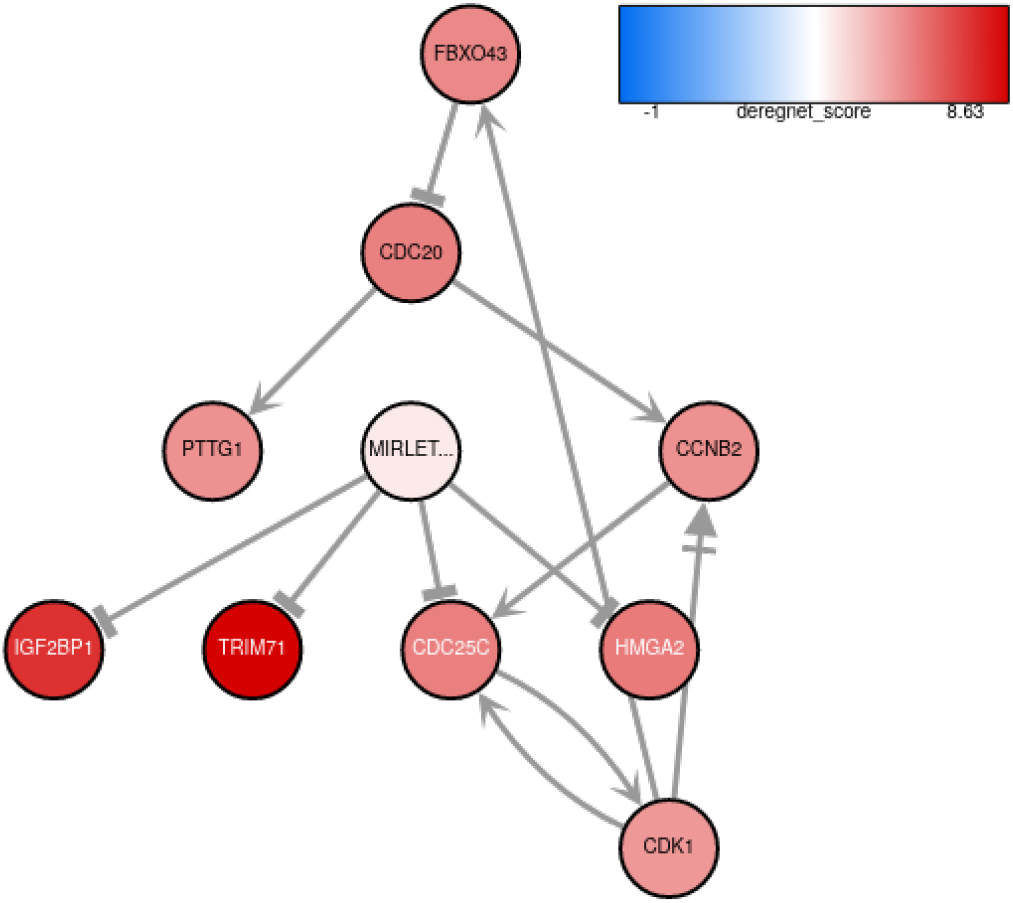
2^*nd*^ suboptimal upregulated global subgraph for TCGA-LIHC.

**Figure S 9:**
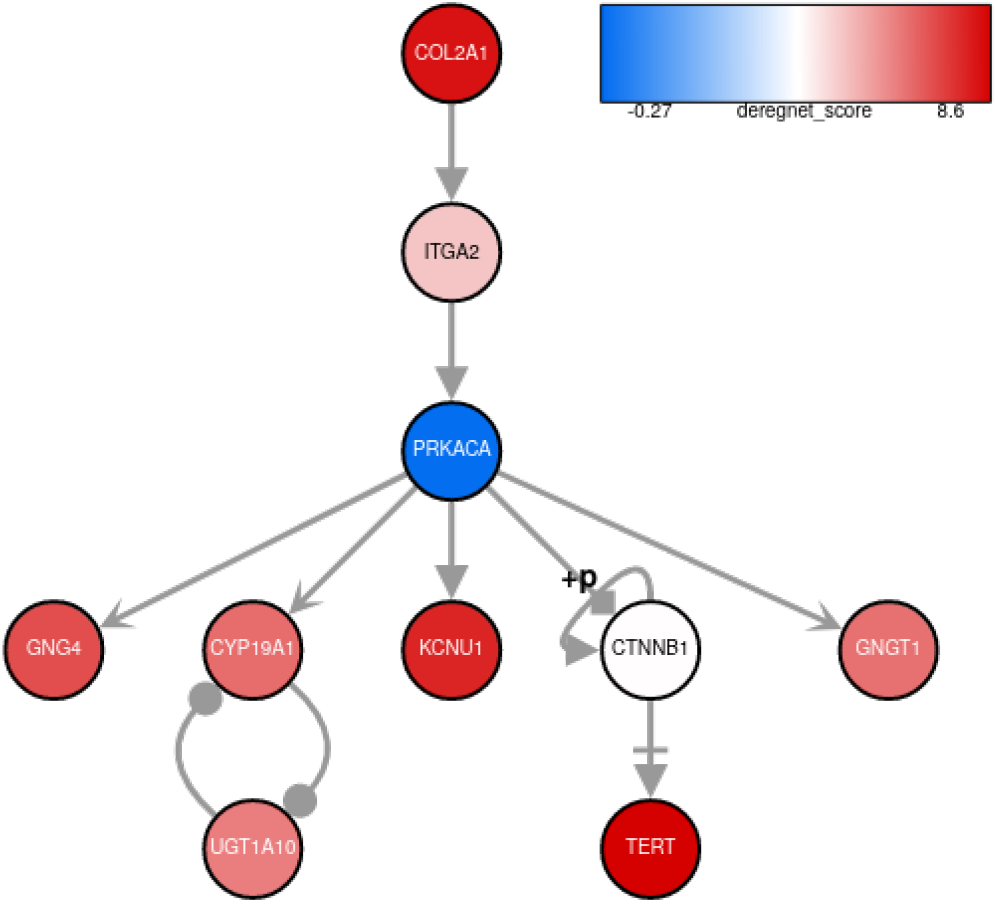
3^*rd*^ suboptimal upregulated global subgraph for TCGA-LIHC.

**Figure S 10:**
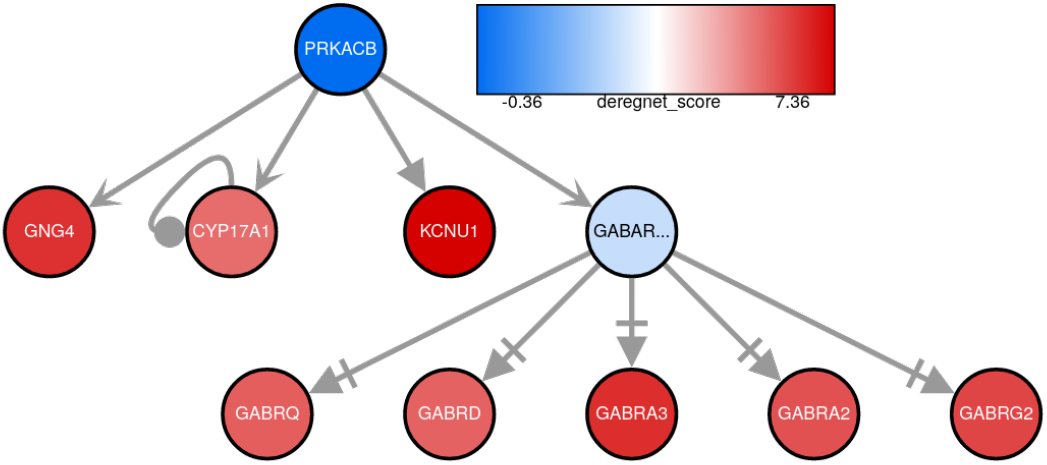
4^*th*^ suboptimal upregulated global subgraph for TCGA-LIHC.

**Figure S 11:**
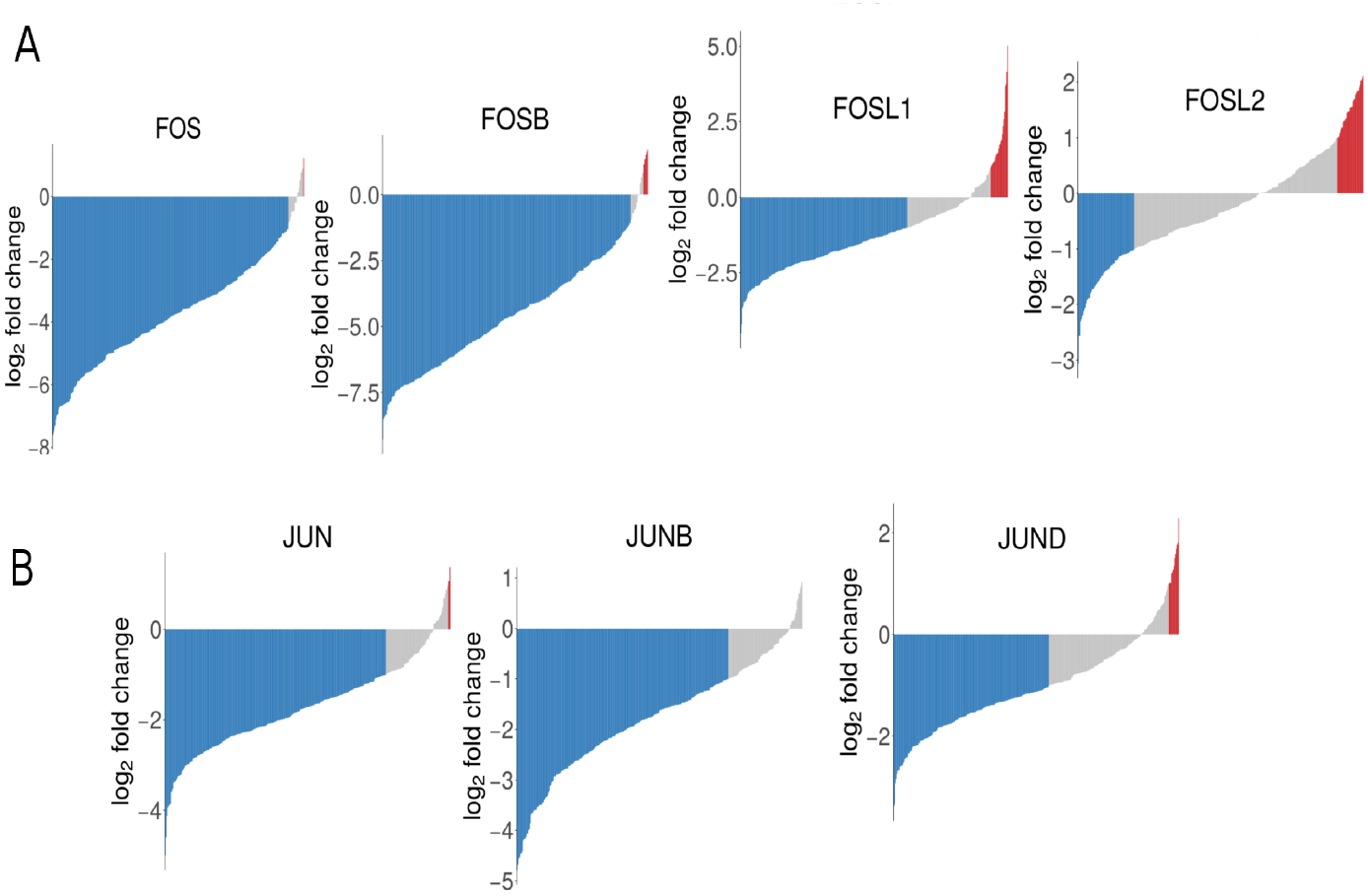
Expression of FOS and JUN isoforms in tumor of TCGA-LIHC cohort. (A) Log_2_-fold changes of FOS isoforms in individual tumors compared to the mean control value of the TCGA-LIHC dataset. (B) Log_2_ fold changes of JUN isoforms in individual tumors compared to the mean control value of the TCGA-LIHC dataset. Bars in waterfall plot indicate mRNA down-regulation ≥ 1.5-fold (blue),mRNA upregulation ≥ 1.5-fold (red).

**Figure S 12:**
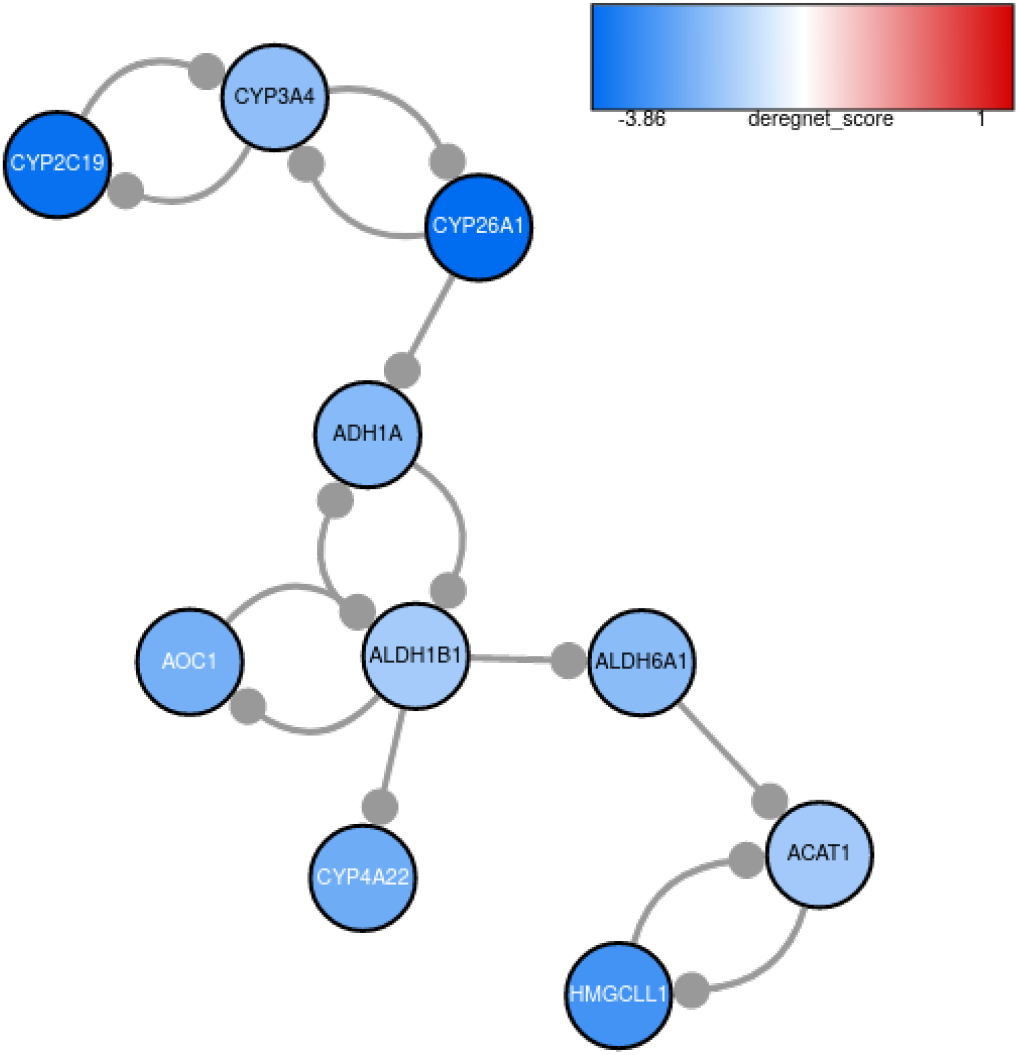
Optimal downregulated global subgraph for TCGA-LIHC.

**Figure S 13:**
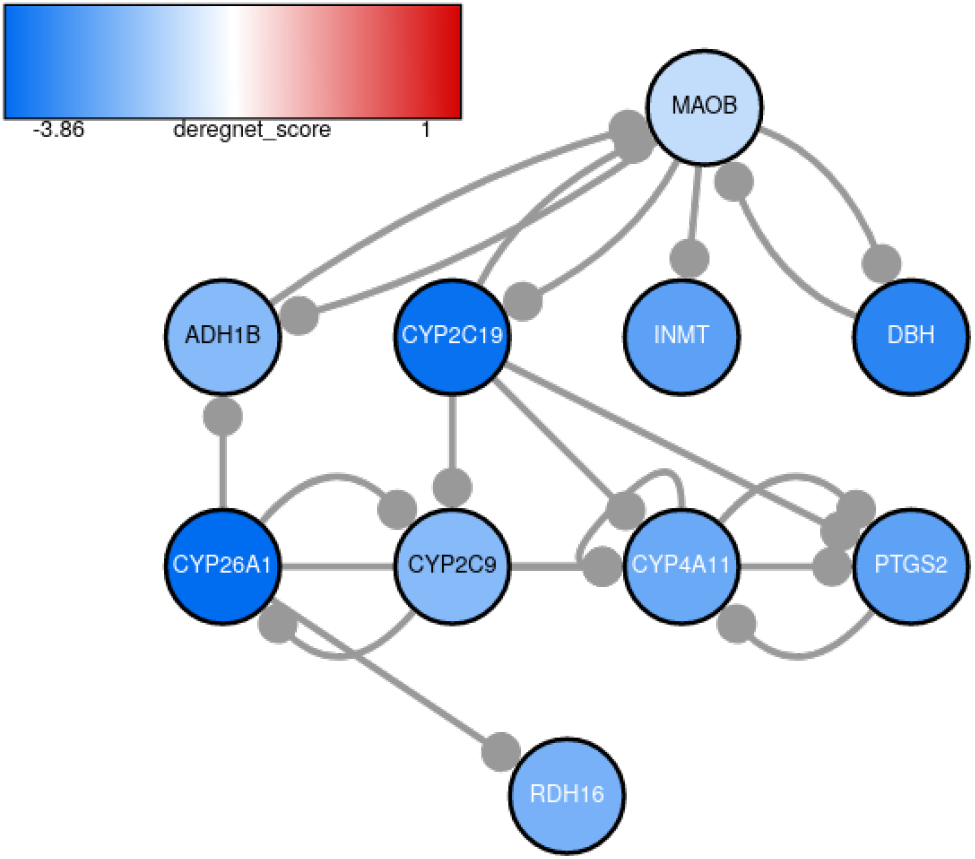
1^*st*^ suboptimal downregulated global subgraph for TCGA-LIHC.

**Figure S 14:**
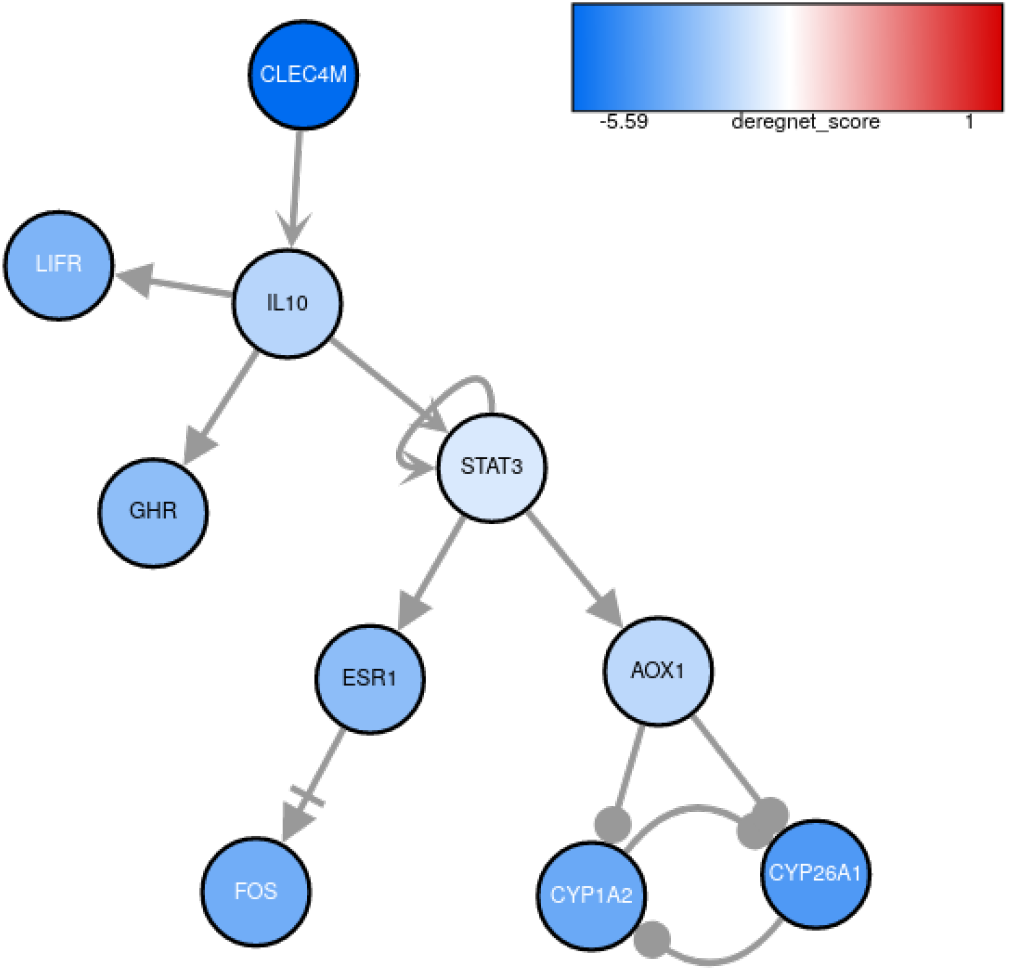
2^*nd*^ suboptimal downregulated global subgraph for TCGA-LIHC.

**Figure S 15:**
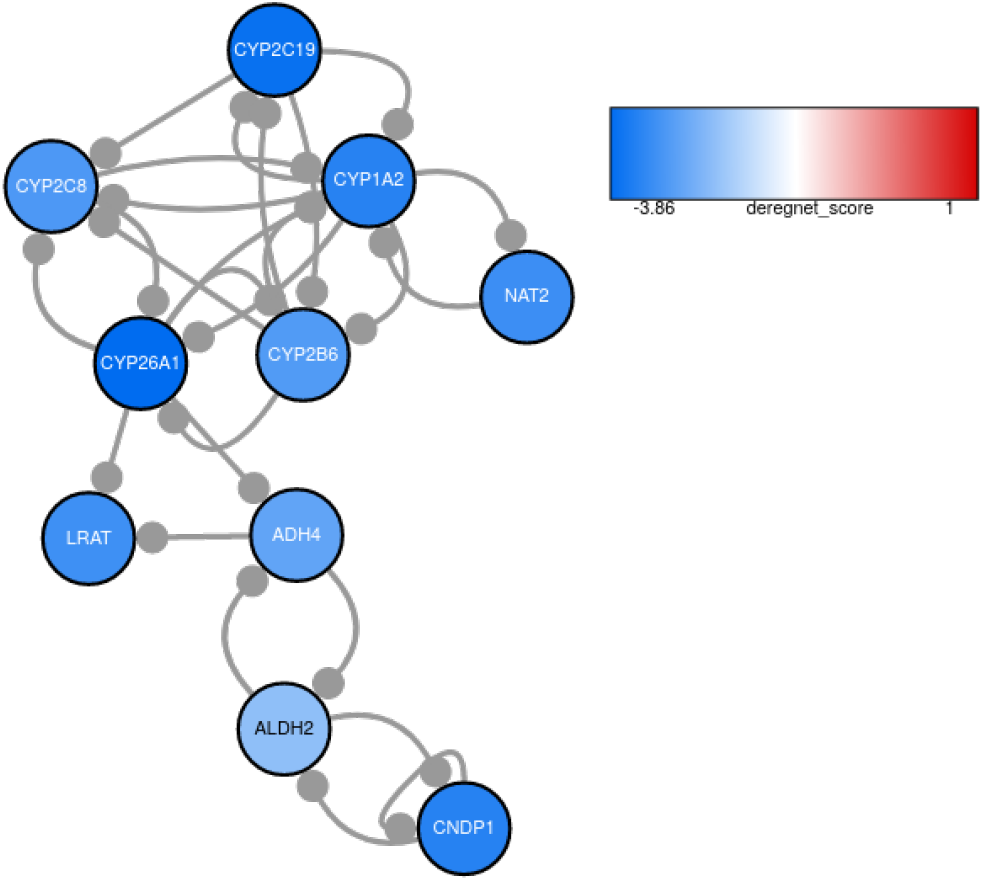
3^*rd*^ suboptimal downregulated global subgraph for TCGA-LIHC.

**Figure S 16:**
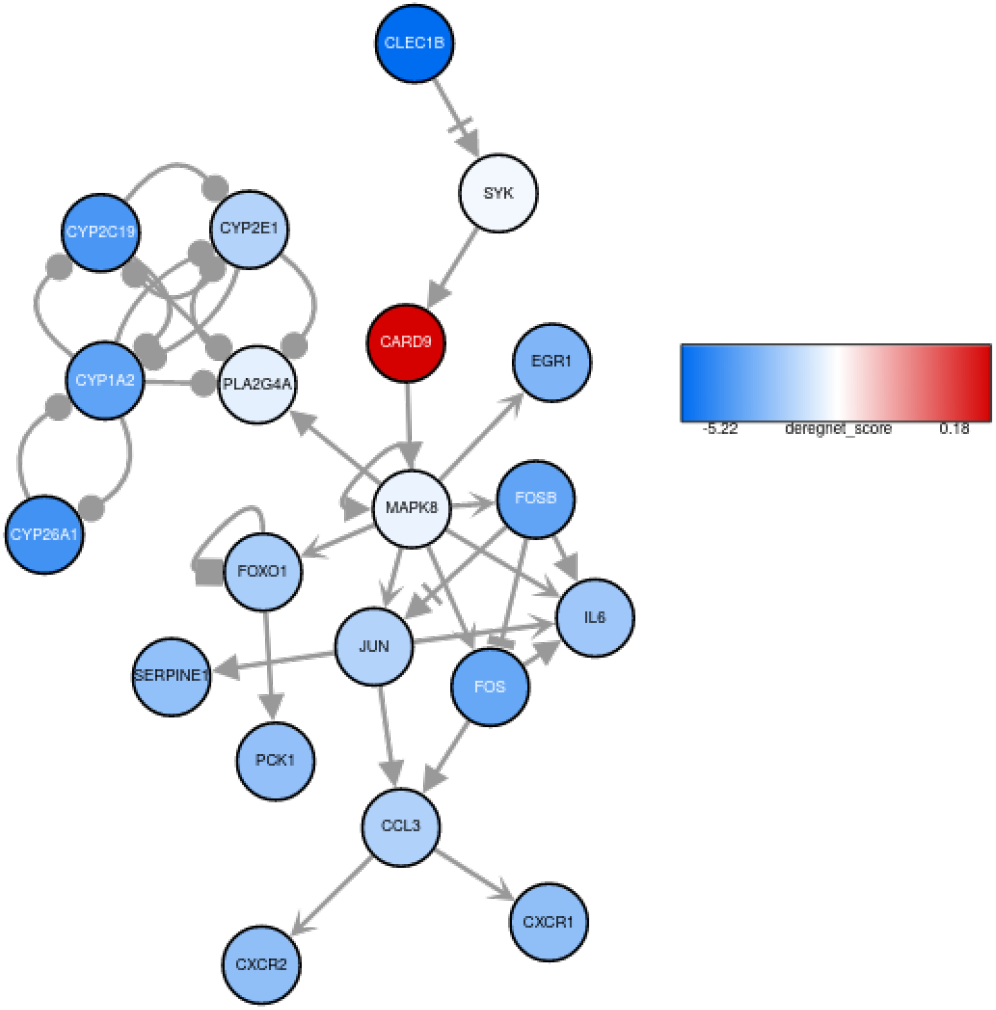
4^*th*^ suboptimal downregulated global subgraph for TCGA-LIHC.

**Figure S 17:**
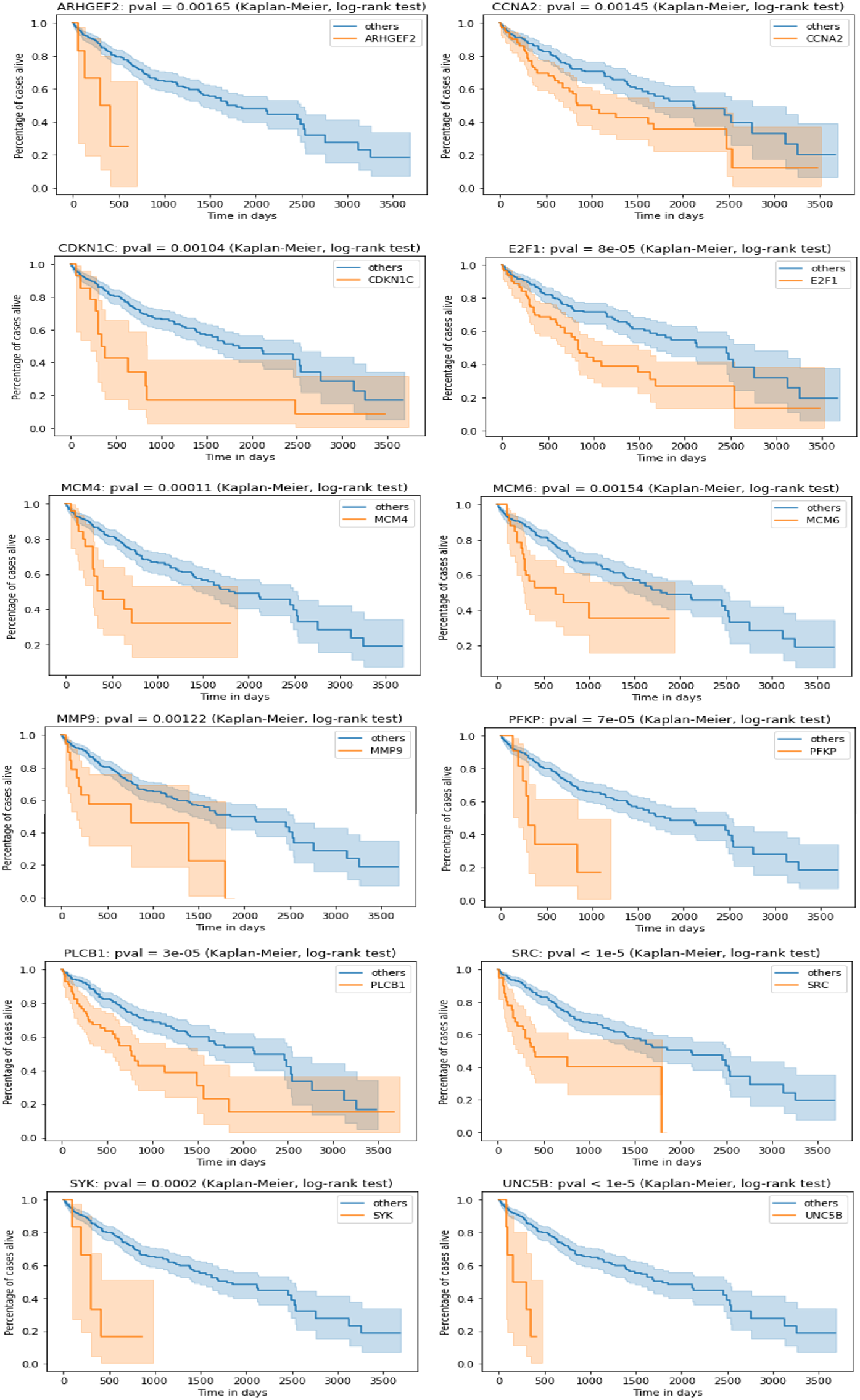
Subset of genes whose presence in a patient’s inferred subgraph indicates a poor survival. Survival difference is calculated using Kaplan-Meier estimates and log-rank test.

**Figure S 18:**
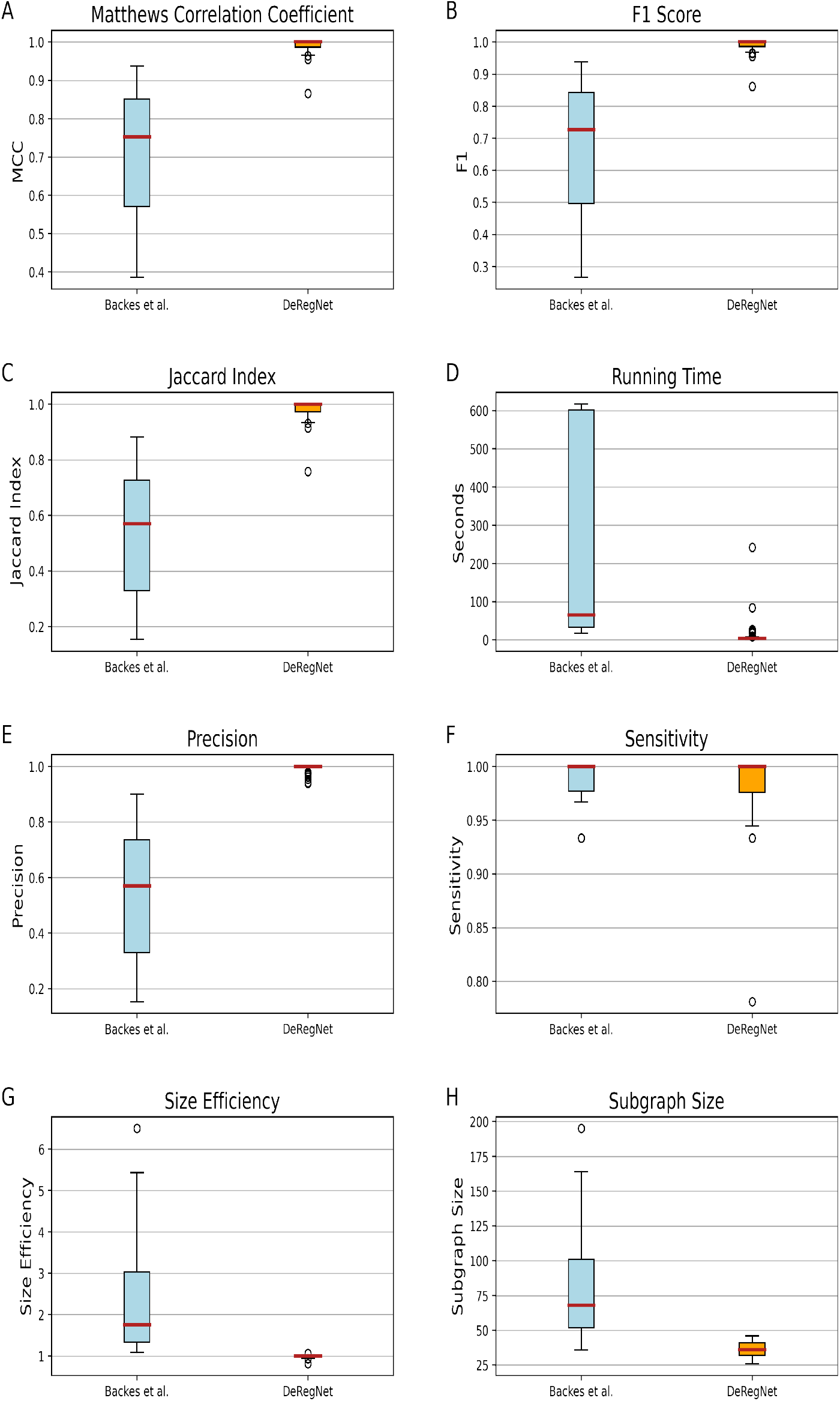
Benchmark results for out-of-subgraph deregulation probability. *p* = 0.0005. *k_min_* = 25, *k_max_* = 50, minimal size of simulated true subgraph = 30, maximal size of simulated true subgraph = 45, in-subgraph deregulation probability *p*′ = 0.99, number of simulated instances = 100, time limit = 600 seconds.

**Figure S 19:**
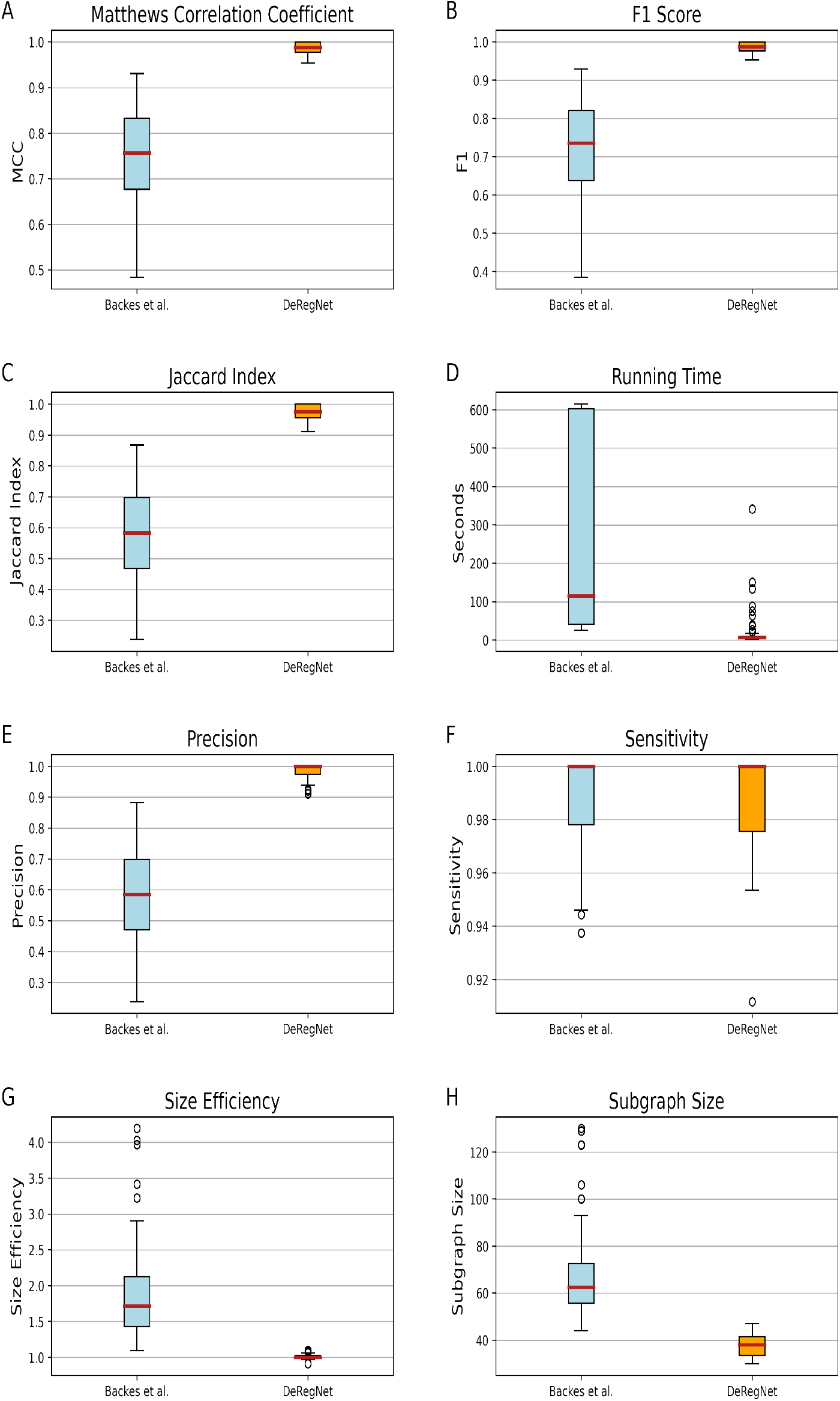
Benchmark results for out-of-subgraph deregulation probability. *p* = 0.001. *k_min_* = 25, *k_max_* = 50, minimal size of simulated true subgraph = 30, maximal size of simulated true subgraph = 45, in-subgraph deregulation probability *p*′ = 0.99, number of simulated instances = 100, time limit = 600 seconds.

**Figure S 20:**
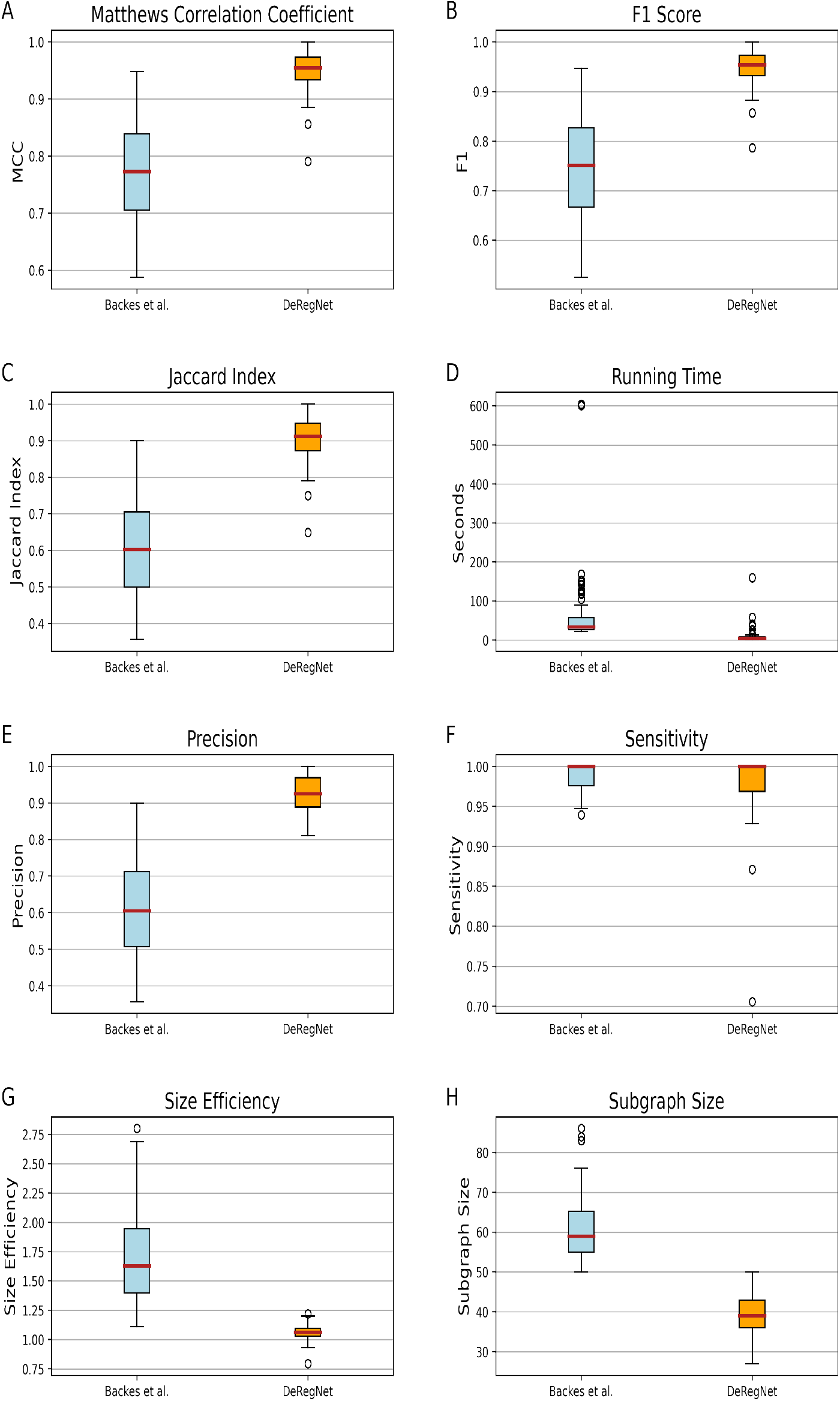
Benchmark results for out-of-subgraph deregulation probability. *p* = 0.005. *k_min_* = 25, *k_max_* = 50, minimal size of simulated true subgraph = 30, maximal size of simulated true subgraph = 45, in-subgraph deregulation probability *p*′ = 0.99, number of simulated instances = 100, time limit = 600 seconds.

**Figure S 21:**
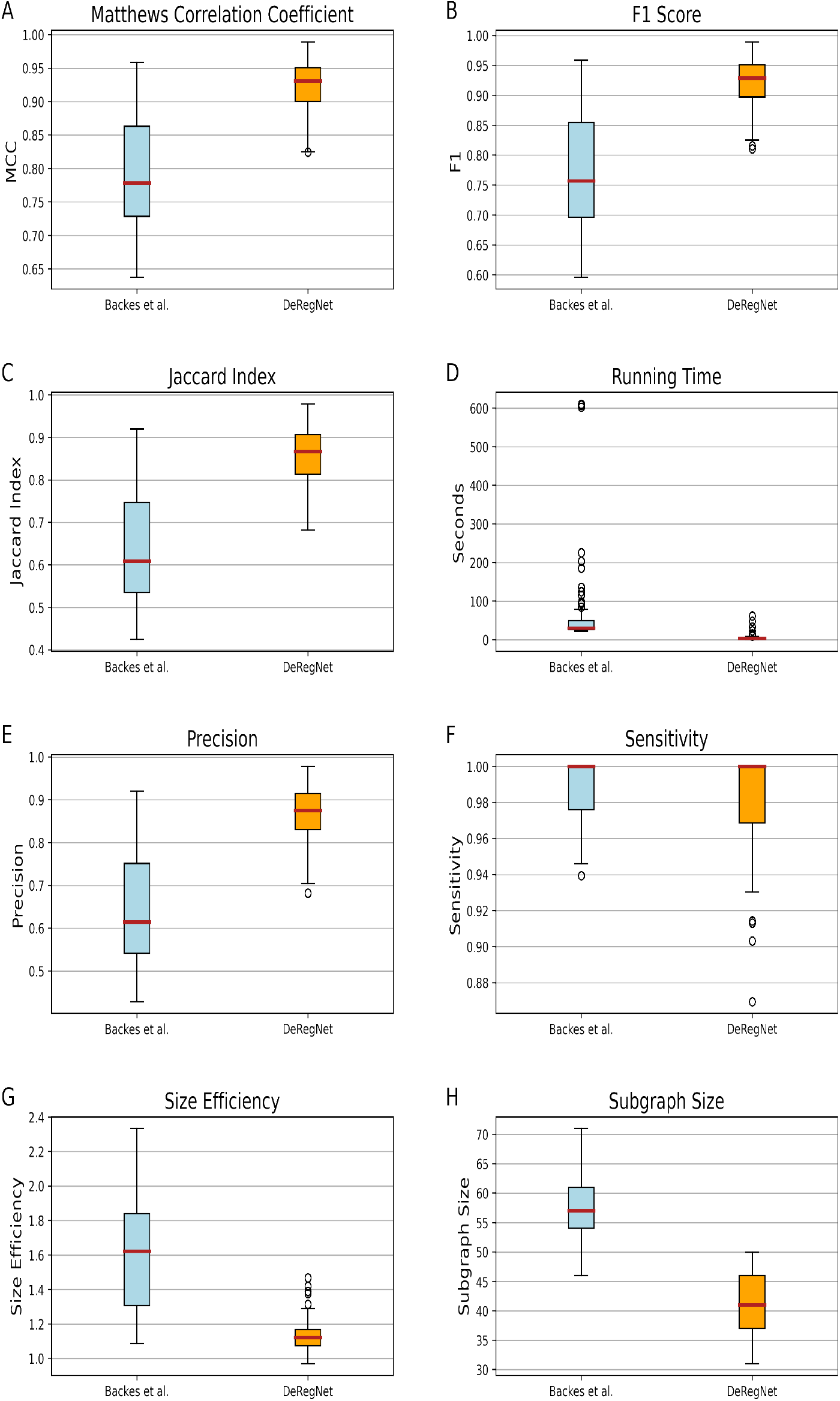
Benchmark results for out-of-subgraph deregulation probability. *p* = 0.01. *k_min_* = 25, *k_max_* = 50, minimal size of simulated true subgraph = 30, maximal size of simulated true subgraph = 45, in-subgraph deregulation probability *p*′ = 0.99, number of simulated instances = 100, time limit = 600 seconds.

**Figure S 22:**
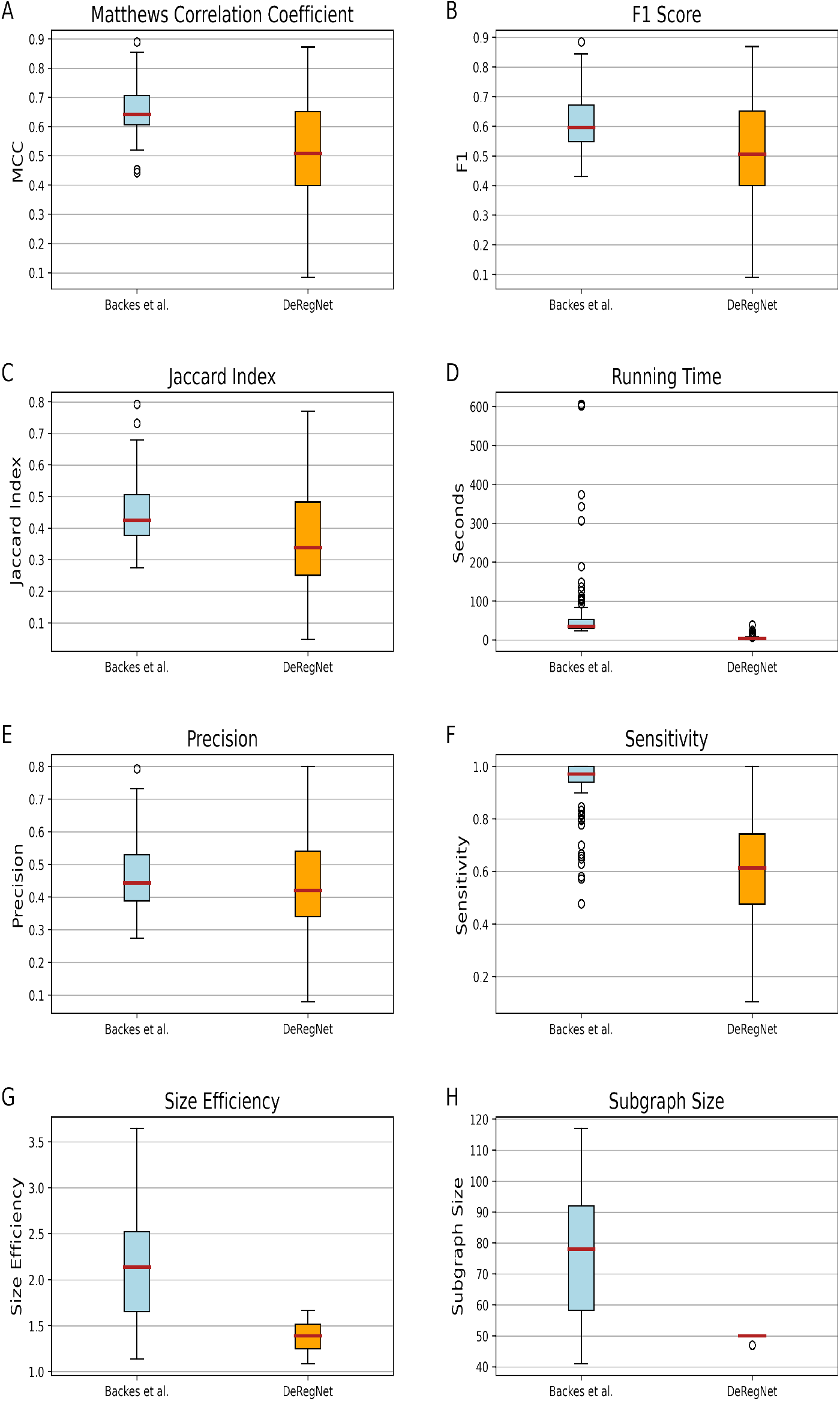
Benchmark results for out-of-subgraph deregulation probability. *p* = 0.05. *k_min_* = 25, *k_max_* = 50, minimal size of simulated true subgraph = 30, maximal size of simulated true subgraph = 45, in-subgraph deregulation probability *p*′ = 0.99, number of simulated instances = 100, time limit = 600 seconds.

**Figure S 23:**
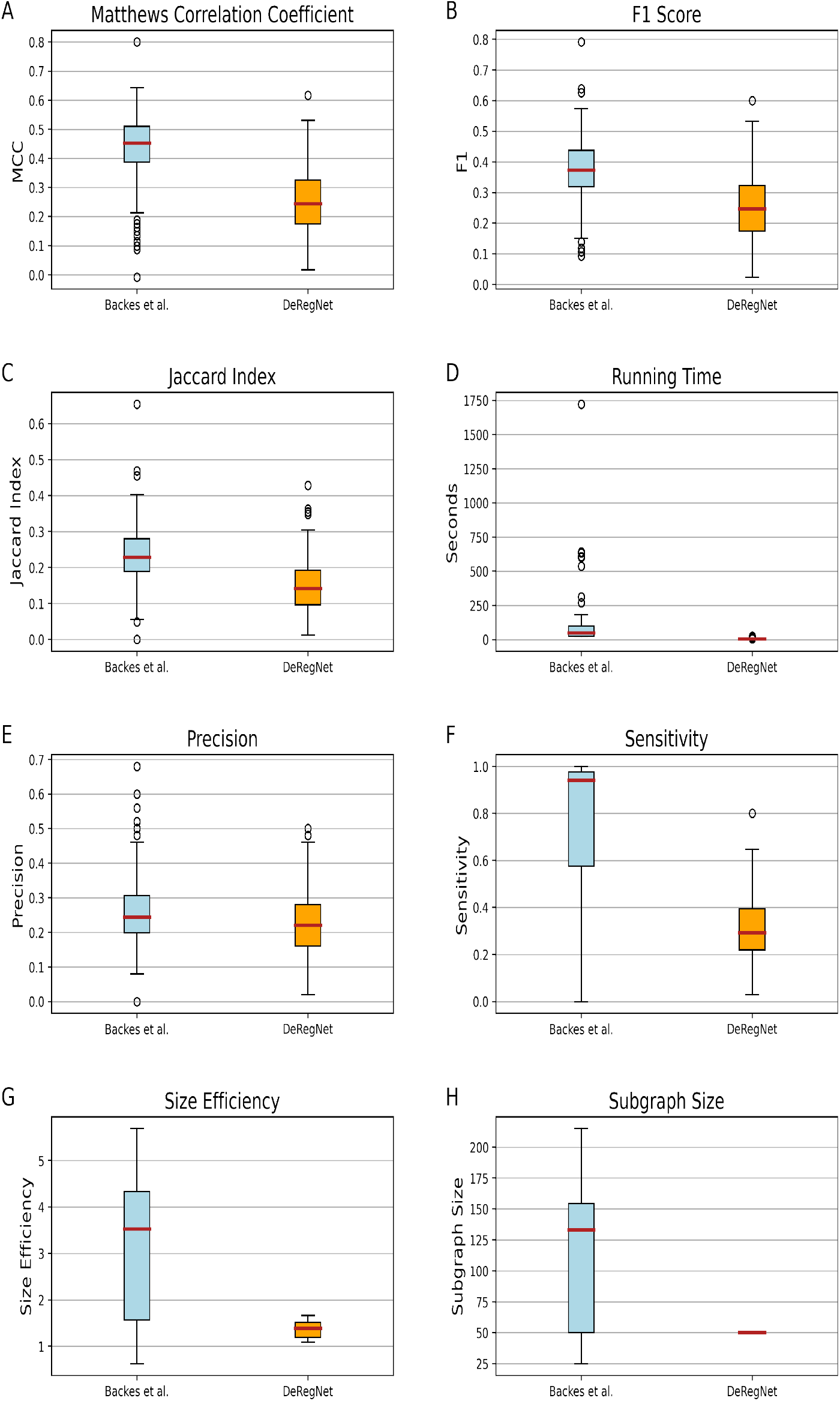
Benchmark results for out-of-subgraph deregulation probability. *p* = 0.1. *k_min_* = 25, *k_max_* = 50, minimal size of simulated true subgraph = 30, maximal size of simulated true subgraph = 45, in-subgraph deregulation probability *p*′ = 0.99, number of simulated instances = 100, time limit = 600 seconds.

**Figure S 24:**
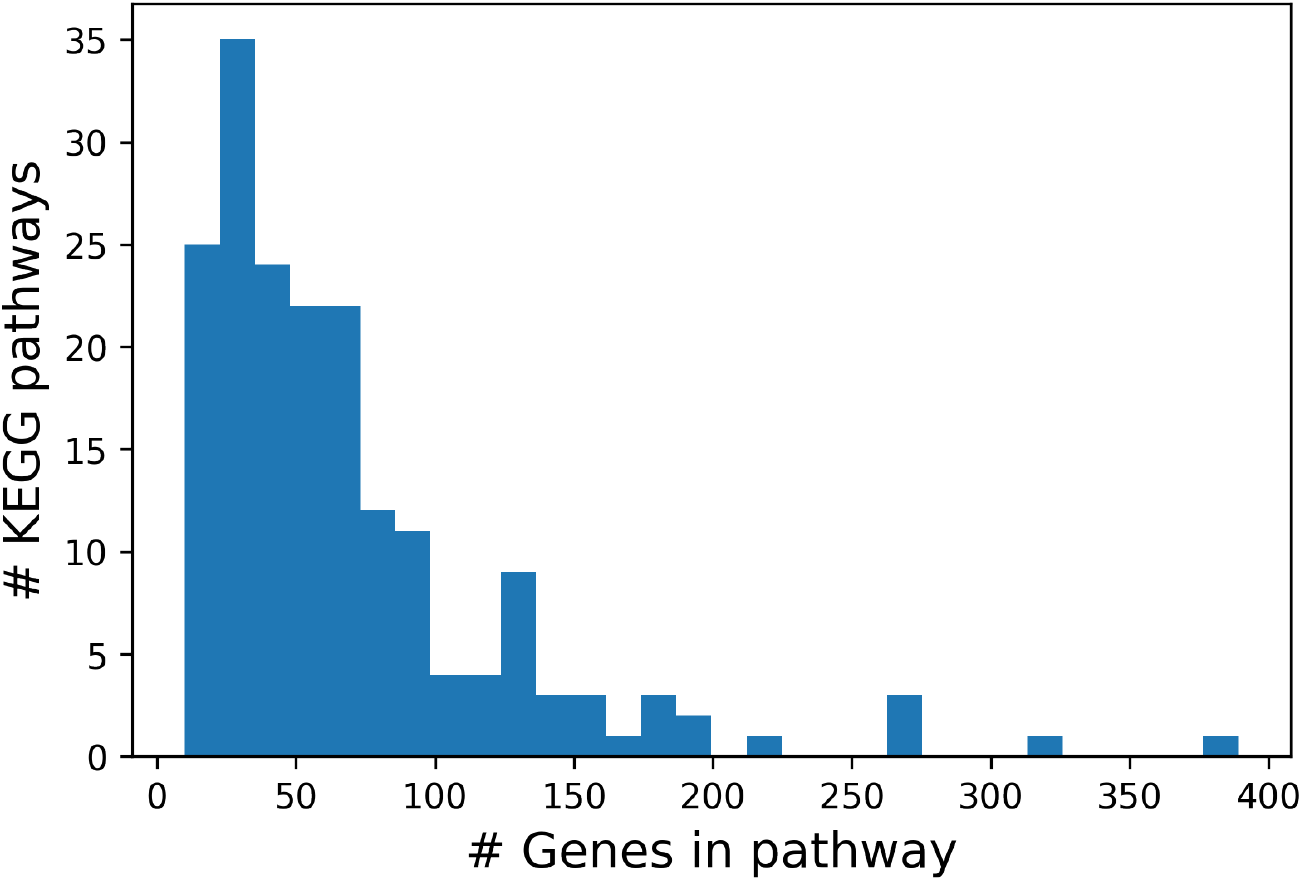
Distribution of KEGG pathway sizes. As can be seen from the distribution pre-defined KEGG pathways have a median size around 50. Most pathways with much higher number of genes are in fact meta pathways representing a considerable extent of biomolecular function like functions related to cancer across cancer types for example. Most pathways which are more narrow/specific in scope are smaller and hence one option to choose *k_min_* and *k_max_* is to aim for subnetworks e.g. between 10 to 60 in order to aim for reasonably sized subnetworks given the median size of common pre-defined pathways.

## References

1. Wang, Z., Gerstein, M., Snyder, M.: Rna-seq: a revolutionary tool for transcriptomics. Nat. Rev. Genet. 10, 57–63 (2009)

2. Altelaar, A.F.M., Munoz, J., Heck, A.J.R.: Next-generation proteomics: towards an integrative view of proteome dynamics. Nat. Rev. Genet. 14, 35–48 (2013)

3. Tomczak, K., Czerwi?ska, P., Wiznerowicz, M.: The Cancer Genome Atlas (TCGA): an immeasurable source of knowledge. Contemp Oncol (Pozn) 19(1A), 68–77 (2015)

4. Maciejewski, H.: Gene set analysis methods: statistical models and methodological differences. Brief. Bioinformatics 15(4), 504–518 (2014)

5. D’Eustachio, P.: Pathway databases: making chemical and biological sense of the genomic data flood. Chem. Biol. 20(5), 629–635 (2013)

6. Kanehisa, M., Furumichi, M., Tanabe, M., Sato, Y., Morishima, K.: KEGG: new perspectives on genomes, pathways, diseases and drugs. Nucleic Acids Res. 45(D1), 353–361 (2017)

7. Kutmon, M., Riutta, A., Nunes, N., Hanspers, K., Willighagen, E.L., Bohler, A., Melius, J., Waagmeester, A., Sinha, S.R., Miller, R., Coort, S.L., Cirillo, E., Smeets, B., Evelo, C.T., Pico, A.R.: WikiPathways: capturing the full diversity of pathway knowledge. Nucleic Acids Res. 44(D1), 488–494 (2016)

8. Fabregat, A., Jupe, S., Matthews, L., Sidiropoulos, K., Gillespie, M., Garapati, P., Haw, R., Jassal, B., Korninger, F., May, B., Milacic, M., Roca, C.D., Rothfels, K., Sevilla, C., Shamovsky, V., Shorser, S., Varusai, T., Viteri, G., Weiser, J., Wu, G., Stein, L., Hermjakob, H., D’Eustachio, P.: The Reactome Pathway Knowledgebase. Nucleic Acids Res. 46(D1), 649–655 (2018)

9. Subramanian, A., Tamayo, P., Mootha, V.K., Mukherjee, S., Ebert, B.L., Gillette, M.A., Paulovich, A., Pomeroy, S.L., Golub, T.R., Lander, E.S., Mesirov, J.P.: Gene set enrichment analysis: a knowledge-based approach for interpreting genome-wide expression profiles. Proc. Natl. Acad. Sci. U.S.A. 102(43), 15545–15550 (2005)

10. Caspi, R., Dreher, K., Karp, P.D.: The challenge of constructing, classifying, and representing metabolic pathways. FEMS Microbiol. Lett. 345(2), 85–93 (2013)

11. Biggin, M.D.: Animal transcription networks as highly connected, quantitative continua. Dev. Cell 21(4), 611–626 (2011)

12. Li, T., Wernersson, R., Hansen, R.B., Horn, H., Mercer, J., Slodkowicz, G., Workman, C.T., Rigina, O., Rapacki, K., St?rfeldt, H.H., Brunak, S., Jensen, T.S., Lage, K.: A scored human protein-protein interaction network to catalyze genomic interpretation. Nat. Methods 14(1), 61–64 (2017)

13. Szklarczyk, D., Morris, J.H., Cook, H., Kuhn, M., Wyder, S., Simonovic, M., Santos, A., Doncheva, N.T., Roth, A., Bork, P., Jensen, L.J., von Mering, C.: The STRING database in 2017: quality-controlled protein-protein association networks, made broadly accessible. Nucleic Acids Res. 45(D1), 362–368 (2017)

14. Jaakkola, M.K., Elo, L.L.: Empirical comparison of structure-based pathway methods. Brief. Bioinformatics 17(2), 336–345 (2016)

15. Mitrea, C., Taghavi, Z., Bokanizad, B., Hanoudi, S., Tagett, R., Donato, M., Voichita, C., Draghici, S.: Methods and approaches in the topology-based analysis of biological pathways. Front Physiol 4, 278 (2013)

16. Ihnatova, I., Popovici, V., Budinska, E.: A critical comparison of topology-based pathway analysis methods. PLoS ONE 13(1), 0191154 (2018)

17. Tarca, A.L., Draghici, S., Khatri, P., Hassan, S.S., Mittal, P., Kim, J.S., Kim, C.J., Kusanovic, J.P., Romero, R.: A novel signaling pathway impact analysis. Bioinformatics 25(1), 75–82 (2009)

18. Mitra, K., Carvunis, A.R., Ramesh, S.K., Ideker, T.: Integrative approaches for finding modular structure in biological networks. Nat. Rev. Genet. 14(10), 719–732 (2013)

19. Batra, R., Alcaraz, N., Gitzhofer, K., Pauling, J., Ditzel, H.J., Hellmuth, M., Baumbach, J., List, M.: On the performance of de novo pathway enrichment. NPJ Syst Biol Appl 3, 6 (2017)

20. Ideker, T., Ozier, O., Schwikowski, B., Siegel, A.F.: Discovering regulatory and signalling circuits in molecular interaction networks. Bioinformatics 18 Suppl 1, 233–240 (2002)

21. Patil, K.R., Nielsen, J.: Uncovering transcriptional regulation of metabolism by using metabolic network topology. Proc. Natl. Acad. Sci. U.S.A. 102(8), 2685–2689 (2005)

22. Ulitsky, I., Shamir, R.: Identification of functional modules using network topology and high-throughput data. BMC Syst Biol 1, 8 (2007)

23. Dittrich, M.T., Klau, G.W., Rosenwald, A., Dandekar, T., Muller, T.: Identifying functional modules in protein-protein interaction networks: an integrated exact approach. Bioinformatics 24(13), 223–231 (2008)

24. Zhao, X.M., Wang, R.S., Chen, L., Aihara, K.: Uncovering signal transduction networks from high-throughput data by integer linear programming. Nucleic Acids Res. 36(9), 48 (2008)

25. Ulitsky, I., Shamir, R.: Identifying functional modules using expression profiles and confidence-scored protein interactions. Bioinformatics 25(9), 1158–1164 (2009)

26. Ulitsky, I., Krishnamurthy, A., Karp, R.M., Shamir, R.: DEGAS: de novo discovery of dysregulated pathways in human diseases. PLoS ONE 5(10), 13367 (2010)

27. Dao, P., Wang, K., Collins, C., Ester, M., Lapuk, A., Sahinalp, S.C.: Optimally discriminative subnetwork markers predict response to chemotherapy. Bioinformatics 27(13), 205–213 (2011)

28. Bailly-Bechet, M., Borgs, C., Braunstein, A., Chayes, J., Dagkessamanskaia, A., Francois, J.M., Zecchina, R.: Finding undetected protein associations in cell signaling by belief propagation. Proc. Natl. Acad. Sci. U.S.A. 108(2), 882–887 (2011)

29. Alcaraz, N., Friedrich, T., Kotzing, T., Krohmer, A., Muller, J., Pauling, J., Baumbach, J.: Efficient key pathway mining: combining networks and OMICS data. Integr Biol (Camb) 4(7), 756–764 (2012)

30. Alcaraz, N., Pauling, J., Batra, R., Barbosa, E., Junge, A., Christensen, A.G., Azevedo, V., Ditzel, H.J., Baumbach, J.: KeyPathwayMiner 4.0: condition-specific pathway analysis by combining multiple omics studies and networks with Cytoscape. BMC Syst Biol 8, 99 (2014)

31. Alcaraz, N., List, M., Dissing-Hansen, M., Rehmsmeier, M., Tan, Q., Mollenhauer, J., Ditzel, H.J., Baumbach, J.: Robust de novo pathway enrichment with KeyPathwayMiner 5. F1000Res 5, 1531 (2016)

32. Vaske, C.J., Benz, S.C., Sanborn, J.Z., Earl, D., Szeto, C., Zhu, J., Haussler, D., Stuart, J.M.: Inference of patient-specific pathway activities from multi-dimensional cancer genomics data using PARADIGM. Bioinformatics 26(12), 237–245 (2010)

33. Vandin, F., Raphael, B.J., Upfal, E.: On the Sample Complexity of Cancer Pathways Identification. J. Comput. Biol. 23(1), 30–41 (2016)

34. Vandin, F., Upfal, E., Raphael, B.J.: De novo discovery of mutated driver pathways in cancer. Genome Res. 22(2), 375–385 (2012)

35. Zhang, J., Zhang, S.: The Discovery of Mutated Driver Pathways in Cancer: Models and Algorithms. IEEE/ACM Trans Comput Biol Bioinform 15(3), 988–998 (2018)

36. Cerami, E., Demir, E., Schultz, N., Taylor, B.S., Sander, C.: Automated network analysis identifies core pathways in glioblastoma. PLoS ONE 5(2), 8918 (2010)

37. Hofree, M., Shen, J.P., Carter, H., Gross, A., Ideker, T.: Network-based stratification of tumor mutations. Nat. Methods 10(11), 1108–1115 (2013)

38. Vandin, F., Upfal, E., Raphael, B.J.: Finding driver pathways in cancer: models and algorithms. Algorithms Mol Biol 7(1), 23 (2012)

39. Keller, A., Backes, C., Gerasch, A., Kaufmann, M., Kohlbacher, O., Meese, E., Lenhof, H.P.: A novel algorithm for detecting differentially regulated paths based on gene set enrichment analysis. Bioinformatics 25(21), 2787–2794 (2009)

40. Backes, C., Rurainski, A., Klau, G.W., Muller, O., Stockel, D., Gerasch, A., Kuntzer, J., Maisel, D., Ludwig, N., Hein, M., Keller, A., Burtscher, H., Kaufmann, M., Meese, E., Lenhof, H.P.: An integer linear programming approach for finding deregulated subgraphs in regulatory networks. Nucleic Acids Res. 40(6), 43 (2012)

41. Atias, N., Sharan, R.: iPoint: an integer programming based algorithm for inferring protein subnetworks. Mol Biosyst 9(7), 1662–1669 (2013)

42. Gaire, R.K., Smith, L., Humbert, P., Bailey, J., Stuckey, P.J., Haviv, I.: Discovery and analysis of consistent active sub-networks in cancers. BMC Bioinformatics 14 Suppl 2, 7 (2013)

43. Melas, I.N., Sakellaropoulos, T., Iorio, F., Alexopoulos, L.G., Loh, W.-Y., Lauffenburger, D.A., Saez-Rodriguez, J., Bai, J.P.F.: Identification of drug-specific pathways based on gene expression data: application to drug induced lung injury. Integrative Biology 7(8), 904–920 (2015). doi:10.1039/c4ib00294f. https://academic.oup.com/ib/article-pdf/7/8/904/27282429/c4ib00294f3.pdf

44. Liu, A., Trairatphisan, P., Gjerga, E., Didangelos, A., Barratt, J., Saez-Rodriguez, J.: From expression footprints to causal pathways: contextualizing large signaling networks with carnival. npj Systems Biology and Applications 5(1), 40 (2019). doi:10.1038/s41540-019-0118-z

45. Huang, S.S., Fraenkel, E.: Integrating proteomic, transcriptional, and interactome data reveals hidden components of signaling and regulatory networks. Sci Signal 2(81), 40 (2009)

46. Huang, S.S., Clarke, D.C., Gosline, S.J., Labadorf, A., Chouinard, C.R., Gordon, W., Lauffenburger, D.A., Fraenkel, E.: Linking proteomic and transcriptional data through the interactome and epigenome reveals a map of oncogene-induced signaling. PLoS Comput. Biol. 9(2), 1002887 (2013)

47. Tuncbag, N., Braunstein, A., Pagnani, A., Huang, S.S., Chayes, J., Borgs, C., Zecchina, R., Fraenkel, E.: Simultaneous reconstruction of multiple signaling pathways via the prize-collecting steiner forest problem. J. Comput. Biol. 20(2), 124–136 (2013)

48. Tuncbag, N., Gosline, S.J., Kedaigle, A., Soltis, A.R., Gitter, A., Fraenkel, E.: Network-Based Interpretation of Diverse High-Throughput Datasets through the Omics Integrator Software Package. PLoS Comput. Biol. 12(4), 1004879 (2016)

49. Charnes, A., Cooper, W.W.: Programming with linear fractional functionals. Naval Research Logistics Quaterly 9, 181–186 (1962)

50. Yue, D., Guillen-Gosalbez, G., You, F.: Global optimization of large-scale mixed-integer linear fractional programming problems: a reformulation-linearization method and process scheduling applications. AIChE Journal 59(11), 4255–4272 (2013)

51. Dinkelbach, W.: Die maximierung eines quotienten zweier linearer funktionen unter linearen nebenbedingungen. Z. Wahrscheinlichkeitstheorie 1, 141–145 (1962)

52. Dinkelbach, W.: On nonlinear fractional programming. Managment Science 13(7), 492–498 (1967)

53. Anzai, Y.: On integer fractional programming. J. Operations Research Soc. of Japan 17(1), 49–66 (1974)

54. You, F., Castro, P.M., Grossmann, I.E.: Dinkelbach’s algorithm as an efficient method to solve a class of minlp models for large-scale cyclic scheduling problems. Computers & Chemical Engineering 33, 1879–1889 (2009)

55. Glover, F.: Improved linear integer programming formulations of nonlinear integer problems. Managment Science 22(4), 455–460 (1975)

56. Adams, W.P., Forrester, R.J.: A simple recipe for concise mixed 0-1 linearizations. Operations Research Letters 33, 55–61 (2005)

57. Adams, W.P., Forrester, R.J., Glover, F.: Comparison and enhancement strategies for linearizing mixed 0-1 quadratic programs. Discrete Optimization 1, 99–120 (2004)

58. Sharir, M.: A strong-connectivity algorithm and its applications to data flow analysis. Computers and Mathematics with applications 7(1), 67–72 (1981)

59. Cerami, E.G., Gross, B.E., Demir, E., Rodchenkov, I., Babur, O., Anwar, N., Schultz, N., Bader, G.D., Sander, C.: Pathway Commons, a web resource for biological pathway data. Nucleic Acids Res. 39(Database issue), 685–690 (2011)

60. Zhang, J.D., Wiemann, S.: KEGGgraph: a graph approach to KEGG PATHWAY in R and bioconductor. Bioinformatics 25(11), 1470–1471 (2009)

61. Love, M.I., Huber, W., Anders, S.: Moderated estimation of fold change and dispersion for RNA-seq data with DESeq2. Genome Biol. 15(12), 550 (2014)

62. Touleimat, N., Tost, J.: Complete pipeline for infinium® human methylation 450k beadchip data processing using subset quantile normalization for accurate dna methylation estimation. Epigenomics 4(3), 325–341 (2012). doi:10.2217/epi.12.21. PMID: 22690668. https://doi.org/10.2217/epi.12.21

63. Wang, Z., Wu, X., Wang, Y.: A framework for analyzing dna methylation data from illumina infinium humanmethylation450 beadchip. BMC Bioinformatics 19(5), 115 (2018). doi:10.1186/s12859-018-2096-3

64. Kaplan, E.L., Meier, P.: Nonparametric estimation from incomplete observations. Journal of the American Statistical Association 53(282), 457–481 (1958). doi:10.1080/01621459.1958.10501452. https://www.tandfonline.com/doi/pdf/10.1080/01621459.1958.10501452

65. Aalen, O., Borgan, O., Gjessing, H.: Survival and Event History Analysis: A Process Point of View. Springer, ??? (2008)

66. Li, E., Zhang, Y.: DNA methylation in mammals. Cold Spring Harb Perspect Biol 6(5), 019133 (2014)

67. Ehrlich, M.: Dna hypomethylation in cancer cells. Epigenomics 1(2), 239–259 (2009). doi:10.2217/epi.09.33.20495664[pmid]

68. Arzumanyan, A., Reis, H.M., Feitelson, M.A.: Pathogenic mechanisms in HBV- and HCV-associated hepatocellular carcinoma. Nat. Rev. Cancer 13(2), 123–135 (2013)

69. Zhou, F., Shang, W., Yu, X., Tian, J.: Glypican-3: A promising biomarker for hepatocellular carcinoma diagnosis and treatment. Med Res Rev 38(2), 741–767 (2018)

70. Wu, Y., Liu, H., Ding, H.: GPC-3 in hepatocellular carcinoma: current perspectives. J Hepatocell Carcinoma 3, 63–67 (2016)

71. Feng, M., Ho, M.: Glypican-3 antibodies: a new therapeutic target for liver cancer. FEBS Lett. 588(2), 377–382 (2014)

72. Filmus, J., Capurro, M.: Glypican-3: a marker and a therapeutic target in hepatocellular carcinoma. FEBS J. 280(10), 2471–2476 (2013)

73. Ho, M., Kim, H.: Glypican-3: a new target for cancer immunotherapy. Eur. J. Cancer 47(3), 333–338 (2011)

74. Bertino, G., Ardiri, A., Malaguarnera, M., Malaguarnera, G., Bertino, N., Calvagno, G.S.: Hepatocellualar carcinoma serum markers. Semin. Oncol. 39(4), 410–433 (2012)

75. Llovet, J.M., Zucman-Rossi, J., Pikarsky, E., Sangro, B., Schwartz, M., Sherman, M., Gores, G.: Hepatocellular carcinoma. Nat Rev Dis Primers 2, 16018 (2016)

76. Nault, J.C., Zucman-Rossi, J.: TERT promoter mutations in primary liver tumors. Clin Res Hepatol Gastroenterol 40(1), 9–14 (2016)

77. Quaas, A., Oldopp, T., Tharun, L., Klingenfeld, C., Krech, T., Sauter, G., Grob, T.J.: Frequency of TERT promoter mutations in primary tumors of the liver. Virchows Arch. 465(6), 673–677 (2014)

78. Totoki, Y., Tatsuno, K., Covington, K.R., Ueda, H., Creighton, C.J., Kato, M., Tsuji, S., Donehower, L.A., Slagle, B.L., Nakamura, H., Yamamoto, S., Shinbrot, E., Hama, N., Lehmkuhl, M., Hosoda, F., Arai, Y., Walker, K., Dahdouli, M., Gotoh, K., Nagae, G., Gingras, M.C., Muzny, D.M., Ojima, H., Shimada, K., Midorikawa, Y., Goss, J.A., Cotton, R., Hayashi, A., Shibahara, J., Ishikawa, S., Guiteau, J., Tanaka, M., Urushidate, T., Ohashi, S., Okada, N., Doddapaneni, H., Wang, M., Zhu, Y., Dinh, H., Okusaka, T., Kokudo, N., Kosuge, T., Takayama, T., Fukayama, M., Gibbs, R.A., Wheeler, D.A., Aburatani, H., Shibata, T.: Trans-ancestry mutational landscape of hepatocellular carcinoma genomes. Nat. Genet. 46(12), 1267–1273 (2014)

79. Anastas, J.N., Moon, R.T.: WNT signalling pathways as therapeutic targets in cancer. Nat. Rev. Cancer 13(1), 11–26 (2013)

80. Sun, C., Sun, L., Li, Y., Kang, X., Zhang, S., Liu, Y.: Sox2 expression predicts poor survival of hepatocellular carcinoma patients and it promotes liver cancer cell invasion by activating Slug. Med. Oncol. 30(2), 503 (2013)

81. Wen, W., Han, T., Chen, C., Huang, L., Sun, W., Wang, X., Chen, S.Z., Xiang, D.M., Tang, L., Cao, D., Feng, G.S., Wu, M.C., Ding, J., Wang, H.Y.: Cyclin G1 expands liver tumor-initiating cells by Sox2 induction via Akt/mTOR signaling. Mol. Cancer Ther. 12(9), 1796–1804 (2013)

82. Liu, L., Liu, C., Zhang, Q., Shen, J., Zhang, H., Shan, J., Duan, G., Guo, D., Chen, X., Cheng, J., Xu, Y., Yang, Z., Yao, C., Lai, M., Qian, C.: SIRT1-mediated transcriptional regulation of SOX2 is important for self-renewal of liver cancer stem cells. Hepatology 64(3), 814–827 (2016)

83. Min, L., Ji, Y., Bakiri, L., Qiu, Z., Cen, J., Chen, X., Chen, L., Scheuch, H., Zheng, H., Qin, L., Zatloukal, K., Hui, L., Wagner, E.F.: Liver cancer initiation is controlled by AP-1 through SIRT6-dependent inhibition of survivin. Nat. Cell Biol. 14(11), 1203–1211 (2012)

84. Montorsi, M., Maggioni, M., Falleni, M., Pellegrini, C., Donadon, M., Torzilli, G., Santambrogio, R., Spinelli, A., Coggi, G., Bosari, S.: Survivin gene expression in chronic liver disease and hepatocellular carcinoma. Hepatogastroenterology 54(79), 2040–2044 (2007)

85. Su, C.: Survivin in survival of hepatocellular carcinoma. Cancer Lett. 379(2), 184–190 (2016)

86. Takigawa, Y., Brown, A.M.: Wnt signaling in liver cancer. Curr Drug Targets 9(11), 1013–1024 (2008)

87. Liu, L.J., Xie, S.X., Chen, Y.T., Xue, J.L., Zhang, C.J., Zhu, F.: Aberrant regulation of Wnt signaling in hepatocellular carcinoma. World J. Gastroenterol. 22(33), 7486–7499 (2016)

88. Vilchez, V., Turcios, L., Marti, F., Gedaly, R.: Targeting Wnt/î^2^-catenin pathway in hepatocellular carcinoma treatment. World J. Gastroenterol. 22(2), 823–832 (2016)

89. Clevers, H., Nusse, R.: Wnt/î^2^-catenin signaling and disease. Cell 149(6), 1192–1205 (2012)

90. Nusse, R., Clevers, H.: Wnt/P-Catenin Signaling, Disease, and Emerging Therapeutic Modalities. Cell 169(6), 985–999 (2017)

91. Bellahcene, A., Castronovo, V., Ogbureke, K.U., Fisher, L.W., Fedarko, N.S.: Small integrin-binding ligand N-linked glycoproteins (SIBLINGs): multifunctional proteins in cancer. Nat. Rev. Cancer 8(3), 212–226 (2008)

92. Wen, Y., Jeong, S., Xia, Q., Kong, X.: Role of Osteopontin in Liver Diseases. Int. J. Biol. Sci. 12(9), 1121–1128 (2016)

93. Karni, R., Gus, Y., Dor, Y., Meyuhas, O., Levitzki, A.: Active Src elevates the expression of beta-catenin by enhancement of cap-dependent translation. Mol. Cell. Biol. 25(12), 5031–5039 (2005)

94. Eferl, R., Wagner, E.F.: AP-1: a double-edged sword in tumorigenesis. Nat. Rev. Cancer 3(11), 859–868 (2003)

95. Luch, A.: Nature and nurture - lessons from chemical carcinogenesis. Nat. Rev. Cancer 5(2), 113–125 (2005)

96. Undevia, S.D., Gomez-Abuin, G., Ratain, M.J.: Pharmacokinetic variability of anticancer agents. Nat. Rev. Cancer 5(6), 447–458 (2005)

97. Lowell, C.A.: Src-family and Syk kinases in activating and inhibitory pathways in innate immune cells: signaling cross talk. Cold Spring Harb Perspect Biol 3(3) (2011)

98. Krisenko, M.O., Geahlen, R.L.: Calling in SYK: SYK’s dual role as a tumor promoter and tumor suppressor in cancer. Biochim. Biophys. Acta 1853(1), 254–263 (2015)

99. Mocsai, A., Ruland, J., Tybulewicz, V.L.: The SYK tyrosine kinase: a crucial player in diverse biological functions. Nat. Rev. Immunol. 10(6), 387–402 (2010)

100. Hong, J., Yuan, Y., Wang, J., Liao, Y., Zou, R., Zhu, C., Li, B., Liang, Y., Huang, P., Wang, Z., Lin, W., Zeng, Y., Dai, J.L., Chung, R.T.: Expression of variant isoforms of the tyrosine kinase SYK determines the prognosis of hepatocellular carcinoma. Cancer Res. 74(6), 1845–1856 (2014)

101. Shin, S.H., Lee, K.H., Kim, B.H., Lee, S., Lee, H.S., Jang, J.J., Kang, G.H.: Downregulation of spleen tyrosine kinase in hepatocellular carcinoma by promoter CpG island hypermethylation and its potential role in carcinogenesis. Lab. Invest. 94(12), 1396–1405 (2014)

102. Hong, J., Hu, K., Yuan, Y., Sang, Y., Bu, Q., Chen, G., Yang, L., Li, B., Huang, P., Chen, D., Liang, Y., Zhang, R., Pan, J., Zeng, Y.X., Kang, T.: CHK1 targets spleen tyrosine kinase (L) for proteolysis in hepatocellular carcinoma. J. Clin. Invest. 122(6), 2165–2175 (2012)

103. Qu, C., Zheng, D., Li, S., Liu, Y., Lidofsky, A., Holmes, J.A., Chen, J., He, L., Wei, L., Liao, Y., Yuan, H., Jin, Q., Lin, Z., Hu, Q., Jiang, Y., Tu, M., Chen, X., Li, W., Lin, W., Fuchs, B.C., Chung, R.T., Hong, J.: Tyrosine kinase SYK is a potential therapeutic target for liver fibrosis. Hepatology (2018)

104. Bataller, R., Brenner, D.A.: Liver fibrosis. J. Clin. Invest. 115(2), 209–218 (2005)

105. Thorpe, L.M., Yuzugullu, H., Zhao, J.J.: PI3K in cancer: divergent roles of isoforms, modes of activation and therapeutic targeting. Nat. Rev. Cancer 15(1), 7–24 (2015)

106. Uen, Y.H., Fang, C.L., Hseu, Y.C., Shen, P.C., Yang, H.L., Wen, K.S., Hung, S.T., Wang, L.H., Lin, K.Y.: VAV3 oncogene expression in colorectal cancer: clinical aspects and functional characterization. Sci Rep 5, 9360 (2015)

107. Citterio, C., Menacho-Marquez, M., Garcia-Escudero, R., Larive, R.M., Barreiro, O., Sanchez-Madrid, F., Paramio, J.M., Bustelo, X.R.: The rho exchange factors vav2 and vav3 control a lung metastasis-specific transcriptional program in breast cancer cells. Sci Signal 5(244), 71 (2012)

108. Chen, X., Chen, S.I., Liu, X.A., Zhou, W.B., Ma, R.R., Chen, L.: Vav3 oncogene is upregulated and a poor prognostic factor in breast cancer patients. Oncol Lett 9(5), 2143–2148 (2015)

109. Li, X., Xu, W., Kang, W., Wong, S.H., Wang, M., Zhou, Y., Fang, X., Zhang, X., Yang, H., Wong, C.H., To, K.F., Chan, S.L., Chan, M.T.V., Sung, J.J.Y., Wu, W.K.K., Yu, J.: Genomic analysis of liver cancer unveils novel driver genes and distinct prognostic features. Theranostics 8(6), 1740–1751 (2018)

110. Roussos, E.T., Condeelis, J.S., Patsialou, A.: Chemotaxis in cancer. Nat. Rev. Cancer 11(8), 573–587 (2011)

111. Hardwick, J.M., Soane, L.: Multiple functions of BCL-2 family proteins. Cold Spring Harb Perspect Biol 5(2) (2013)

112. Mandriota, S.J., Jussila, L., Jeltsch, M., Compagni, A., Baetens, D., Prevo, R., Banerji, S., Huarte, J., Montesano, R., Jackson, D.G., Orci, L., Alitalo, K., Christofori, G., Pepper, M.S.: Vascular endothelial growth factor-C-mediated lymphangiogenesis promotes tumour metastasis. EMBO J. 20(4), 672–682 (2001)

113. Tammela, T., Zarkada, G., Wallgard, E., Murtomaki, A., Suchting, S., Wirzenius, M., Waltari, M., Hellstrom, M., Schomber, T., Peltonen, R., Freitas, C., Duarte, A., Isoniemi, H., Laakkonen, P., Christofori, G., Yla-Herttuala, S., Shibuya, M., Pytowski, B., Eichmann, A., Betsholtz, C., Alitalo, K.: Blocking VEGFR-3 suppresses angiogenic sprouting and vascular network formation. Nature 454(7204), 656–660 (2008)

114. Tvorogov, D., Anisimov, A., Zheng, W., Leppanen, V.M., Tammela, T., Laurinavicius, S., Holnthoner, W., Helotera, H., Holopainen, T., Jeltsch, M., Kalkkinen, N., Lankinen, H., Ojala, P.M., Alitalo, K.: Effective suppression of vascular network formation by combination of antibodies blocking VEGFR ligand binding and receptor dimerization. Cancer Cell 18(6), 630–640 (2010)

115. Skålhegg, B.S., Task’n, K.: Specificity in the cAMP/PKA signaling pathway. differential expression, regulation, and subcellular localization of subunits of PKA. Front Biosci 2, 331–342 (1997)

## References

1. Backes, C., Rurainski, A., Klau, G.W., Muller, O., Stockel, D., Gerasch, A., Kuntzer, J., Maisel, D., Ludwig, N., Hein, M., Keller, A., Burtscher, H., Kaufmann, M., Meese, E., Lenhof, H.P.: An integer linear programming approach for finding deregulated subgraphs in regulatory networks. Nucleic Acids Res. 40(6), 43 (2012)

2. Dittrich, M.T., Klau, G.W., Rosenwald, A., Dandekar, T., Muller, T.: Identifying functional modules in protein-protein interaction networks: an integrated exact approach. Bioinformatics 24(13), 223–231 (2008)

3. Buchanan, A., Wang, Y., Butenko, S.: Algorithms for node-weighted steiner tree and maximum-weight connected subgraph. Networks 72 (2017). doi:10.1002/net.21825

4. Loboda, A.A., Artyomov, M.N., Sergushichev, A.A.: Solving generalized maximum-weight connected subgraph problem for network enrichment analysis. In: Frith, M., Storm Pedersen, C.N. (eds.) Algorithms in Bioinformatics, pp. 210–221. Springer, Cham (2016)

## References

5. EI-Kebir, M., Klau, G.: Solving the maximum-weight connected subgraph problem to optimality. 11th DIMACS implementation challenge (2014)

6. Alvarez-Miranda, E., Ljubic, I., Mutzel, P.: The Maximum Weight Connected Subgraph Problem. In: Juenger, M., Reinelt, G. (eds.) The Maximum Weight Connected Subgraph Problem, pp. 245–270. Springer, Berlin, Heidelberg (2013)

7. Álvarez-Miranda, E., Ljubić, I., Mutzel, P.: The rooted maximum node-weight connected subgraph problem. In: Gomes, C., Sellmann, M. (eds.) Integration of AI and OR Techniques in Constraint Programming for Combinatorial Optimization Problems, pp. 300–315. Springer, Berlin, Heidelberg (2013)

8. Althaus, E., Blumenstock, M.: Algorithms for the maximum weight connected subgraph and prize-collecting steiner tree problems. 11th DIMACS Implementation Challenge in Collaboration with ICERM (2011)

9. Álvarez-Miranda, E., Sinnl, M.: A relax-and-cut framework for large-scale maximum weight connected subgraph problems. Computers & Operations Research 87, 63–82 (2017). doi:10.1016/j.cor.2017.05.015

10. Rehfeldt, D., Koch, T., Maher, S.J.: Reduction techniques for the prize collecting steiner tree problem and the maximum-weight connected subgraph problem. Networks 73(2), 206–233 (2019). doi:10.1002/net.21857. https://onlinelibrary.wiley.com/doi/pdf/10.1002/net.21857

11. Rehfeldt, D., Koch, T.: Combining np-hard reduction techniques and strong heuristics in an exact algorithm for the maximum-weight connected subgraph problem. SIAM Journal on Optimization 29(1), 369–398 (2019). doi:10.1137/17M1145963. https://doi.org/10.1137/17M1145963

12. You, F., Castro, P.M., Grossmann, I.E.: Dinkelbach’s algorithm as an efficient method to solve a class of minlp models for large-scale cyclic scheduling problems. Computers & Chemical Engineering 33, 1879–1889 (2009)

13. Yue, D., Guillén-Gosálbez, G., You, F.: Global optimization of large-scale mixed-integer linear fractional programming problems: a reformulation-linearization method and process scheduling applications. AIChE Journal 59(11), 4255–4272 (2013)

14. Charnes, A., Cooper, W.W.: Programming with linear fractional functionals. Naval Research Logistics Quaterly 9, 181–186 (1962)

15. Dinkelbach, W.: Die maximierung eines quotienten zweier linearer funktionen unter linearen nebenbedingungen. Z. Wahrscheinlichkeitstheorie 1, 141–145 (1962)

16. Dinkelbach, W.: On nonlinear fractional programming. Managment Science 13(7), 492–498 (1967)

17. Anzai, Y.: On integer fractional programming. J. Operations Research Soc. of Japan 17(1), 49–66 (1974)

18. Adams, W.P., Forrester, R.J., Glover, F.: Comparison and enhancement strategies for linearizing mixed 0-1 quadratic programs. Discrete Optimization 1, 99–120 (2004)

19. Adams, W.P., Forrester, R.J.: A simple recipe for concise mixed 0-1 linearizations. Operations Research Letters 33, 55–61 (2005)

20. Glover, F.: Improved linear integer programming formulations of nonlinear integer problems. Managment Science 22(4), 455–460 (1975)

21. Conforti, M., Cornuéjols, G., Zanbelli, G.: Integer Programming. Springer, ??? (2014)

22. Sharir, M.: A strong-connectivity algorithm and its applications to data flow analysis. Computers and Mathematics with applications 7(1), 67–72 (1981)

23. Tarjan, R.: Depth-first search and linear graph algorithms. SIAM Journal on Computing 1(2), 146–160 (1972)

24. Dijkstra, E.W.: A Discipline of Programming. Prentice-Hall, ??? (1972)

25. Berthold, T.: Primal heuristics for mixed integer programs. PhD thesis, Technische Universität Berlin (2006)

26. Glover, F., M., L.: General purpose heuristics forinteger pro-gramming - part i. Journal of Heuristics 2, 343–358 (1997)

27. Glover, F., M., L.: General purpose heuristics forinteger pro-gramming - part ii. Journal of Heuristics 3, 161–179 (1997)

28. Fischetti, M., Glover, F., A., L.: The feasibility pump. Mathematical Programming 104, 91–104 (2005)

29. Balas, E., Schmieta, S., Wallace, C.: Pivot and shift - amixed integerprogramming heuristic. Discrete Optimization 1, 3–12 (2004)

30. Balas, E., Martin, C.H.: Pivot-and-complement: A heuristic for 0-1 programming. Management science 26, 86–96 (1980)

31. Dijkstra, E.W.: A note on two problems in connexion with graphs. Numerische Mathematik 1, 269–271 (1959)

32. Johnson, D.B.: Efficient algorithms for shortest paths in sparse networks. Journal of the ACM 24(1) (1977)

33. Ahuja, R.K., Mehlhorn, K., Orlin, J., Tarjan, R.E.: Faster algorithms for the shortest path problem. Journal of the ACM 37(2) (1990)

34. Taccari, L.: Integer programming formulations for the elementary shortest path problem. European Journal of Operational Research 252(1) (2016)

35. Yuan, Y., Van Allen, E.M., Omberg, L., Wagle, N., Amin-Mansour, A., Sokolov, A., Byers, L.A., Xu, Y., Hess, K.R., Diao, L., Han, L., Huang, X., Lawrence, M.S., Weinstein, J.N., Stuart, J.M., Mills, G.B., Garraway, L.A., Margolin, A.A., Getz, G., Liang, H.: Assessing the clinical utility of cancer genomic and proteomic data across tumor types. Nat. Biotechnol. 32(7), 644–652 (2014)

36. Subramanian, A., Tamayo, P., Mootha, V.K., Mukherjee, S., Ebert, B.L., Gillette, M.A., Paulovich, A., Pomeroy, S.L., Golub, T.R., Lander, E.S., Mesirov, J.P.: Gene set enrichment analysis: a knowledge-based approach for interpreting genome-wide expression profiles. Proc. Natl. Acad. Sci. U.S.A. 102(43), 15545–15550 (2005)

37. Foroutan, M., Bhuva, D.D., Lyu, R., Horan, K., Cursons, J., Davis, M.J.: Single sample scoring of molecular phenotypes. BMC Bioinformatics 19(1), 404 (2018)

